# A Subcutaneous Implant of Tenofovir Alafenamide Fumarate Causes Local Inflammation and Tissue Necrosis in Rabbits and Macaques

**DOI:** 10.1101/775452

**Authors:** Jonathan T. Su, Solange M Simpson, Samuel Sung, Ewa Bryndza Tfaily, Ronald Veazey, Mark Marzinke, Jiang Qiu, David Watrous, Lakmini Widanapathirana, Elizabeth Pearson, M. Melissa Peet, Dipu Karunakaran, Brooke Grasperge, Georgina Dobek, Charlette M. Cain, Thomas Hope, Patrick F. Kiser

## Abstract

We describe the *in vitro* and *in vivo* evaluation of a subcutaneous reservoir implant delivering tenofovir alafenamide hemifumarate (TAF) for the prevention of HIV infection. These long-acting reservoir implants were able to deliver antiretroviral drug for over 90 days *in vitro* and *in vivo*. We evaluated the implants for implantation site histopathology and pharmacokinetics in plasma and tissues for up to 12 weeks in New Zealand White rabbits and rhesus macaque models. A dose-ranging study in rabbits demonstrated dose-dependent pharmacokinetics and local inflammation up to severe necrosis around the active implants. The matched placebos showed normal wound healing and fibrous tissue encapsulation of the implant. We designed a second implant with a lower release rate and flux of TAF and achieved a median cellular level of tenofovir diphosphate of 42 fmol per 10^6^ rhesus macaque peripheral blood mononuclear cells at a dose of 10 µg/kg/day. This dose and flux of TAF also resulted in adverse local inflammation and necrosis near the implant in rhesus macaques. Inflammation in the primates was markedly lower in the placebo group than the active implant. The histological inflammatory response to the TAF implant at 4 and 12 weeks in primates was graded as a severe reaction. Thus, while we were able to achieve sustained target dose we observed unacceptable inflammatory response locally at the implant tissue interface.

## INTRODUCTION

Clinical availability of antiretroviral (ARV) delivery systems that provide durable protection from HIV transmission could revolutionize the way we fight the global HIV/AIDS pandemic. (1) Once daily oral Truvada™ (emtricitabine (FTC) 200 mg/tenofovir disoproxil fumarate (TDF) 300 mg) prevents the sexual transmission of HIV when used before sexual exposure to HIV (2–6). However, not all pre-exposure prophylaxis (PrEP) trials with oral Truvada™ have been efficacious (2–6), likely because of poor adherence to the regimen (7, 8). Therefore, many groups in the HIV prevention field are striving to develop long-acting, acceptable, and effective methods of HIV prevention for use in high-risk populations. For example, a recent clinical study demonstrated the effective drug levels of a subdermal drug-eluting implant releasing islatravir (MK-8591) for prevention of sexual transmission of HIV(9).

Long-acting drug-delivery systems are fundamentally easier for individuals to use than once-daily oral pills. Studies on adherence to methods of contraception generally show that increased duration and subsequent reduced need for daily-repeated action by the user is correlated with increased contraceptive efficacy (10–13). Subcutaneous contraceptive implants generate durable, sustained progestin exposure over several years and allow the recipient to undergo a minimally invasive procedure for implant placement without any further clinical follow up until removal. Therefore, long-acting implants are the most effective contraceptive formulations. Similarly, long-acting delivery systems of ARVs have the potential to increase adherence, providing the durable drug concentrations required to prevent HIV infection, while users are sexually exposed to the virus.

Long-acting drug delivery systems require the most potent and slowly eliminated ARVs to enable durable ARV exposure and protection for durations on the order of months to a year. Tenofovir alafenamide hemifumarate (TAF, or GS-7340) is a caspase-activated prodrug of tenofovir (TFV) (14). Based on *in vitro* analysis, the 50% effective concentration [EC_50_] of TAF is in the low nanomolar range (5 – 11.2 nM) (15, 16). TAF is intracellularly converted by kinases into TFV diphosphate (TFV-DP); this is the active form of the drug that competitively inhibits HIV reverse transcriptase and the generation of viral transcripts. The diphosphate, due to its charge and pK_a_, is highly impermeable to cellular membranes and is therefore trapped in the intracellular volume versus the parent species. TFV-DP cellular half-lives in humans have been measured at ∼150 hours (17). For several other molecular reasons that have been reviewed (18, 19), TAF is a more potent pro-drug than TDF, resulting in lower TFV drug exposures and reduced side effects as compared to TDF (14). Together, these characteristics make TAF one of the leading drug molecules for long-acting ARV delivery because TAF is so potent and cellularly long-acting that one can plausibly load many ARV daily doses inside a small controlled-release device to achieve durable protection from HIV infection.

Accordingly, four subcutaneous implants delivering TAF exist in the literature. The first implant, presented by Gunawardana and Baum, delivered 0.92 mg/day (∼80 µg/kg/day) and was evaluated in beagle dogs for over 40 days (20). This implant consisted of 1.9 mm silicone tubing with 14 poly(vinyl alcohol) coated 1.0 mm diameter delivery channels punched into the walls and filled with pure TAF powder (20). The second implant was presented by Schlesinger and Desai, and consisted of a heat-sealed poly(caprolactone) film cylinder containing TAF and polyethylene glycol 300 at 1:2, 1:1, or 2:1 w/w ratios (21). Release ranging from 0.5 – 4.4 mg/day was demonstrated *in vitro* (21). A third implant was presented by Johnson *et al,* and was a reservoir formed from extruded poly(caprolactone) filled with TAF and castor oil excipient; release rates of 0.15 – 0.91 mg/day were demonstrated *in vitro* (22). The fourth implant, presented by Chua and Gratonni, consisted of a refillable titanium device that delivered TAF and FTC through silicon nanochannels (23). This refillable implant demonstrated the sustained release of TAF of ∼ 0.2 mg/day (∼ 20 µg/kg/day) for 83 days in rhesus macaques (23). They rapidly achieved TFV-DP benchmark levels in macaque peripheral blood mononuclear cells (PBMCs) with means of 72 fmol/10^6^ PBMCs early in the pharmacokinetic (PK) study to 533 fmol/10^6^ PBMCs to day 70. None of the studies have reported placebo-controlled histopathology, but all authors have suggested that the implants are safe. (20, 23, 24)

When a subcutaneous implant is placed under the skin, the cells and tissues surrounding the implant respond to the presence of the foreign-body implant and potentially to the drug near the implantation site. In the normal foreign body response, the implant is walled off at its site of implantation by a fibrin-containing capsule (25). Focal toxicity can result in inflammation and necrosis, potentially leading to skin disruption and potential infection (23, 26). Ultimately, any process that disrupts the multistage wound healing response can result in a non-biocompatible drug delivery implant. It is also possible that these toxic effects are dependent on the route of administration, thus driving the need for careful evaluation of the biological and cellular response at the implant-tissue interface.

We evaluated the potential viability of TAF for systemic, long-term drug delivery, using a subcutaneous implant made of a heat-sealed polyurethane rate-controlling membrane. Our objective was to develop a subcutaneous TAF reservoir implant that, after implantation, would show no signs of pathology at the implant site, yet provide levels of TFV-DP that could prevent sexual transmission of HIV. (26) In this work, we evaluated a TAF implant in both New Zealand White (NZW) rabbit and rhesus macaque models. We describe conducted studies below to assess the local biological reaction to active TAF implants versus matched placebo implants, TFV-DP pharmacokinetics and *in vivo* release rates in both NZW rabbits and rhesus macaques for up to 12 weeks.

## RESULTS

### Implant design and in vitro performance

TAF reservoir implants were formed by compressing the TAF drug substance and small amounts of NaCl and magnesium stearate into a pellet that was impulse-sealed into a 150 to 170 µm thin, medical-grade polyurethane tube (**Figure 1**). The tube wall acts as a mechanical capsule and rate controlling membrane whose composition or thickness can be changed to tune the drug release rate. Accordingly, we were able to successfully control the release of TAF by changing the geometry of the implant, as well as the composition of the polyurethane membrane. Drug release from implants was evaluated *in vitro* (**Figure 2**) using the shake flask method. We observed slow rates of TAF degradation (27) in phosphate-buffered saline (PBS) in *in vitro* release testing media (half-life of ∼48 hours at pH 7.4 at 37 °C in PBS), as well as in the implant (see **Suppl. 1**). Although until the end of the release curve we observed that greater than 90% of the internal contents was the parent form. Molar amounts of TAF and its two main TAF related substances (monophenyl PMPA and PMPA monoamidate) were calculated and converted to a mass of TAF equivalents released. Consequently, in this work, TAF equivalents were used in all of our calculations of drug release. Two generations of implants are described in this study: Generation A and Generation B.

**Figure 1.**
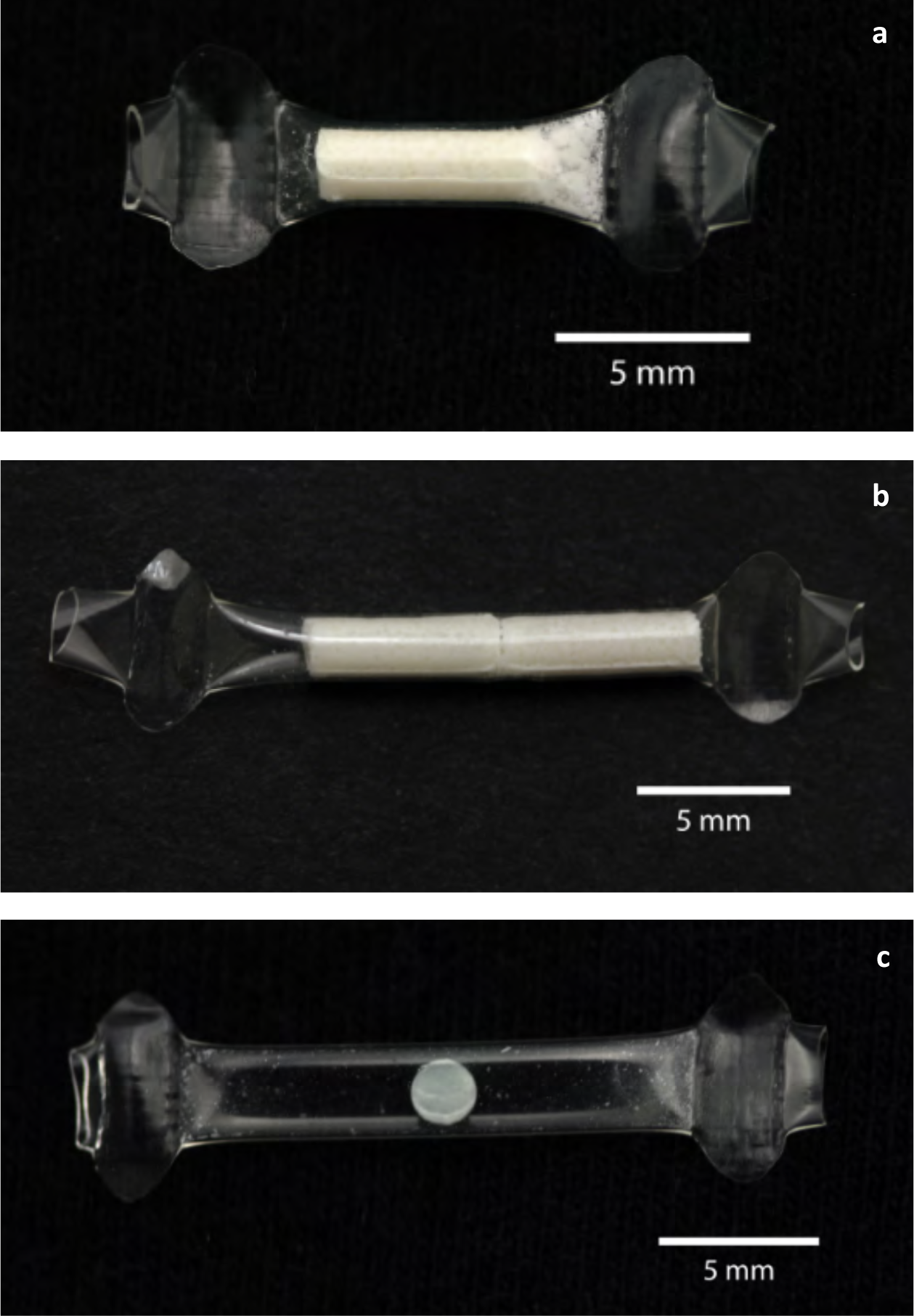
Photos of representative Generation A TAF long-acting reservoir implants with lumen lengths of (**a**) 0.8 cm and (**b**) 1.6 cm and (**c**) a placebo implant that is empty except for a pellet of NaCl and magnesium stearate.

**Figure 2.**
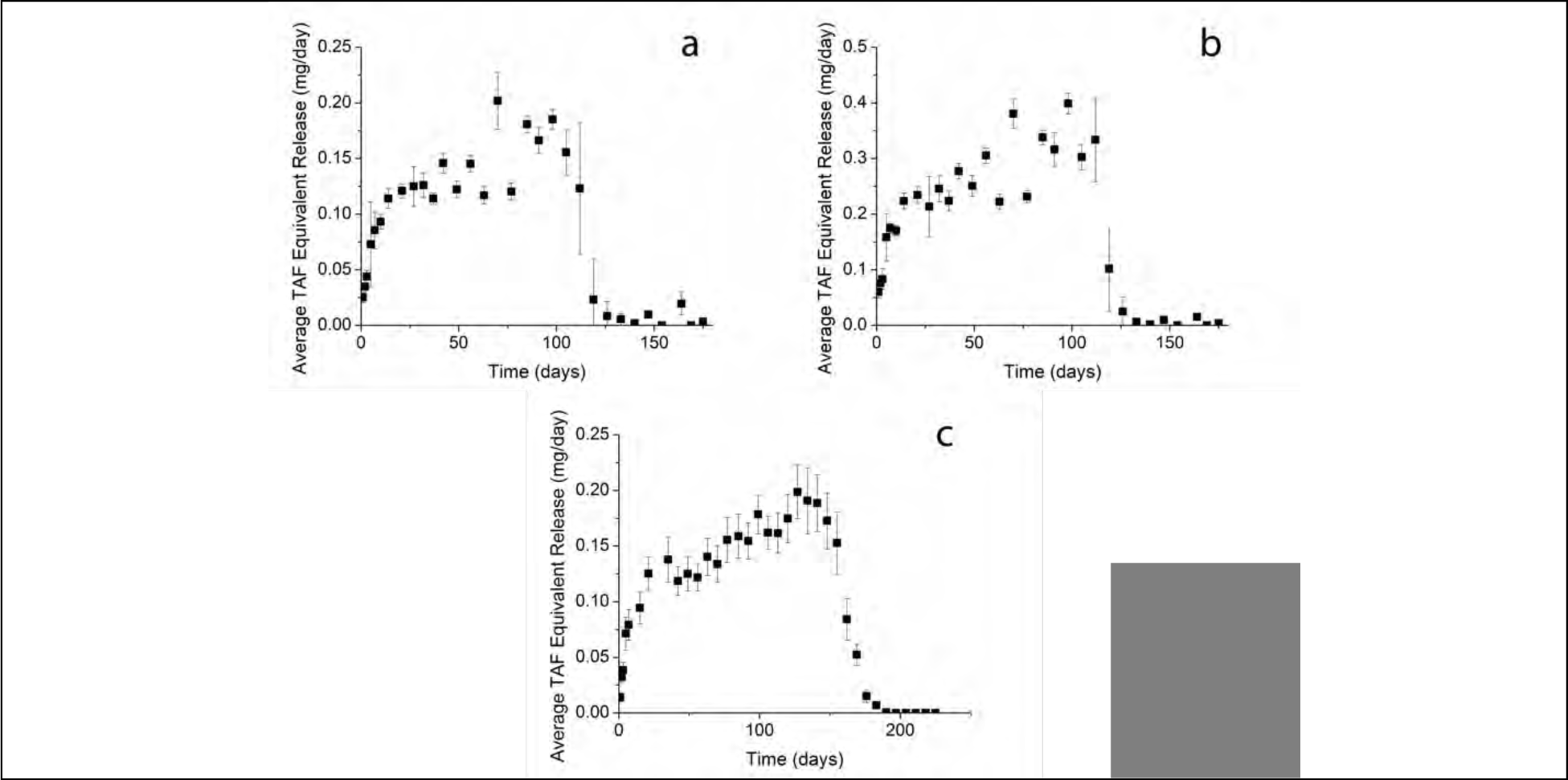
(**a)** TAF Gen A 0.8 cm lumen length implant *in vitro* release (n=10) error bars ± SD. (**b)** TAF Gen A 1.6 cm lumen length implant *in vitro* release (n=9) error bars ± SD. (**c)** TAF Gen B implant *in vitro* release (n=10) error bars ± SD.

Two types of Generation A implants were created one with a 0.8 cm lumen length, and one with a 1.6 cm lumen length. The average TAF equivalent *in vitro* release rate over days 7 to 91 from the Generation A implant was 0.13 mg/day for the 0.8 cm long implants, and 0.26 mg/day for the 1.6 cm long implants with a flux of 0.24 mg TAF/cm^2^/day and 0.23 mg TAF/ cm^2^/day, respectively. We were able to obtain a sustained release of the drug *in vitro* (**Figure 2a, b**) for over one hundred days. Guided by our pharmacokinetics and histopathology from studies on the Generation A implants, we developed a second-generation implant, called Generation B that was designed to release a lower amount and flux of drug, but also provide a steady release of TAF *in vitro* (**Figure 2c**). The mean TAF equivalent *in vitro* release rate over days 7 to 91 from the Generation B implants was 0.13 mg/day, with an average flux of 0.08 mg TAF/cm^2^/day.

### Pharmacokinetic and local safety evaluation in rabbits

Both generations of TAF implants were evaluated in animals. Four studies were performed: (1) a PK and safety dose-ranging study using Generation A implants in NZW rabbits using four dose groups, (2) a PK study using Generation A implants in rhesus macaques, (3) a PK and local response study using Generation B implants in rhesus macaques and (4) an exploratory study in rhesus to assess the local reaction to Generation B implants when inserted by trocar. **Table 1** summarizes the results from the first three animal studies in this series of analogous implants delivering TAF.

**Table 1:**
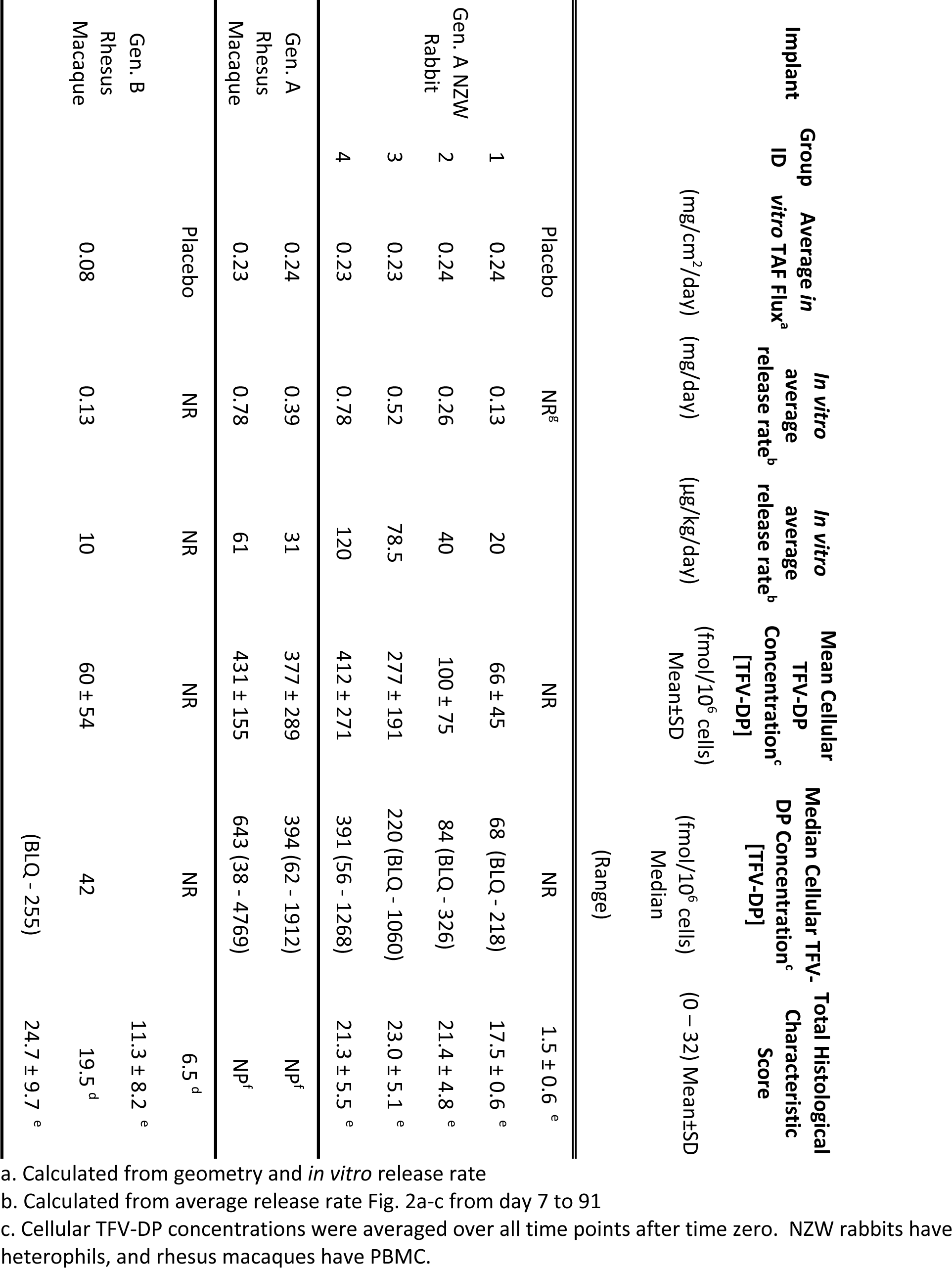
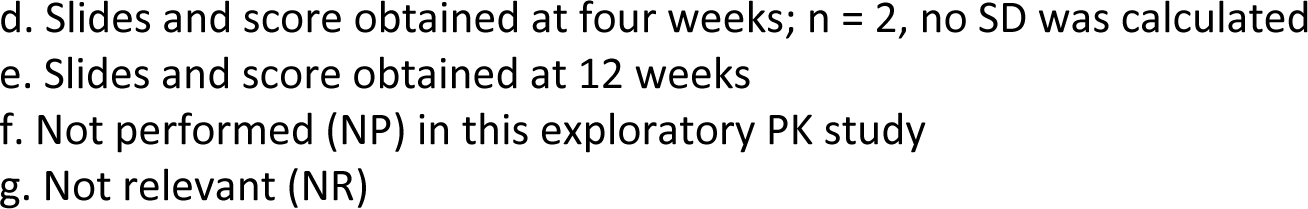
Summary of PK and pathohistological scores in NZW rabbits and rhesus macaques.

A series of four doses was evaluated in NZW rabbits (**Table 1**). TFV-DP was found in blood heterophils for all active implant treated animals after week 1, and concentrations were quantifiable throughout the study (**Figure 4**). In general, increases in the in *vitro* dose were correlated with increasing median TFV-DP (**Figure 3**). Group 1, our lowest *in vitro* dose, 0.13 mg/day, corresponded to a median TFV-DP level of 68 fmol/10^6^ cells (range from below the level of quantification (BLQ) to 218 fmol/10^6^ cells); the median was calculated from weeks 1 – 12 cellular PK data. Group 4, our highest *in vitro* dose of 0.78 mg/day—5.5 times higher than the lowest dose—provided a median TFV-DP level of nearly 391 fmol/10^6^ cells doses over weeks 1 – 12 of the study. This implant gave a median cellular TFV-DP level 5.75 times higher than the lower dose.

**Figure 3.**
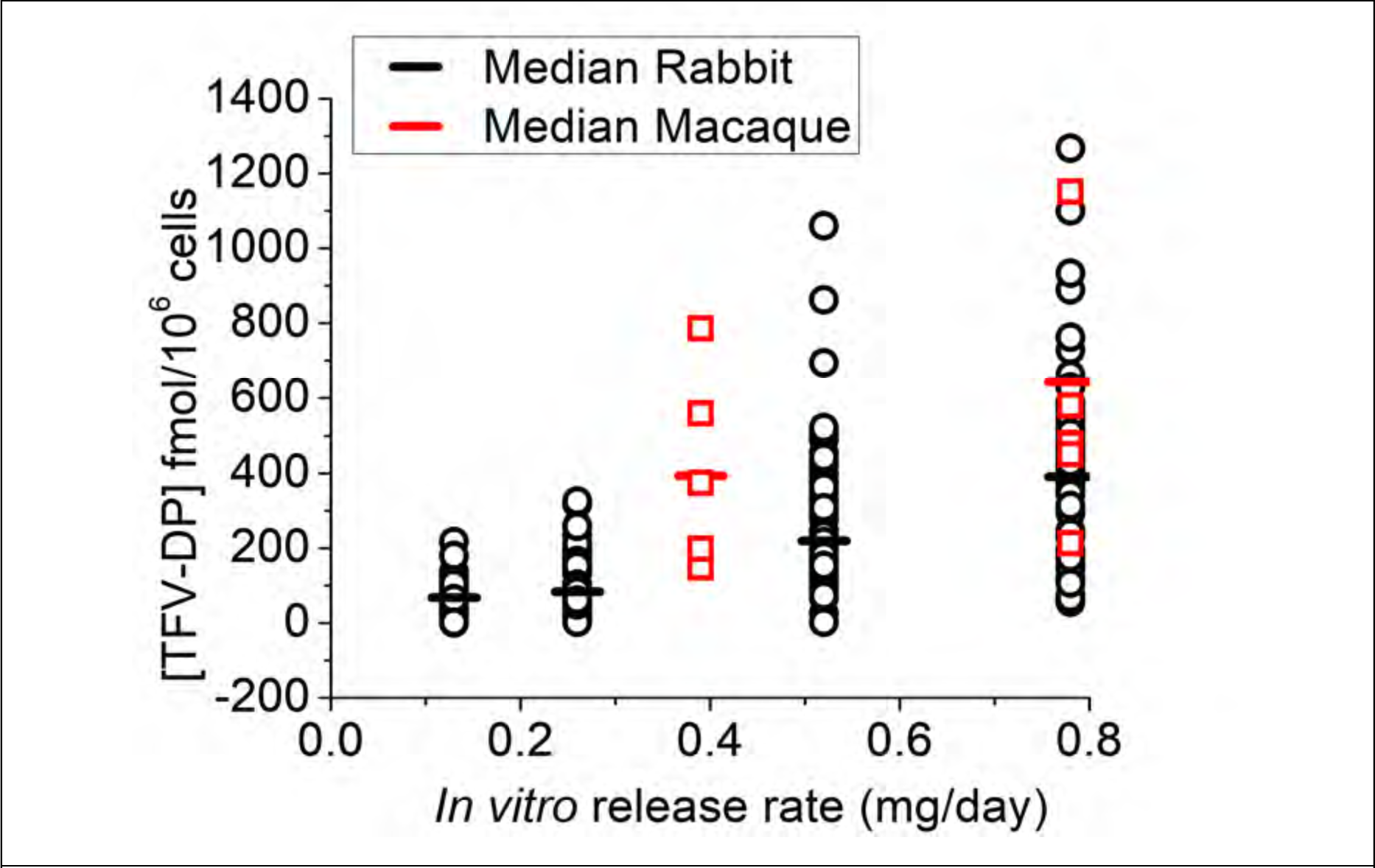
TFV-DP concentrations were determined in NZW rabbits and macaques for Generation A TAF implants. The median was determined from data points from weeks 1 – 12.

**Figure 4.**
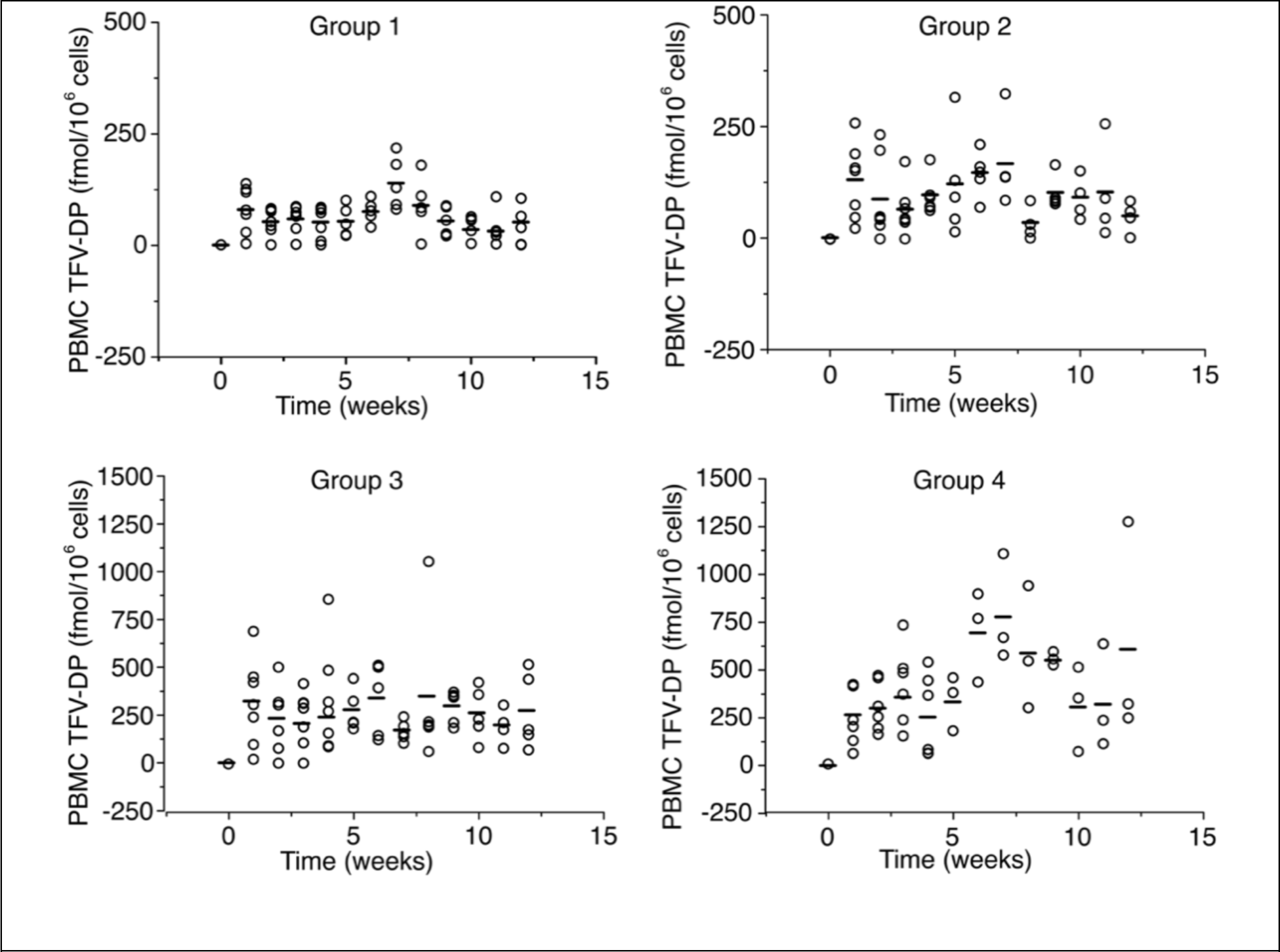
Dose-dependent PK was observed in our NZW rabbit experiments, with higher average TFV-DP levels leading to higher plasma levels of TFV-DP in circulating heterophils. TFV-DP levels increased within a week of implantation. BLQ values are plotted as 1/10th of the calculated LOQ value. Drug levels for all placebo implants were BLQ. The horizontal bar is mean TFV-DP. As per Table 1*, in vitro* release from Group 1: 0.13 mg/day; Group 2: 0.26 mg/day; Group 3: 0.52 mg/day, Group 4: 0.78 mg/day.

Plasma TFV concentrations remained low for all NZW rabbits, with TFV concentrations ranging from BLQ to 20 ng/mL (LLOQ = 0.31 ng/mL) (weeks 1 - 12; see **Suppl. 2, Table S2**). Similarly, drug and metabolite concentrations in vaginal and rectal tissue were generally low in NZW rabbits (Full data are contained in **Suppl. 2, Table S1**). Values of TFV-DP in vaginal tissues and rectal tissues ranged from BLQ to 169 fmol/mg and from BLQ to 50 fmol/mg (LLOQ = 50 fmol/sample; weeks 1 - 12), respectively. Samples were taken near the implant site at necropsy, and the TFV-DP levels were scattered; large concentrations of TFV-DP were found at week 12. Local tissue TFV-DP concentrations ranged from 0.86 to 69,941 fmol/mg of TFV-DP.

All NZW rabbits appeared healthy, with no superficial observations of poor tolerability at the implant site throughout the study. **Figure 5** shows representative sections of histology from NZW rabbits with Generation A implants after 12 weeks. In the placebo implants (**Figure 5a** and **5b**), the implant region was demarcated by a thin fibrous tissue capsule, two to five cells thick, but most of the sections showed no or mild inflammation in the area around the implant, with two implants showing mild to moderate inflammation. We saw that some rabbits appeared to have inflammation near the end of the implant adjacent to the sutures. The active implants are shown in **figure 5c**, and **5d** displayed severe granulomatous and suppurative inflammation with necrosis and abundant necrotic cell debris and proteinaceous fluid in the implant space, which is lined by marked infiltrations of lymphocytes and heterophils. Marked infiltrations of lymphocytes and macrophages into the adjacent muscle tissues were seen. There was also abundant eosinophilic fluid-like material with pockets of necrotic cellular debris with associated granulomatous inflammation and giant cells. Some slides showed scattered yet diffuse infiltrations of lymphocytes, heterophils, and macrophages in the dermis and muscle fibers. Additionally, in some NZW rabbits, chronic granulomatous inflammation and necrosis were observed around the implant.

**Figure 5.**
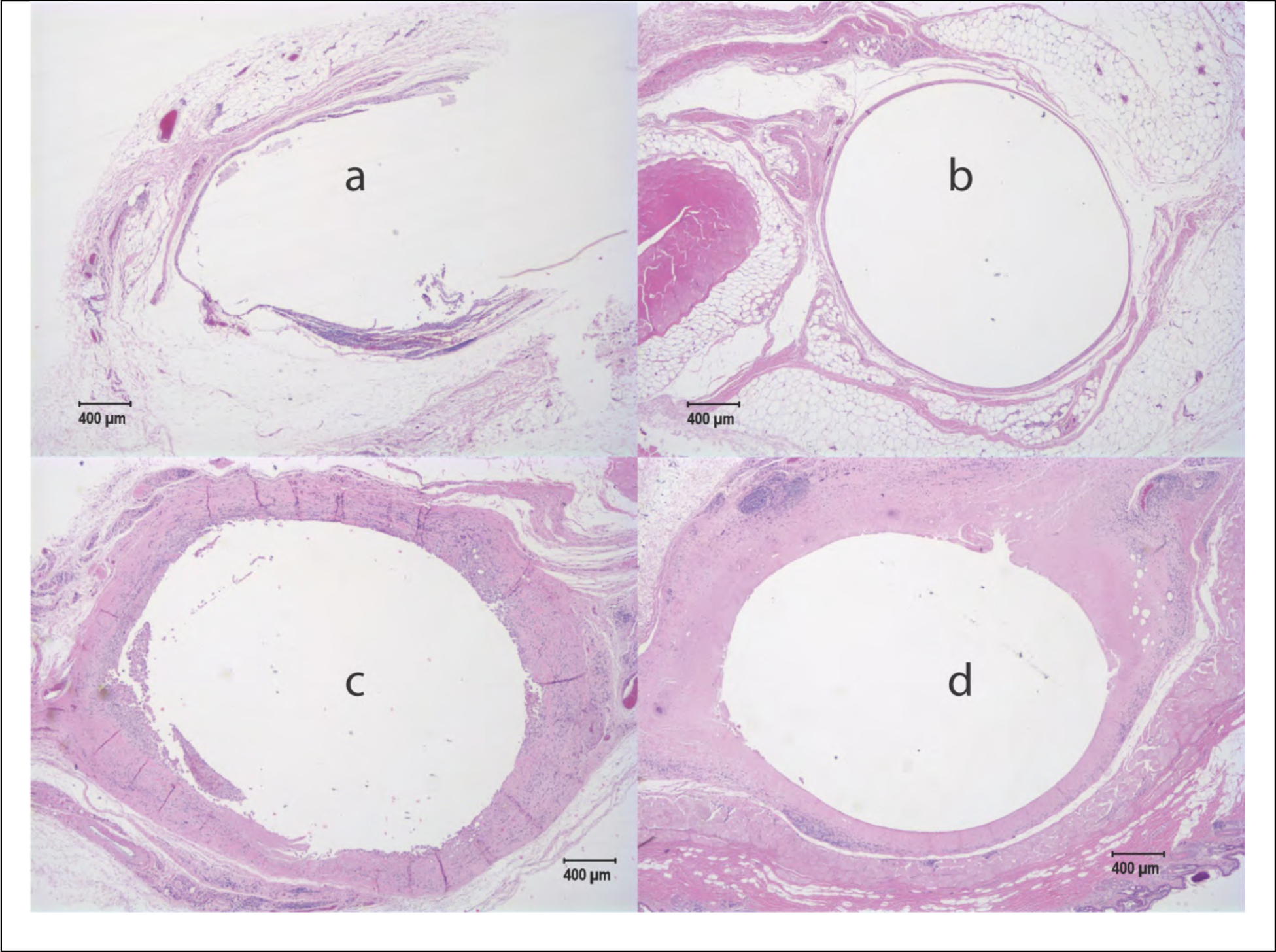
Contralateral sections from NZW rabbits after 12 weeks with Generation A implant. Sections from two animals (**a, c**: M29051; **b, d**: M29049) are shown. Minimal inflammation was observed for our placebo implants (**a, b**), however extensive inflammation with some necrosis was observed for implants containing active drug (**c, d**).

Histopathological characteristics were scored semi-quantitatively on an animal and implant basis from multiple slides taken from the ends and center of fixed tissue containing implants. The scoring system (0-4) logged the presence of five cellular characteristics (polymorphonuclear cells, lymphocytes, plasma cells, macrophages, giant cells) and three tissue characteristics (necrosis, capsule thickness, and tissue infiltrate). High levels of inflammation were observed in the peri-implant space from all drug-loaded implants at all doses in NZW rabbits (ranging from an average total histological characteristic score of 17.5 ± 0.6 to 23.0 ± 5.1) (±SD, averaged over all time points weeks 1 - 12, **Table 1**). These total histological characteristic scores were far higher than those observed for the placebo implants, which had an average score of 1.5 ± 0.6 (±SD, averaged over all time points weeks 1 – 12, **Table 1**) (see scores tables and micrographs from all animals in **Suppl. 3**). There was no statistically significant difference (p > 0.05) in total histological characteristic score with TAF exposed NZW rabbit groups. The 0.8 and 1.6 cm Generation A implants obtained a reactivity grade of *severe reaction* (see Methods and **Suppl. 5**).

After our *in vivo* experiments, implants were placed on *in vitro* release to verify the implants were intact, not leaking, and were releasing drug at the moment of removal (**Figure 6**). Release after explantation from the 1.6 cm lumen length implants was roughly double that of the 0.8 cm lumen length implants, as expected. Additionally, we extracted these implants for the calculation of an average *in vivo* release rate (**Figure 7**). Comparison of our *in vivo* release with our *in vitro* release demonstrated a correlation of close to 0.8 at four weeks and approximately 0.7 - 0.8 at 12 weeks (**Figure 7**). All returned implants remained intact, and the wall was not compromised. We investigated if any molecular weight changes had occurred in the polymers during their residence *in vivo*. No notable changes in molecular weight distributions were detected that would indicate *in vivo* polymer degradation (**Suppl. 6, Table S15**).

**Figure 6.**
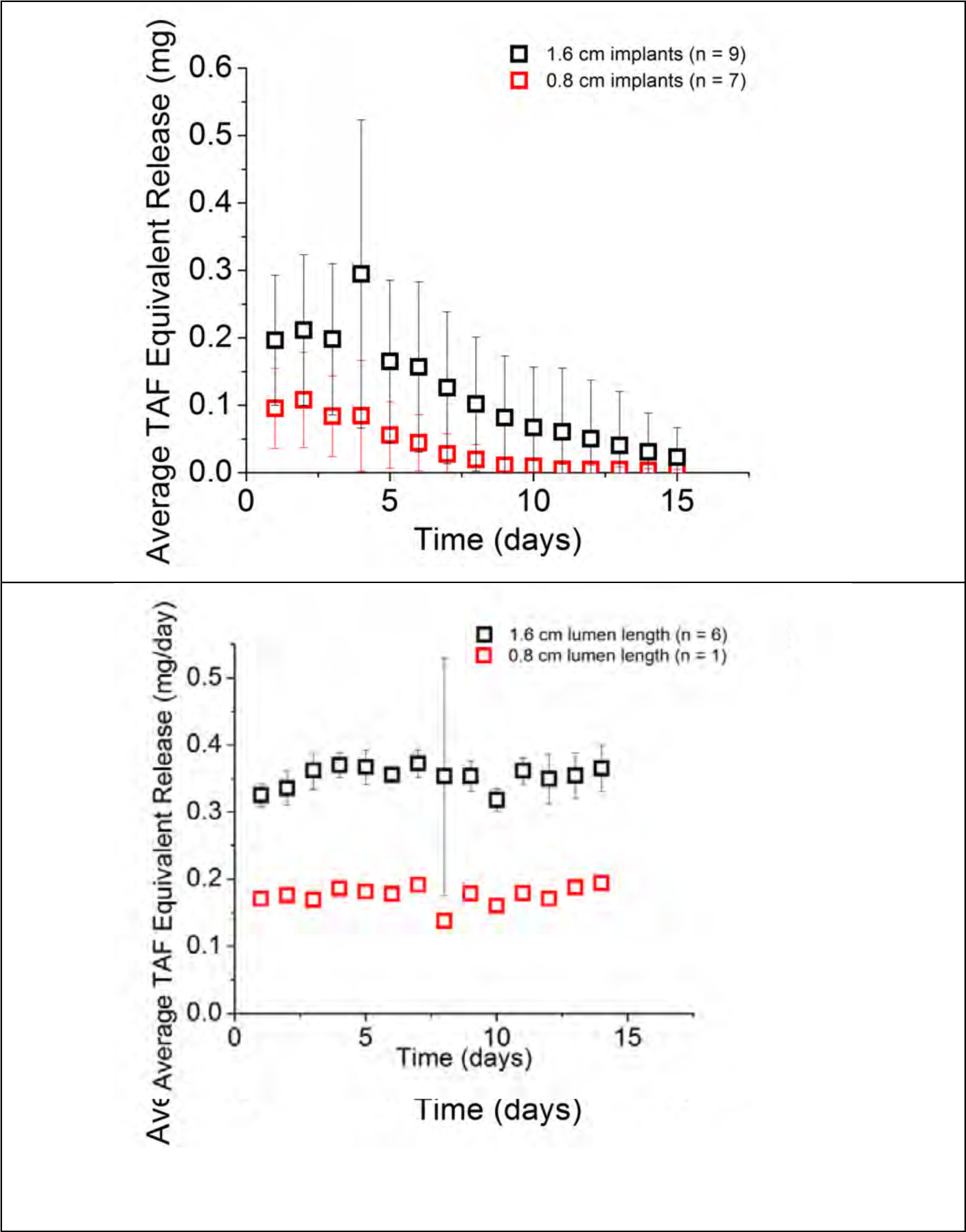
Generation A Implants not selected for histology were removed intact from NZW rabbits (top) at 12 weeks and rhesus macaques (bottom) at 12 weeks and placed on *in vitro* release to verify that expected release was being achieved.

**Figure 7.**
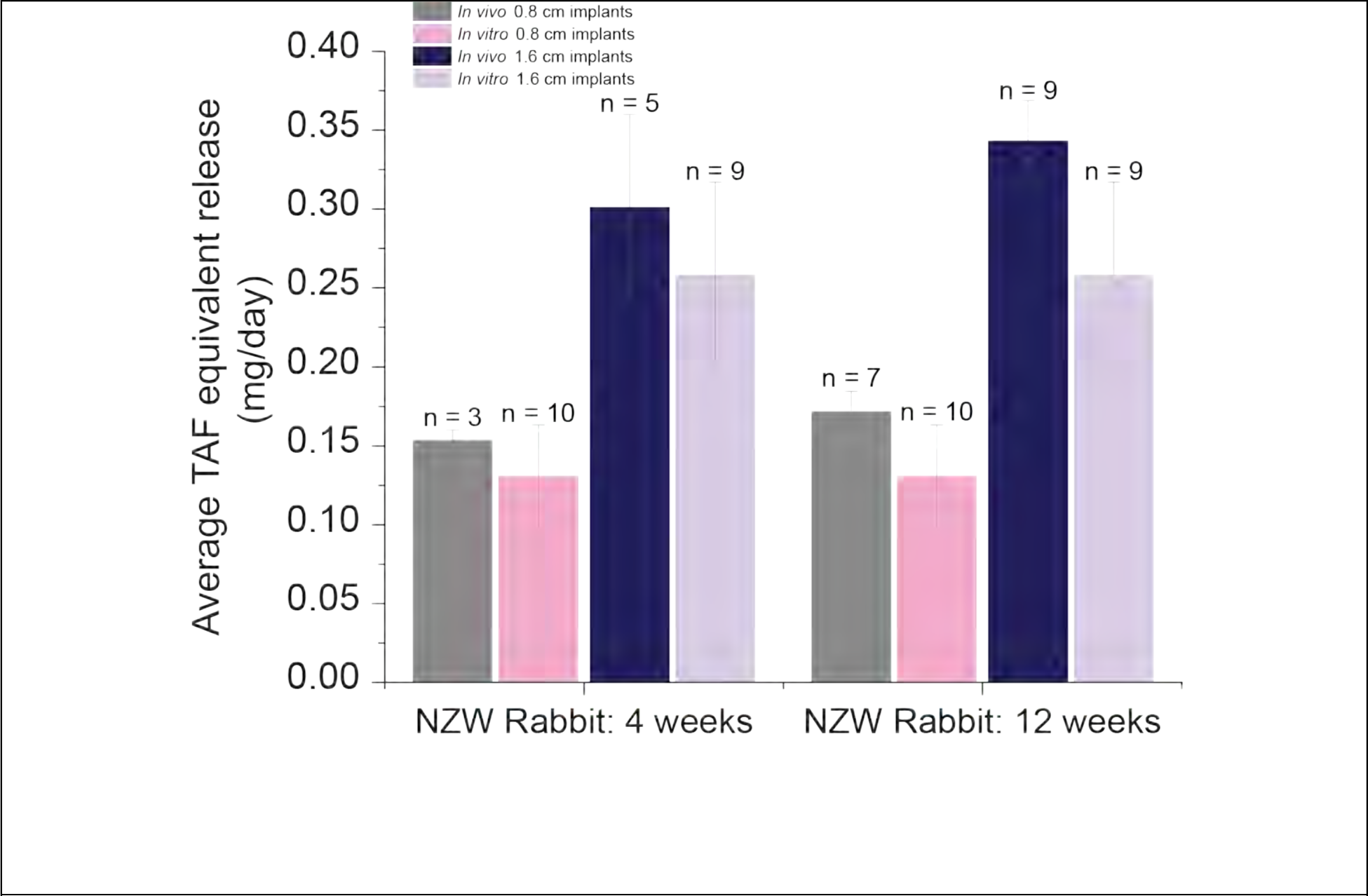
Comparison of average daily *in vitro* release over 7 to 91 days and the estimated *in vivo* release in NZW rabbits received at four weeks (left) and 12 weeks (right) from TAF Generation A implants.

### Pharmacokinetic and local safety evaluation in rhesus macaques

Following experiments with Generation A implants in NZW rabbits, we conducted a dose-finding pharmacokinetic experiment in rhesus macaques with the same implant system (See **Table 1**). In Generation A implanted macaques, there was no statistically significant difference found (P > 0.05) between TFV-DP levels for either our 0.39 mg/day *in vitro* dose or our 0.78 mg/day *in vitro* dose (**Figure 3**). This study allowed us to design a lower dose and flux Generation B implant to achieve lower levels of TFV-DP. In the low and high dose one animal each lost implants due to abscess formation. Comparison of our *in vivo* release with our *in vitro* release demonstrated a close correlation (**Figure 8**).

**Figure 8.**
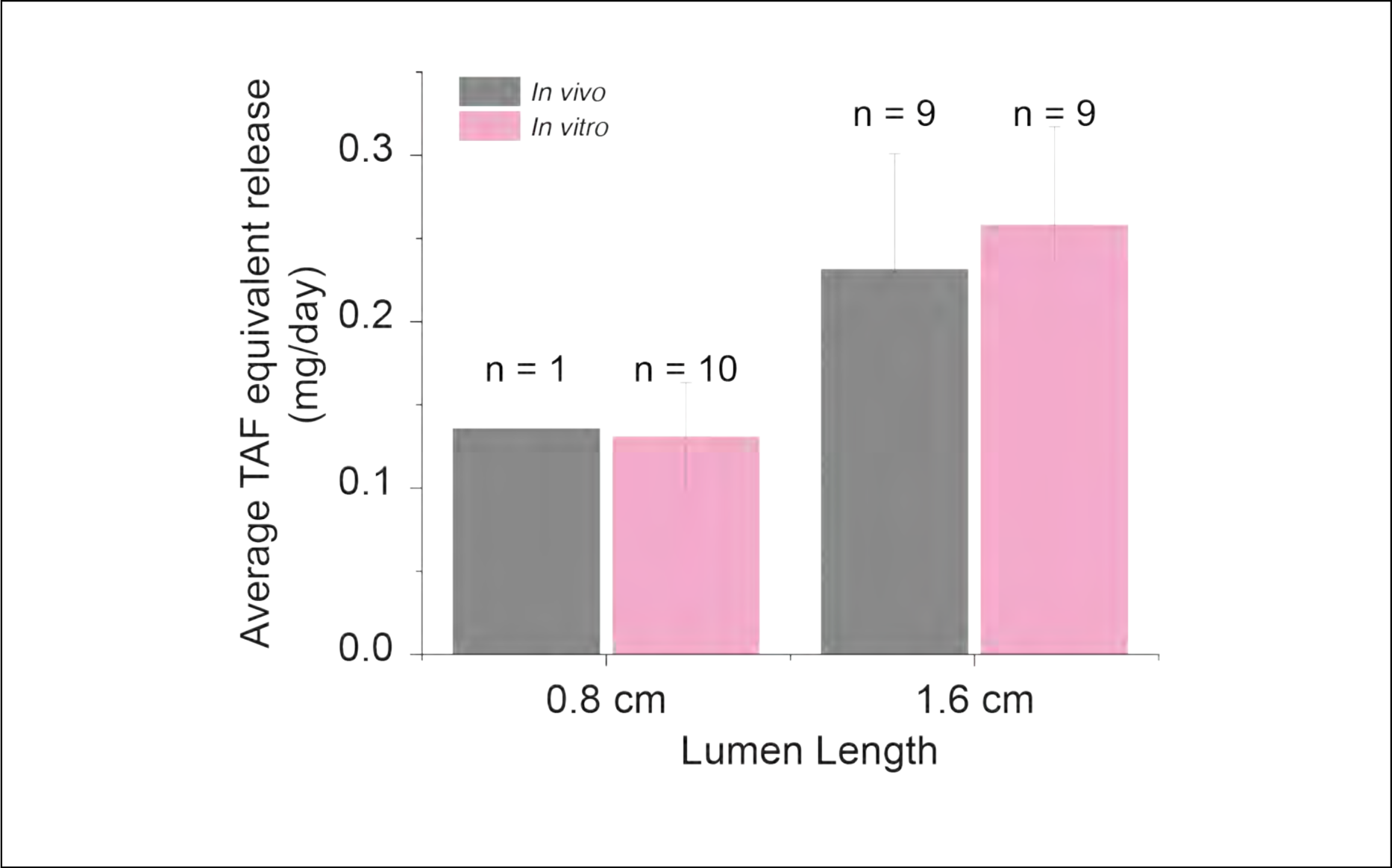
*In vitro* and *in vivo* comparison average daily release from TAF Generation A implants in rhesus macaques after 12 weeks. The *in vitro* release is the average over days 7 to 91 from representative implants in the same batch as the *in vivo* implants.

For Generation B implants releasing an average of 0.13 mg TAF/day *in vitro*, the median TFV-DP concentrations in rhesus macaque PBMCs were 42 fmol/10^6^ cells (range: BLQ-255), calculated from weeks 1 – 12 data (**Figure 9, and Table 1**). Plasma TFV and TAF concentrations remained low for all rhesus macaques, with TFV concentrations ranging from BLQ to 7 ng/mL (LLOQ = 0.35 ng/mL) and TAF concentrations ranging from BLQ to 4 ng/mL (TAF LLOQ = 0.03 ng/mL) (see **Suppl. 7, Table S18**). Similarly, TFV and TFV-DP levels in tissue were generally low for all macaques, with full data provided in **Suppl. 7, Table S17**. Samples were taken near the implant site, the vagina, and the rectum. All TFV concentrations near the implant site were BLQ (LLOQ = 0.05 ng/sample), and TFV-DP concentrations near the implant site were low, ranging over BLQ – 27 fmol/mg (LLOQ = 5 fmol/sample). Rectal concentrations were similarly low, with a TFV range from BLQ to 0.24 ng/mg; TFV-DP ranged from BLQ to 12 fmol/mg. In the vagina, concentrations were again low: TFV ranged from BLQ to 0.02 ng/mg, while TFV-DP ranged from BLQ to 6 fmol/mg.

**Figure 9.**
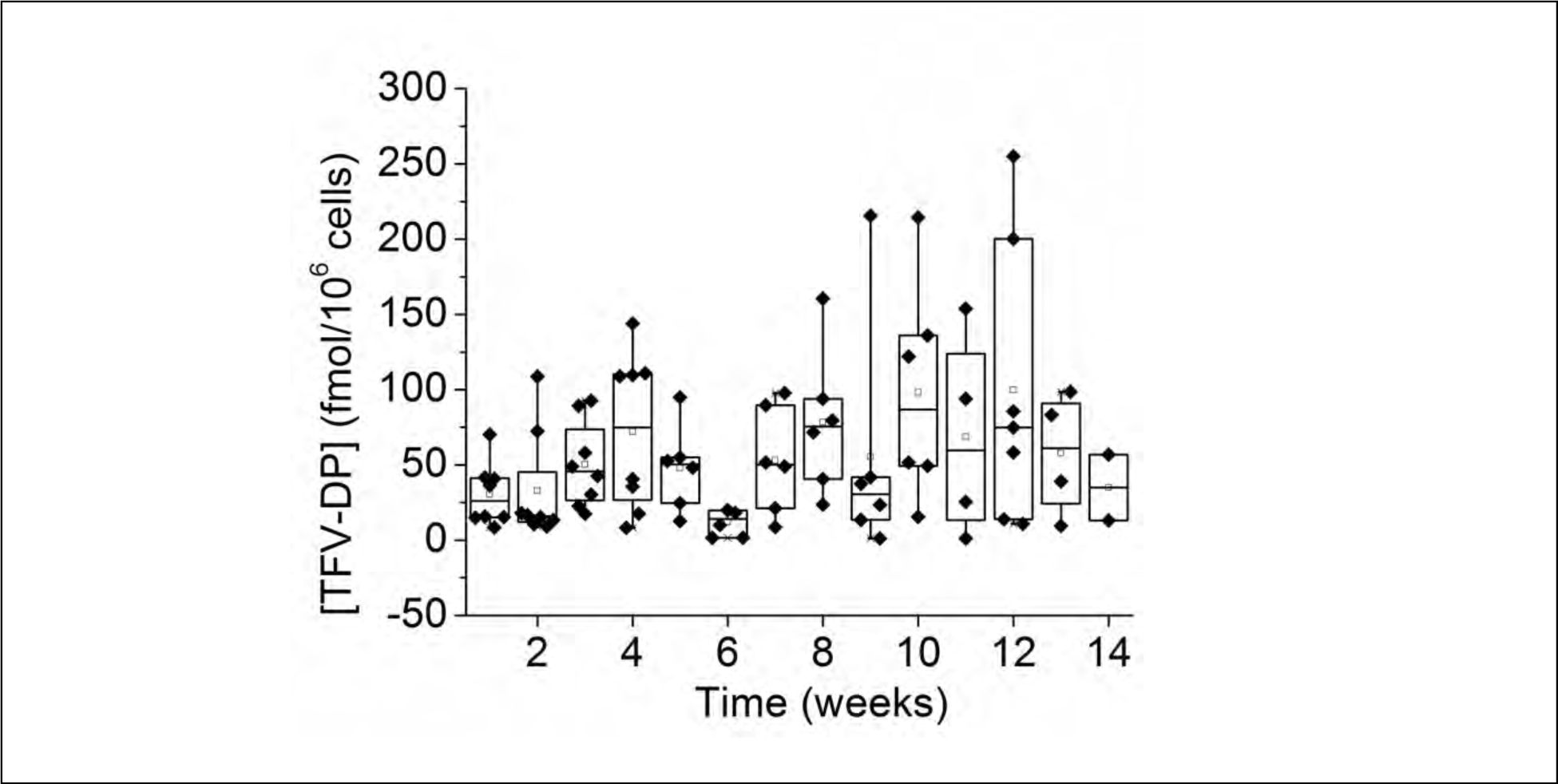
TFV-DP levels in macaque PBMCs from the Generation B implants.

Despite the lower flux of TAF release in the Generation B implants, inflammation remained high in all rhesus macaques exposed to the active implants. For the active Generation B implants in macaques, we observed gross redness around the implant and neovascularization indicative of inflammation. All placebos showed visually clear encapsulation with no neovascularization and redness around the placebo implant. Representative pathohistology for the Generation B implants in macaques are shown in **Figure 10**, with the full set of images in **Suppl. 4** (**Figures S25 – S30**). Histologically we observed moderate to severe inflammation in the peri-implant space, with thick fibrous capsules filled with neutrophils, plasma cells, necrotic cellular debris, proteinaceous fluid, and occasional multinucleated giant cells. Multifocal aggregates of densely packed lymphocytes were observed in surrounding tissues. In two of the four macaques, we observed an abscess above the implant site where it appeared that the implant caused a topical wound. The two animals that were necropsied at four weeks also displayed markedly more inflammation than the placebo contralateral implants, but the necrosis scores in the active implant sites were mild at four weeks (**Suppl. 4, Tables S8-S9**) and became moderately to severely necrotic by 12 weeks (**Suppl. 4, Tables S10**-**S13**).

**Figure 10.**
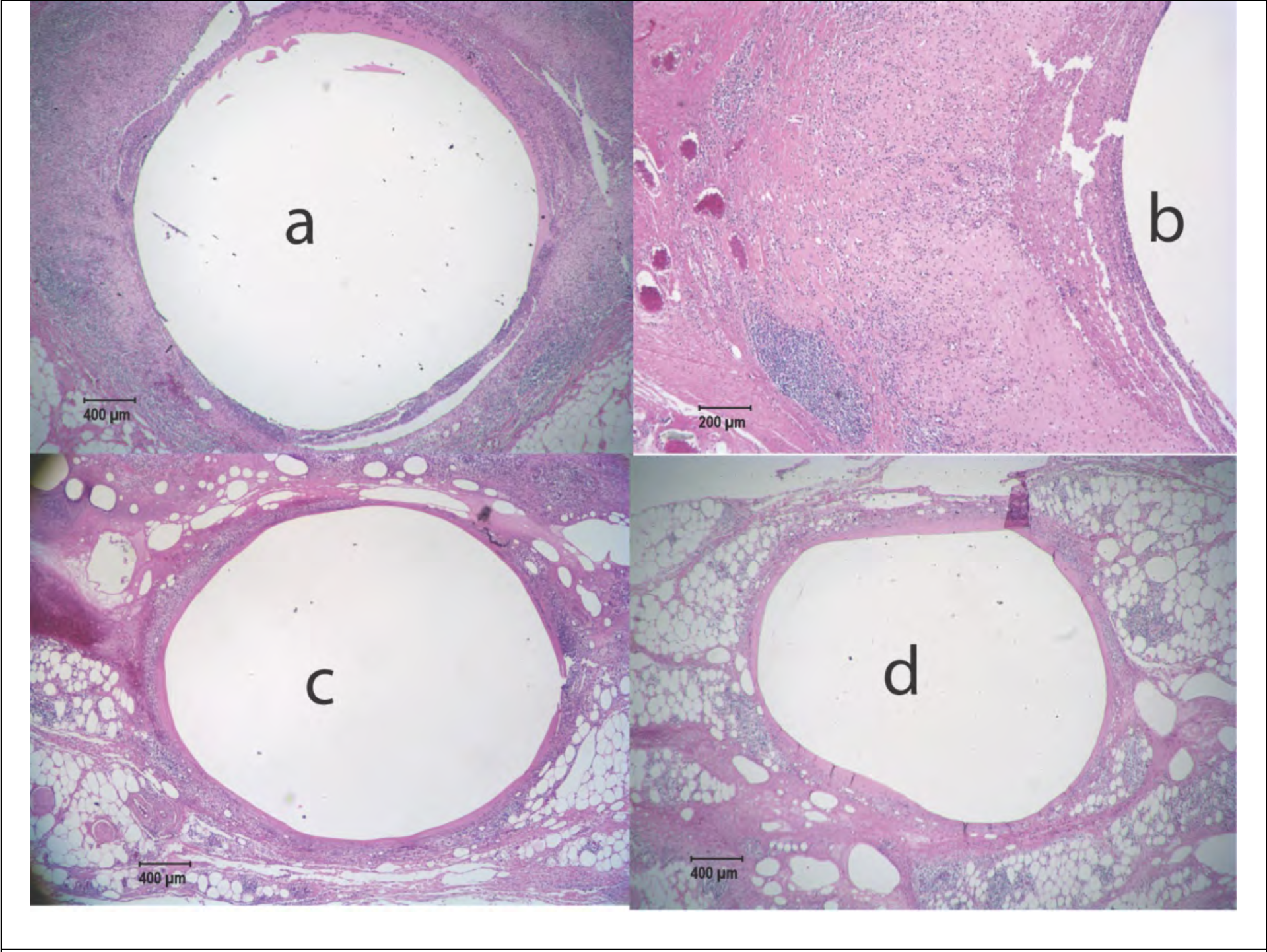
Representative histology slides from rhesus macaque FC48 after 12 weeks with a Generation B implant. Minimal inflammation and a distinct fibrous tissue capsule was observed for the placebo implant (**c,d**), however extensive inflammation was observed for implant containing active drug (**a,b**) despite a lower *in vitro* release rate of Generation B implants (0.13 mg/day *in vitro* release rate) versus the Generation A implants.

In contrast, two out of four placebo implants showed minimal fibrosis or inflammation, with a few neutrophils and plasma cells in the implant lumen; except in one case, where marked accumulations of macrophages and lymphocytes with moderate numbers of giant cells were observed around the placebo implant. In this animal (EC74, see **Table S11**), moderate inflammation was observed around the placebo implant adjacent to the sutures, but in this animal, the active paired implant demonstrated a purulent hemorrhagic abscess with fibrosis resulting in the loss of the drug-loaded implant before necropsy. Thus, while the score of the placebo implant (20) was relatively higher than the score of the other placebo implants, it was still lower than the score of the corresponding active implant (31). Overall, high levels of inflammation were observed in the peri-implant space in rhesus (ranging from an average total histological characteristic score of 24.7 ± 9.7 at week 12 (±SD, **Table 1**). These total histological characteristic scores were far higher than those observed for the placebo implants, which had an average score of 11.3 ± 8.2 at 12 weeks (see scores tables and micrographs from all animals in these studies in the **Suppl. 4**). Finally, the computed reaction grade indicated a *severe reaction* to the reduced dose and flux Generation B implant (see **Suppl. 5, Table S14**).

We sought to determine if the use of a trocar instead of surgical-pocket formation would modify the local inflammation. Here, two male and two female rhesus macaques were implanted with contralateral matched placebos and single Generation B implants. We observed that subdermal wounds in several animals developed into abscesses and surface lesions (see **Suppl. 8, Figure S31**). The use of a trocar allowed more efficient implant insertion with lower trauma but did not reduce or eliminate the local reaction to the Generation B implants.

## DISCUSSION

The dosing calculations for the design of the Generation A and B TAF subcutaneous implants were targeted to cellular benchmark concentrations of TFV-DP in PBMCs, established in humans from the STRAND, HPTN066, and iPrEX trials, all of which involved administration of TDF or TDF/FTC to seronegative individuals (20, 28, 29). Median PBMC TFV-DP concentrations in the STRAND trial were TFV-DP 42 fmol/10^6^ cells with once daily oral dosing (28). The HPTN066 study, conducted in healthy HIV negative individuals, was a dose-ranging DOT study to establish PrEP thresholds in plasma and PBMCs. PBMC TFV-DP concentrations >16.8 fmol/10^6^ cells were associated with daily adherence (29) at a 90% sensitivity threshold. Further, the iPrEX randomized control trial demonstrated that moderate adherence to PrEP was associated with 92% protection from seroconversion in men who have sex with men; a *post-hoc* analysis of PBMC TFV-DP concentrations estimated that PBMC-TFV-DP concentrations higher than 16 fmol/10^6^ cells are associated with protection with 90% sensitivity (28). Due to upwards of 66% TFV-DP losses during cryopreservation, using the iPrEX analysis, Gunawadarna *et al*. estimated a conservative EC_90_ of 24-48 fmol/10^6^ cells for PBMC TFV-DP concentrations (20). This TFV metabolite level was the benchmark applied in this work.

Both Generation A and Generation B implant formulations demonstrated the controlled release of TAF with sustained concentrations of intracellular TFV-DP throughout implant exposure (12 weeks). We found clear TAF dose dependence of cellular TFV-DP levels from the Generation A TAF implants in NZW rabbits (**Table 1**). TFV-DP in rabbit heterophils was found for all implant groups after week 1 and was observable throughout our study (**Figure 3 and 4**). Our lowest dose implant (*in vitro* release of 0.13 mg TAF/day) provided a median TFV-DP level of 42 fmol/10^6^ cells, a dose, and level of TFV-DP that is not extreme but still showed clear local drug-related histopathology (20). Our experiments demonstrate that we can achieve our benchmark cellular TFV-DP level using this reservoir implant at 10 μg/kg/day.

Demonstration of the lack of adverse local effects after implantation is required by the US FDA for all implanted medical devices (30, 31). To address all the stages of foreign body response to ARV releasing implants, we assessed the local response to the drug-eluting device sub-acutely and chronically (32). Critically, when evaluating the local implant site, one should not mechanically disturb the peri-implant cell-structure by extracting the implant from the tissue at necropsy. Instead, one must remove the tissue with the implant intact and fix the whole device and the surrounding tissue for histological analysis (25, 33). Also, one must section through the implant and not just collect dermal biopsies near the site. By fixing the implant and the surrounding tissue, we saw that the implant site was obviously demarcated in all histologic images and groups. Histopathology of the rabbit implant sites displayed evident local inflammation in all active Generation A implants (**Figure 5**).

We hypothesized that this inflammatory reaction might be TAF dose (mg/day) and TAF flux (mg/cm^2^/day) dependent. By reducing both, we hoped to attenuate the cellular inflammatory reactions shown above, while achieving protective levels of TFV-DP in PBMCs of primates. We also thought that there may be species-dependent toxicity in NZW rabbits that might not manifest itself in non-human primates. Thus, in our design of the Generation B subcutaneous TAF implants, we reduced our *in vitro* release rate from 0.39 mg/day to 0.13 mg/day, and reduced TAF flux from 0.24 to 0.08 mg TAF/cm^2^/day (**Table 1**). Assuming linear dose scaling from the Generation A implant in rhesus, we expected this implant to yield roughly 100 fmol/10^6^ cells of TFV-DP.

While we achieved our TFV-DP cellular benchmark PK levels in rhesus and the flux and dose were significantly reduced in Generation B, we continued to observe significant inflammation at 4 and 12 weeks in the peri-implant volume in rhesus macaques (**Figure 10**; for full histology reports, see **Suppl. 4**). Local inflammation and necrosis around the implant in all cases was much higher in the TAF implant arms than in the matched placebos, suggesting that inflammation is caused by TAF exposure to the local cells and tissue around the implant. This local inflammation occurred even at the lowest TAF dose and flux implant that achieves commonly used cellular TFV-DP benchmark levels. Furthermore, the lower inflammation surrounding the placebo implants strongly suggests that the local histopathology is neither due to the polymer nor due to the surgical procedure of blunt dissection or trocar administration. By comparing animals necropsied at four and 12 weeks, we observed a trend that inflammation appeared to become worse with time, with the active implants clearly more inflamed than the placebos at all times. Semiquantitative histopathology in primates computed a grade of severe inflammatory reaction and much higher than the cut-off score (*S̄_pair_*) of 15 (**Table S14**). Further evidence of the accumulating severity of the response is the loss of some implants through abscesses observed at 12 weeks.

Chua *et al.* are the only group to report data on TAF implants in rhesus macaques, and they exceeded benchmark levels of TFV-DP in PBMCs. With a dose in rhesus of 200 µg/day of TAF or approx. 20 µg/kg/day, they report a mean TFV-DP level of 533 fmol/10^6^ PBMCs. (23) Gundawarna et al. delivered controlled doses of 920 µg/day in beagle dogs with a corresponding dose of 85 µg/kg/day. (20) We delivered a lower dose of 130 µg/day in rhesus, or 10 µg/kg/day, and obtained lower median TFV-DP levels than Chua *et al*., yet we still observed unacceptable histopathology and inflammation around the active implants and not the placebos at a 10 µg/kg/day TAF dose (**Table 1**).

Although we are the first to report extensive histological data on TAF implants, we are not the first to report on implant safety or tissue pathology near an active TAF implant. When Chua *et al.* evaluated their system in rhesus macaques (23), they reported the presence of wound formation along the surgical incision or “dehiscence” over their TAF implant in two out of the three macaques, in addition to skin ulceration over the implant in two animals at day 70 (23). Ultimately, Chua *et al.* concluded that histopathological analysis of punch biopsy of skin sampled adjacent to the implant was normal in rhesus macaques, but no control implant data was collected. (23) It is not clear from Chua *et al.* if the whole implant was removed and fixed for the two published tissue sections. In a PCL implant similar to Schlesinger and Desai (21), Gatto and van der Straten reported on a “cutaneous response” to the devices in rabbits. (24) In Gundawarna *et al*., safety in beagle dogs was “evaluated by body weight and cage observations.”(20) No histopathological analysis data is provided, but from animal weight and cage observations, they concluded the implant was safe. However, in later unpublished pharmacokinetic studies in beagle dogs, this implant was associated with “erythema and/or edema at the implantation site” as well as “multiple instances of discharge”(34). It was argued that these reactions would not be observed at doses of less than 1 mg/day of TAF(34); our findings belie this expectation, as we observe inflammation at doses well below 1 mg/day in rhesus.

Earlier generations of our TAF implant design tended to leak because of poor sealing, resulting in high drug release. When these implants were evaluated in rhesus macaques, we observed an effect similar to a chemical burn that formed a hole in the primate skin. We believe that wound dehiscence, skin ulceration, and the ‘cutaneous response’ observations described above align with the results of the studies presented here.

TAF is safe when administered orally (14) and TFV has been safely administered vaginally, both as a gel (35, 36) and as an intravaginal ring (21, 37). It is well known that the route of administration can significantly modify the toxicological response to the drug delivery system. For example, the opiate receptor antagonist naltrexone is safe orally, but when formulated in an intramuscular injection of naltrexone microparticles this orally safe drug is reported not to be safe via injected dosage forms (38). Yamaguchi and Anderson subcutaneously injected sustained release microspheres containing placebo beads, naltrexone containing beads, and 100% naltrexone microspheres into rabbits and rats. They found significant inflammation located focally around naltrexone containing beads and microspheres, but not placebo beads (39). Entecavir, an antiretroviral used for the treatment of chronic hepatitis B, is another drug which is safe orally(40). However, Henry *et al.* evaluated polymer-coated pellets of entecavir in rats and observed local swelling, scab formation, and necrosis in their drug-loaded implants, but not their placebo(41). Similarly, we have observed little inflammation around the placebo implants, but observe significant inflammation around the drug-containing implants. Much as Yamaguchi and Anderson, we removed the entire implantation site for histopathological analysis after exposure to the drug delivery system.

Others have acknowledged that a possible cause of toxicity in eluting drug devices and formulations can arise from local high drug concentrations and high and long drug exposures to the local tissues around the implant (36). Subcutaneous implants inherently result in elevated local drug concentrations around the implant, since the entire dose is being released at the boundary of the device. In fact, in some cases, we were able to detect exceedingly high levels of TFV-DP in the tissue around the implant. For a set drug release rate, as the size of the device becomes small, the flux at the boundary must increase. This increased flux of drug should also increase exposure to the tissue local to the implant site. It is also possible that the fibrous capsule membrane, which forms as part of the foreign body response, can also act as a barrier to drug diffusion, thus causing higher local concentrations of eluted materials, which can lead to an even more exaggerated local response (30).

Additionally, the cellular toxicity may be caused by off-target effects of the nucleoside analog itself. Topical TFVs effect on gene expression is known (e.g. (42)). Previous tissue culture based on TFV has observed inhibition of wound healing in models using epithelial cells and fibroblasts isolated from the upper and lower human female reproductive tract (43). A phase 1 clinical trial of a TDF intravaginal ring was stopped early due to the development of Grade 1 vaginal ulceration near the ring after 32 days of use (44). Rodriguez-Garcia *et al.* observed significant inhibition of wound closure with one mg/mL (3277 μM) TFV and with concentrations of TAF higher than eight μM (43). Our results are, therefore consistent with previous observations of TFV’s interference with wound healing, but the exact molecular mechanisms that cause the observed local toxicity of these implants is unknown.

TAF undergoes pH-dependent hydrolysis into two main related substances (see **Suppl. 1**). Our work has not eliminated the possibility that differences in the interior microenvironment of other implant compositions could lead to other implants releasing a mixture of TAF related substances that differs from the implants described herein. These different mixtures of TAF related substances could result in modifications in the toxicokinetics of a TAF implant system. Because we have not studied this implant system side-by-side with the other published TAF implants, it remains a possibility that other differently constructed TAF implants will not suffer the same local toxicity issues we observed in our reservoirs.

## CONCLUSION

We describe a reservoir implant capable of delivering TAF in the subcutaneous space for a period of several months, and we tested this implant system in NZW rabbits and rhesus macaques for up to 12 weeks. We demonstrated that these implants could deliver TAF in a range of doses through modification of the implant geometry (wall thickness and length) and the material composition of the rate controlling membrane (see compositions of Generation A and B implants). Our main finding in this work is that this TAF implant always induces local inflammation around the implant at a low dose of the drug (10 µg/kg/day). We felt we could not reasonably decrease the dose or flux further and still generate a viable TAF implant that would protect from HIV transmission and be a reasonable size in humans. More importantly, our results could not exclude the possibility that this reservoir TAF implant, loaded with hundreds of milligrams of the drug, could leak from a manufacturing or mechanical failure and cause tissue damage to the user because of exposure to a large acutely applied dose of TAF in the sub-cutaneous volume. Together, these factors caused us to conclude this implant is unsafe and to terminate pre-clinical development efforts towards this TAF implant for long-acting HIV prevention and treatment.

Our work was informed by guidance for studying local inflammation around an implanted device described in ISO 10993-6 for the biological evaluation of medical devices. (32) ISO 10993-6 directs us to, “excise the implant site together with sufficient unaffected surrounding tissue to enable evaluation of the local histopathological response.” (32) Leaving the implant in the tissue allows histological evaluation of the pericapsular space around the implant without disturbing the fragile cellular and biopolymer structures that are used to measure histopathological response versus control. We followed this procedure, and our histological analysis clearly shows a local toxic response with the TAF loaded implant and a considerable reduction of such response in our matched placebos. We urge that the international standard (32) is followed for all future ARV implant local reaction studies. We also observed that the inflammation and necrosis around the implants became more severe at longer time points, suggesting that a one-month safety study would be insufficient for these long-acting devices. Finally, we recommend extensive stress testing of TAF and other ARV eluting devices to exclude rupture, device failure, and dose dumping.

While the oral route of administration of TAF (14, 45) is clinically proven to be safe, the TAF implants described herein are unsafe. Alternatively, another potent, long-acting small molecule ARV like cabotegravir (46–48) or GS-6297 (49) could be progressed with studies like those shown in this work to achieve the goal of a long-acting subcutaneous implant for the treatment and prevention of HIV infection. 4’-ethynyl-2-fluoro-2’-deoxyadenosine (EFdA, Islatravir, MK-8591) (46–48) is a highly potent nucleoside reverse transcriptase translocation inhibitor: once-weekly oral dosing has demonstrated its ability to protect male rhesus macaques in challenge studies(50). Implantable islatravir implants have demonstrated prophylactic concentrations in a human trial (51, 52). We further suggest that long-acting ARV drug substances should be evaluated for biocompatibility at the site of administration earlier in the preclinical development process. For example, an Alzet® osmotic pump delivering a set dose would permit the screening of local inflammation as a function of dose in an animal without requiring the full design of a drug delivery device capable of long-acting durations. Thus, similar reactions can be studied and avoided in future devices using other ARV drugs.

## MATERIALS AND METHODS

### Materials

TAF (CAS 1392275-56-7; GS-7340-03) and TFV (CAS 147127-20-6) were obtained from Gilead Sciences (Foster City, CA). Tecoflex® polyurethane was obtained from Lubrizol (Wickliffe, OH). Tips die, and the die head used for extrusion were sourced from Guill Tool (West Warwick, RI). The dies used to press pellets for TAF and placebo implants were purchased from Natoli (St Charles, MO). Sodium chloride and magnesium stearate (USP grade) used to manufacture implants were obtained from Spectrum Chemical (New Brunswick, NJ). Barium sulfate (USP grade) used in manufacturing radiopaque rods was obtained from Fisher Scientific (Fair Lawn, NJ). Sodium azide, ammonium acetate, phosphate buffered saline solution, and solvents used for high-performance liquid chromatography (HPLC) and mass spectrometry (36) work were obtained from Fisher Scientific (Fair Lawn, NJ). Isotopically-labeled [adenine-^13^C(U)] TFV (TFV*) was obtained from Moravek Biochemicals (Brea, CA). Syringe filter tips, weighing dishes, and centrifuge tubes were obtained from Fisherbrand (New Hampton, NH). Scintillation vials for *in vitro* release were obtained from Wheaton (Rockford, TN).

### Animal care and welfare

All animal studies were conducted in accordance with protocols approved by Northwestern University and Tulane National Primate Research Center Local Institutional Animal Care and Use Committees, Northwestern protocol IS00006125, Tulane protocol P0307R. This study was carried out in accordance with the Guide for the Care and Use of Laboratory Animals of the Institute of Laboratory Resources, National Resource Council. All procedures were performed under anesthesia using ketamine/xylazine, and all efforts were made to minimize stress, improve housing conditions, and provide enrichment opportunities. Animals were euthanized by sedation with ketamine/xylazine injection followed by intravenous barbiturate overdose in accordance with the recommendations of the panel on euthanasia of the American Veterinary Medical Association.

### Description of implant manufacturing

Implant manufacturing was completed in a non-sterile environment. However, all drug product-contacting surfaces including the benchtop surfaces, machines, and floors were cleaned with 3% hydrogen peroxide solution and ethanol. All materials used in manufacturing were depyrogenated by the heating glass and stainless-steel materials to 250 °C or rinsing heat-incompatible materials with 3% hydrogen peroxide solution. During manufacturing, all staff wore face masks, hairnets, disposable gowns, gloves, and shoe covers to minimize contamination.

Tecoflex™ EG-85A, and EG-85A:EG-93A (50:50 ratio) tubing was manufactured by hot melt extrusion, much as in Clark *et al.* (37). The EG-85A tubing was extruded on an ATR Plasti-Corder(R) single screw extruder (C.W. Brabender, South Hackensack, NJ), at a draw down ratio of 26.44 and draw balance ratio of 0.99 The zone temperatures for extrusion were 140/185/185/165/140 (all °C). The EG-85A:EG-93A (50:50 ratio) was first extruded into a rod using a twin screw extruder (C.W. Brabender, South Hackensack, NJ), to ensure homogeneity of the blend, with the zone temperatures set to 145/175/180/170/150 (all °C). The extruded rods were pelletized using a micro pelletizer (Randcastle Extrusion Systems Inc., Cedar Grove, NJ), and the pelletized material was extruded on the ATR plasticorder single screw extruder at a draw down ratio of 27.61 and draw balance ratio of 1.02, with the zone temperatures for extrusion set at 150/165/190/180/130 (all °C). The extruded tubing was sized for consistency. Tubing deviating by more than 10% from the required diameter and wall thickness was rejected. Tubing meeting the specifications were cut to length using a single-sided razor blade. 40% barium sulfate loaded EG-93A rods were manufactured by hot melt extrusion. Barium sulfate (40 wt%) was added to EG-93A and then extruded on the twin extruder. The zone temperatures for extrusion were 145/180/180/165/100 (all °C). The resulting extrudate was pelletized on the micro pelletizer. The 40% barium sulfate/EG93A pellets were then extruded at a diameter of 2.2 mm on the ATR plasticorder single screw extruder. The zone temperatures for extrusion were 155/180/180/150/120 (all °C). The 40% barium sulfate loaded EG-93A rods were cut into 3 mm long pellets using a single-sided razor blade. TAF was wet-granulated with NaCl at a ratio of 98:2 wt/wt with ethanol and the vacuum dried granulate was dry-coated with 2% magnesium stearate wt/wt. This mixture of powder was pressed into either 1.8- or 2.0-mm diameter pellets using a Natoli NP-RD10A (Natoli, St. Charles, MO) pellet press fitted with a 1.8 mm x 9.4 mm or 2.0 mm x 10 mm die/punch with the compression force set to 1000 lbs, and a fill depth of 3.3 mm. One end of the cut tubing was heat sealed using a Packworld PW2200 impulse sealer (Packworld, Nazareth, NJ) with the sealing conditions set with the following parameters: sealing temperature 120 °C, sealing time of 4 seconds, and percentage cooling before the clamp deactivates at 50%, and a sealing pressure of 60 psi. Pellets were loaded into the tubing. 1.8 mm or 2.0 mm diameter pellets of an equivalent amount of NaCl and magnesium stearate at a 1:1 ratio (wt/wt) were used for the placebo. For Generation B active and placebo implants, a 3 mm pellet of 40% barium sulfate compounded with the polyurethane Tecoflex EG-93A was also loaded into the tubing. The implant was then sealed at the other end using the heat sealer with the same impulse-sealing conditions described earlier. The ends were trimmed using surgical scissors. The total implant mass was recorded, and the implant placed in a polyethylene backed aluminum pouch (U-line, Pleasant Prairie, WI) which was heat sealed using an AIE300CA vacuum impulse sealer (American International Electric, City of Industry, CA). Implants were then annealed for 15 hours at 40 °C. The implants were shipped to Steri-tek Inc.

(Fremont, CA) for electron beam sterilization with a radiation dose of 25 kGy. Implants received back after sterilization were tested for endotoxin levels.

### Endotoxin testing

Endotoxin levels of all raw materials and pre/post-e-beam sterilized implants were quantified by chromogenic detection of LPS using the method provided by the Pierce Limulus Amebocyte Lysate Chromogenic Endotoxin Quantitation Kit. The FDA pre-defines endotoxin units (3) as 20.0 EU/device, which is approximately 0.5 EU/mL from a 40 mL rinse volume. All materials and implants used in the studies had levels below the assay LLOQ of 0.15 EU/mL. All materials used in this assay were purchased endotoxin or pyrogen-free. Samples were first fully submerged in 3 mL of Ficoll-Plaque Plus endotoxin-free water (GE Healthcare, Uppsala, Sweden) in either a 6-well culture plate or 3-mL test tube for 1 hour, at 37°C, and 800 RPM on a Multitherm benchmark (Benchmark, Edison, NJ). Samples were analyzed (n=3) alongside a positive control spiked sample to a concentration of 0.8 EU/mL on a 96-well, polystyrene bottom plate using a Biotek Synergy 2 Multimode Microplate Reader (Biotek, Winooski, VT) at 405-410 nm wavelength. Any sample with enhanced or inhibited interference signal outside the range of the calibration curve (0.15-2.0 EU/mL) was diluted in endotoxin-free water and reanalyzed.

### The Initial selection of dose in NZW rabbits

To start our dose ranging studies, we allometrically scaled the lowest dose in our rabbit PK study to roughly Oak Crest’s maximum estimate(20) for what they believe to be an efficacious TAF exposure in humans; this turns out to be 0.1 mg/day in a 3.2 kg rabbit. The other rabbit doses we chose were scaled in multiples from this dose and are 0.2 mg/day, 0.4 mg/day, and 0.6 mg/day (Table 1). This would provide a range of doses starting at their maximum estimate of a TAF dose, twice that, and up to six times that in rabbits. Allometrically in the rabbit, our highest dose exceeds the dose that Oak Crest tested in beagles. This range should allow us to develop a PK dose response and toxicokinetic curves and ranges over the doses that should be required for TAF. We have manufactured implant systems that achieve the TAF doses shown in Table 2, and these are described below.

**Table 2.** Description of Generation A implant systems used in the NZW PK and safety study.

| Animal Group ID | *In vitro* release rate<sup>a</sup> (mg/day) | Material | Pellet Diameter (mm) | Implant Diameter (mm) | Implant Lumen Length (cm) | Membrane Thickness (cm) | Implant Strength of TAF (mg ±SD) | Number of Active Implants per Animal |
| --- | --- | --- | --- | --- | --- | --- | --- | --- |
| 1 | 0.13 | Tecoflex EG-85A | 1.8 | 2.2 | 0.8 | 0.015 | 16.8±0.3 | 1 |
| 2 | 0.26 | Tecoflex EG-85A | 1.8 | 2.2 | 0.8 | 0.015 | 16.8±0.3 | 2 |
| 3 | 0.52 | Tecoflex EG-85A | 1.8 | 2.2 | 1.6 | 0.015 | 34.0±0.3 | 2 |
| 4 | 0.78 | Tecoflex EG-85A | 1.8 | 2.2 | 1.6 | 0.015 | 34.0±0.3 | 3 |
| 5 | Placebo | Tecoflex EG-85A | 1.8 | 2.2 | 1.6 | 0.015 | ±0 | 1 or 2 |

a. Calculated from average release rate Fig. 1-3 once the implants reach steady-state over days 7 to 91

### TAF implant formulation characteristics

We evaluated two generations of TAF implants: Generation A and Generation B. Generation A were evaluated in NZW rabbits for PK and safety and rhesus macaques for PK. Characteristics and dimensions of the Generation A implants are in Tables 1 and 2. Based on results from Generation A, Generation B implants with lower doses were manufactured and evaluated in rhesus macaques. Dimensions and characteristics for Generation B are given in Table 3.

**Table 3.**
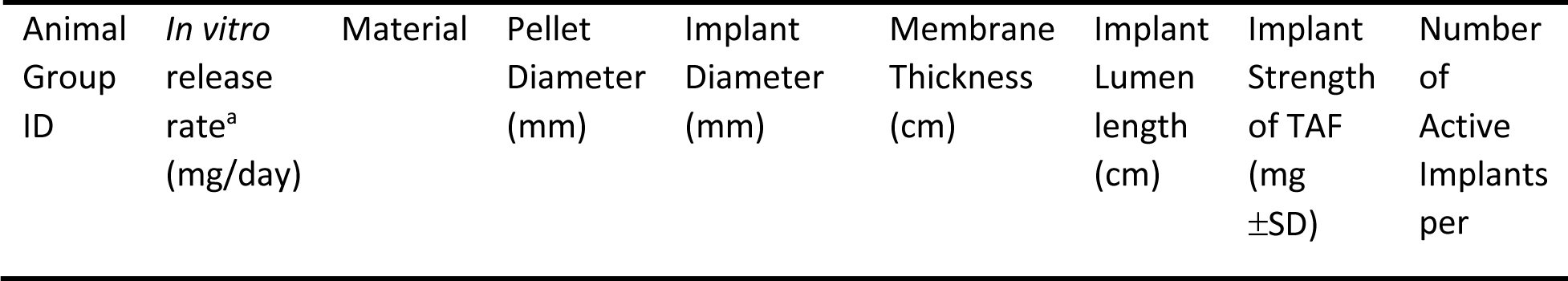
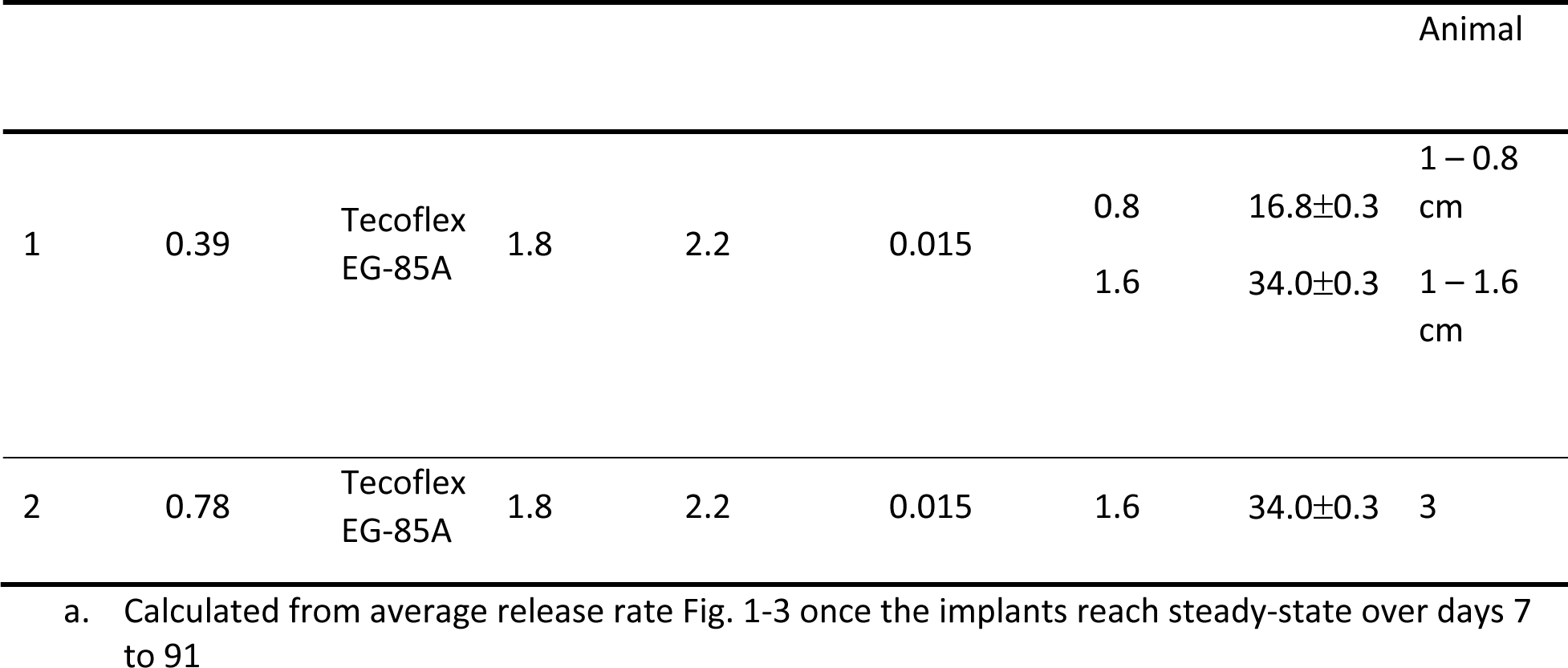
Generation A formulation characteristics and description of manufacturing used in macaques:

**Table 4.** Generation B formulation characteristics and description of manufacturing used in macaques:

| *In vitro* release rate<sup>a</sup> (mg/day) | Material | Pellet Diameter (mm) | Implant Diameter (mm) | Membrane Thickness (mm) | Implant Lumen Length (cm) | Implant Strength of TAF (mg ±SD) |
| --- | --- | --- | --- | --- | --- | --- |
| 0.13 | Tecoflex EG-85A:EG-93A 50:50 | 2.0 | 2.6 | 0.17 | 2.0 | 44.4±1.7 |
| Placebo | Tecoflex EG-85A:EG-93A 50:50 | 2.0 | 2.6 | 0.17 | 2.0 | NA |

a. Calculated from average release rate Fig. 1-3 once the implants reach steady-state over days 7 to 91

### Animal study design, group sizes, and controls

Overall, as mentioned above, we conducted four animal PK studies. The first two sets of studies used the Generation A TAF implant in both NZW rabbits and rhesus macaques. The third study used the findings from the first studies to justify a dose reduction and evaluate a Generation B TAF implant in rhesus macaques. The fourth study used a trocar to implant the devices. The sampling schedule for all studies remained the same (Table 5).

**Table 5.** Design of PK and safety studies. PK and Safety studies followed the same schedule for TAF Generation A in NZW rabbits and TAF Generation B in macaques. Necropsy of the animals was performed at weeks 4 and 12. All necropsy was accompanied by histology and staining. In Generation A implant studies with rhesus macaques only a PK study was performed with no necropsy at the end of the study.

| Proceedure | Time (Weeks) |  |  |  |  |  |  |  |  |  |  |  |  |
| --- | --- | --- | --- | --- | --- | --- | --- | --- | --- | --- | --- | --- | --- |
|  | 0 | 1 | 2 | 3 | 4 | 5 | 6 | 7 | 8 | 9 | 10 | 11 | 12 |
| Implant | x |  |  |  |  |  |  |  |  |  |  |  |  |
| Plasma TFV, TAF | x | x | x | x | x | x | x | x | x | x | x | x | x |
| PBMC TFV-DP | x | x | x | x | x | x | x | x | x | x | x | x | x |
| Vaginal Swab | x |  | x |  | x |  | x |  | x |  | x |  | x |
| Rectal Swab | x |  | x |  | x |  | x |  | x |  | x |  | x |
| Vaginal Biopsy | x |  |  |  | x |  |  |  | x |  |  |  | x |
| Rectal Biopsy | x |  |  |  | x |  |  |  | x |  |  |  | x |
| Necropsy<sup>a</sup> |  |  |  |  | x |  |  |  |  |  |  |  | x |

a. Necropsy of the rhesus macaques were not performed in the Generation A NHP implant study and therefore was only a PK study.

The rabbit study had five groups of female NZW rabbits. In the placebo group, there were three animals. In Group 1, seven animals received a 0.8 cm active implant and a contralateral placebo. In Groups 2 and 3, seven animals each received two 0.8 cm or two 1.6 cm active implants, respectively. In Group 4, six animals received three 1.6 cm implants. Group 5 acted as the control group and consisted of three animals implanted with two placebos each. Two of seven animals in Groups 1, 2, and 3, and three of six in Group 4 were sacrificed during Week 4. Five of seven animals in Groups 1, 2, and 3, three of six in Group 4, and all three placebo controls were sacrificed during Week 12. The average body mass of the rabbits was 3.37 ± 0.21 kg with a range of 2.84 to 3.81 kg.

The macaque Generation A exploratory PK study included two groups, each of which contained three females. The first group received one 0.8 cm implant and one 1.6 cm implant. The second group received three 1.6 cm implants. Animals were maintained on study for 12 weeks, and the mass of the animals had a mean of 6.5 ± 0.3 kg with a range of 6.2 to 7.0 kg. The macaque Generation B study had four groups of animals with two animals each. Each animal received two implants: one placebo and one active implant (both 2 cm) contralaterally. Group 1 (one male and one female) was necropsied at four weeks, and Group 2 (two females) at 12 weeks. Blood, tissue, and PBMCs were still sampled per the overall schedule. All animals had at least one active implant. The rhesus macaques had mean body mass of 12.7 ± 4.4 kg and ranged from 7 to 19.6 kg.

The use of contralateral implantation of implants provided us several opportunities to simultaneously study *in vivo* drug release rates of the implants and local irritation. Contralateral implantation of an active implant and a placebo implant was used in the Generation A rabbit studies (Group 1) and the Generation B rhesus macaque studies (all macaques). In animals with two or more contralateral active implants, one implant was resected in a block of tissue with the intact implant, fixed and sectioned to evaluate local inflammation around the implant histologically. The other implant was removed and tested for *in vitro* performance. These implants were subjected to *in vitro* release, leak testing, and the polymer molecular weight distribution was determined (see **Suppl. 6**). For each implant we tracked its initial mass, the strength of the pellet, and the total pellet mass; this allowed to determine total drug content after explantation and obtain the average daily release rate of the devices over the three-month study by subtraction.

Finally, we conducted a fourth exploratory PK and safety study in rhesus macaques to evaluate if insertion of the Generation B device with a trocar would modify the biological response to the implant. Here there were two groups of two males and two females implanted contralaterally with an active and matched placebo using a 4.5 mm trocar kit. Other than the method of insertion and the number of animals, the study design was identical. The macaques had an average mass of 14.24 ± 3.6 kg and a mass range of 9.4 to 18.0 kg.

### The surgical subcutaneous implantation procedure

Before the surgery, the animals were anesthetized with ketamine/xylazine, and both the blood as well as vaginal and rectal swabs were collected. For implants in the first three studies, two small incisions were made in the skin, and a blunt probe was gently inserted under the skin to ensure room for the implant. The implants were atraumatically and gently inserted between the shoulder blades, and the wound was sutured closed. In the case of the fourth study in rhesus, a 4.5 mm trocar kit (Elemis Corp., Carson City, NV) was used to insert the devices, and the wound was sutured closed.

### Blood collection

Blood samples were collected before the surgical subcutaneous implantation procedure from the ear vein (rabbits) and femoral vein (macaques) and then weekly for up to 12 weeks. Approximately 8 mL of blood was taken from the ear or femoral vein, transferred into EDTA-coated tubes and immediately placed on ice. Additionally, 1 mL of blood was collected before the surgeries and every four weeks for complete blood count and chemistries.

### Plasma isolation

Whole blood samples were taken from the ear (rabbits) or femoral vein (macaques) were collected into EDTA-coated tubes and immediately placed on ice. The tubes were centrifuged (2100 rpm, 20 min, 4°C), to separate plasma from the cells. Isolated plasma, present in the upper layer of the sample, was transferred to new tubes and stored at −80°C for further PK analysis.

### Cell isolation and TFV-DP extraction

Rabbit heterophils and macaque mononuclear cells were isolated from the peripheral blood (2) obtained from the ear (rabbits) or femoral (macaques) vein. Briefly, following blood collection the samples were placed immediately on ice and then centrifuged (2100 rpm, 20 min, 4°C). Next, plasma was removed, and PBS (∼1 mL) was added to the remaining blood. Then, the samples were layered over lymphocyte separation media and centrifuged for 20 min at 2100 rpm to remove red blood cells. The layer containing the PBMCs was collected and transferred into 15 mL centrifuge tubes. Cells were pelleted by centrifugation (1500 rpm, 7 min) and re-suspended in 2 mL of PBS. Then, the cells were counted with 1:10 dilution using Turk’s solution in a counting chamber. The cells were pelleted again (1500 rpm, 7 min) and re-suspended in 2 mL of ice-cold 70% methanol. This solution was then split into two 1 mL vials and stored at −80°C for further PK analysis.

### Histology

Tissue samples located near the implantation site as well as samples of liver, lung, spleen, kidney, vagina, and rectum were collected and fixed in z-fix, embedded in paraffin and cut into tissue sections. Paraffin-embedded specimens were stained with hematoxylin and eosin. Stained tissue sections were evaluated for the signs of the presence and severity of inflammation. Tissue samples from naïve animals were used as control. Samples were blinded and scored by a trained pathologist.

### Histology and scoring algorithm

We used a semiquantitative histological scoring system to identify and characterize the presence of cellular and tissue responses in high powered fields of slides of the peri-implant space. **Table 6** displays the semiquantitative scoring system derived from recommendations in ISO 10993-6:2016 Annex E. (32). Blinded slides were provided to a pathologist for scoring. Cells were counted per high power field (HPF), and a table with scores was filled out as a summary from multiple slides along the length of each fixed implant (e.g., see **Suppl. 3**). Key characteristics that were evaluated are polymorphonuclear cells (heterophils in rabbits and neutrophils in macaques), lymphocytes, plasma cells, macrophages, giant cells, necrosis, capsule thickness, and tissue infiltrate (see **Table 6**).

**Table 6.** Histological characteristics scoring scheme used to evaluate the cellular and tissue characteristics observed near the implants.

| | Score | | | | | Reactive<br>inflammation<br>multiplier<br>$(m_{j,r})$ |
| --- | --- | --- | --- | --- | --- | --- |
|  | 0 | 1 | 2 | 3 | 4 |  |
| **Cell**<br>**characteristic** |  |  |  |  |  |  |
| Polymorphonuclear cells | 0 /<br>HPF | Rare,<br>1-5/HPF | 5-10/HPF | Heavy infiltrate | Packed | 2 |
| Lymphocytes | 0 /<br>HPF | Rare,<br>1-5/HPF | 5-10/HPF | Heavy infiltrate | Packed | 2 |
| Plasma cells | 0 /<br>HPF | Rare,<br>1-5/HPF | 5-10/HPF | Heavy infiltrate | Packed | 2 |
| Macrophages | 0 /<br>HPF | Rare,<br>1-5/HPF | 5-10/HPF | Heavy infiltrate | Packed | 2 |
| Giant cells | 0 /<br>HPF | Rare,<br>1-2/HPF | 3-5/HPF | Heavy infiltrate | Sheets | 2 |
| **Tissue**<br>**characteristic** |  |  |  |  |  |  |
| Necrosis | 0 | Minimal | Mild | Moderate | Severe | 2 |
| Capsule thickness | 0 | Narrow<br>band<br>($<5$ cells) | Moderate<br>(5-10<br>cells) | Thick band (10-20 cells) | Extensive<br>thick band | 1 |
| Tissue infiltrate | 0 | Minimal<br>focal<br>invasion<br>of local<br>tissue | Mild to<br>multifocal<br>inflammation in<br>adjacent<br>tissues | Moderate<br>inflammation in<br>adjacent tissues | Marked<br>inflammation in<br>adjacent<br>tissues | 1 |

We report the scores of the characteristics in two ways. In **Table 1** we simply sum the cellular and tissue characteristic scores and take the average across all the implants in a group to compute the total histological characteristic score. Secondly, we computed an implant reactivity grade for each implant type tested using equation 1.

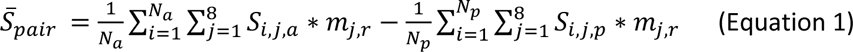

To compute the implant reactivity grade, we first summed the product of each histological characteristic score (*S_j,a_* for placebo and *S_j,p_* for an active implant’s characteristic scores with *j=1 to 8* for all 8 characteristics used in the histological analysis) and each characteristic’s reactive inflammation multiplier (*m_j,r_*) (see **Table 6**) for each implant. This sum was then averaged over all implants of that type (*N_p_* for the number of placebos and *N_a_* for the number of active implants in each group). We call this score for each type the average implant reactivity score that varies from 0 to 48. Inflammatory cellular infiltrate characteristics and the tissue characteristic of necrosis receive an inflammatory reaction multiplier of two to represent the greater importance of inflammation in the endpoints of these studies. Next, the average implant reactivity scores for the active and placebo treatments were subtracted to compute the average placebo adjusted implant reactivity score (*S̄_pair_*). Finally, the implant reactivity grade was determined by lookup as follows: minimal to no reaction (*S̄_pair_* from 0.0 up to 2.9), slight reaction (*S̄_pair_* from 3.0 up to 8.9), moderate reaction (*S̄_pair_* from 9.0 up to 15.0), and severe reaction (*S̄_pair_* >15.1) as per the published standard (32). See **Suppl. 5, Table S14**.

### *In vitro* release testing of implants

*In vitro* release was conducted under sink conditions in 20 mL scintillation vials at 37°C and 80 RPM in an I26 Incubator/Shaker (New Brunswick Scientific, Edison, NJ) using 20 mL of 1×PBS with 0.02% (w/v) sodium azide. Media was changed daily. Sink conditions were maintained at TAF concentrations < 1/10 the maximum solubility of the drug. TAF from *in vitro* samples was measured on a Zorbax Eclipse Plus C18 column (4.6×100 mm, 3.5 µm Agilent, Santa Clara, CA) and a Zorbax Eclipse Plus C18 guard column (4.6×12.5 mm, 5 µm Agilent, Santa Clara, CA) on an Agilent 1200 series HPLC attached to a UV based diode array detector at 260 nm (Agilent, Santa Clara, CA). TAF was separated by gradient elution using 20 mM ammonium acetate buffer (mobile phase A) and acetonitrile (mobile phase B) over 12.5 minutes (gradient t=0: %B=0, t=8: %B=55, t= 9.6: %B=55, t=9.7: %B= 0) at 0.7 mL/min. The column thermostat was set to 25 °C and samples were injected at ambient temperature. Data were acquired and processed using Agilent Chemstation software (Agilent, Santa Clara, CA). Calibration curves were prepared using on column amounts of TAF (40 and 800 µg/mL) and TFV (10 and 100 µg/mL) standards injected at 1, 2, 5, and 10 µL volumes. Sample concentrations were determined using the linear regression of calibration curves to measure TAF and its TFV-related species (PMPA monoamidate, monophenyl PMPA, and TFV). The TAF calibration curve was used to quantitate TAF and PMPA monoamidate, and the TFV calibration curve was used to quantitate TFV and monophenyl PMPA. The sum of TAF, TFV-related species (PMPA monoamidate and monophenyl PMPA), and TFV were quantified as TAF equivalent and reported as the total mass of TAF release.

### *In vitro* release testing and TAF extraction from explanted implants

Implants and surrounding tissue were removed together to preserve underlying tissue histology, as described above. While some implant/tissue specimens were preserved and sectioned, other implants were removed intact from animals for further analysis. Recovered implants were placed on *in vitro* release testing and analyzed as described in the *in vitro* release testing section. Samples were collected daily for two weeks to verify daily release and ensure that release properties from our actual *in vivo* implants were as predicted.

After 14 days, implants were removed from *in vitro* conditions, and TAF was extracted from the implant. The exterior of each implant was carefully dried after removal from the buffer, before being transferred to a scintillation vial and frozen. This was done to minimize the loss of drug while cutting the implant. Implant controls and samples were weighed, cut in half in an aluminum weighing dish with a scalpel, and quantitatively transferred from the dish into a 25 mL volumetric flask using methanol. The samples were shaken overnight until dissolved and diluted to volume. The resulting solution was then diluted 1:10 (v/v) in methanol and passed through a 0.2 μm nylon syringe filter tip into an HPLC vial for analysis. Extracted samples were analyzed using the HPLC method described in the *in vitro* release testing section.

### Extracting and analyzing TFV in rabbit plasma

TFV was separated from rabbit plasma through liquid/liquid extraction and analyzed by ultra-performance liquid chromatography – tandem mass spectrometry (UPLC-MS/MS) analysis. All calibration standards and quality controls (QCs) were prepared using 100 μL of sterile rabbit plasma obtained from GeneTex (Irvine, CA), 50 μL of a 100.8 nM solution of in (TFV*) water TFV*, 250 μL of MS grade acetonitrile, and 50 µL of TFV dissolved in MS grade water. This solution was used to generate a standard curve from 0.2 nM to 2000 nM with low, middle, and high QCs at 111 nM, 222 nM, 1111 nM. Samples were similarly prepared by spiking 100 μL of collected rabbit plasma with 50 μL of TFV*, 50 μL of MS grade water, and 250 μL of MS grade acetonitrile. Spiked solutions were mixed at 400 rpm for 10 min, centrifuged at 10,000 rpm for 10 min, and passed through a 0.2 μm nylon syringe filter tip into a centrifuge tube. Finally, 200 μL of each extract was transferred to a 2 mL, 96-well Nunc DeepWell plate (ThermoSci, Waltham, MA) and vacu-fixed at 50°C for 2 hrs in a Savant SPD111V SpeedVac concentrator (ThermoSci, Waltham, MA). The final plate was reconstituted in 200 μL of MS-grade water, capped with a silicone mat (Axygen, Corning, NY), and mixed at 400 rpm for 30 min at 37°C.

Samples were analyzed by UPLC-MS/MS at 5 µL injection volumes on a Zorbax RRHD Eclipse Plus C18 column (2.1×50mm, 1.8 μm; Agilent, Santa Clara, CA) using a Shimadzu Nexera X2 UHPLC (Shimadzu, Columbia, MD) with a SciEx QTRAP 6500+ mass spectrometer (SciEx, Redwood City, CA). Analytes were separated by gradient at a flow rate of 0.75 mL/min using 0.5% acetic acid in water (mobile phase A) and 0.5% acetic acid in methanol (mobile phase B) over 3.55 minutes (gradient - t=0 min: 0 %B; t=0.5 min: 0 %B; t=2.0 min: 100 %B; t=2.1 min: 0% B). The column thermostat was held at 40 °C while the sample chamber in the autosampler was cooled to 15 °C.

The SciEx detector was set to positive ion mode. A multiple reaction monitoring scan was used to detect transitions for TFV from m/z 288.1 to 176.1, TFV* from m/z 293.1 to 181.2, TAF from m/z 477.1 to 346.1, monophenyl PMPA from m/z 364.1 to 176.2, and PMPA monoamidate from 401.2 to 270.1. Dwell time was set to 100 msec for all analytes except TFV*, which was set for 50 msec. The collision energy was set to 30 eV for TAF, 25 eV for PMPA monoamidate, and 34 eV for TFV, TFV*, and monophenyl PMPA. Curtain gas, nebulizer gas, and auxiliary gas were set to 25 psi, 50 psi, and 55 psi, respectively. Ion spray voltage and source temperature were set to 5500 V and 400 °C. Declustering potential was set to 55 V. Entrance potential, and collision cell exit potential were both set to 10 V. Data was acquired, processed, and quantified using Analyst software (SciEx, Redwood City, CA). TFV in samples was quantified by using linear regression of area under curve ratios of TFV and TFV* from standard preparations.

### Pharmacokinetic measurements

Quantification of TAF, TFV, and TFV-DP in all matrices except TFV rabbit plasma (see above) was conducted by the Johns Hopkins University School of Medicine Clinical Pharmacology Analytical Laboratory and measurements were conducted using previously described liquid chromatographic-mass spectrometric (LC-MS/MS) approaches (53). TFV-DP quantification in PBMCs and tissue was conducted using a previously described, indirect enzymatic approach (54). All assays were validated in accordance with FDA, Guidance for Industry: Bioanalytical Method Validation and assay calibrators and QCs were prepared using human material (55). Assay lower limits of quantification were as follows: plasma TFV, 0.31 ng/mL; plasma TAF: 0.03 ng/mL; tissue TFV, 0.05 ng/sample; PBMC and tissue TFV-DP: 50 fmol or 5 fmol/sample. TFV-DP concentrations were normalized to several cells tested, for final reporting as fmol/10^6^ cells. Cell counts were provided by the collecting site. Tissue results were normalized to the net weight of tissue sample analyzed and reported as ng/mg (TFV) and fmol/mg (TFV-DP). Both TFV and TFV-DP were measured in the same biopsy specimen for all analyses. It is noted that the tissue assays were validated using a human luminal (vaginal, rectal) tissue matrix. Quantification of TAF, TFV, and TFV-DP in all matrices was conducted by the Johns Hopkins University School of Medicine Clinical Pharmacology Analytical Laboratory and measurements were conducted using previously described LC-MS/MS approaches (53). TFV-DP quantification in PBMCs and tissue was conducted using a previously described, indirect enzymatic approach (54). All assays were validated in accordance with FDA, Guidance for Industry: Bioanalytical Method Validation and assay calibrators and QCs were prepared using human material (55). Assay lower limits of quantification were as follows: plasma TFV, 0.31 ng/mL; plasma TAF: 0.03 ng/mL; tissue TFV, 0.05 ng/sample; PBMC and tissue TFV-DP: 50 fmol or 5 fmol/sample. TFV-DP concentrations were normalized to several cells tested, for final reporting as fmol/10^6^ cells. Cell counts were provided by the collecting site. Tissue results were normalized to the net weight of tissue sample analyzed and reported as ng/mg (TFV) and fmol/mg (TFV-DP). Both TFV and TFV-DP were measured in the same biopsy specimen for all analyses. It is noted that the tissue assays were validated using a human luminal (vaginal, rectal) tissue matrix.

## ACKNOWLEDGMENTS

We thank Gilead Sciences, Inc., for generously supplying the drug substance. We give special thanks to James Anderson, M.D., Ph.D. of Case Western University for his help in guiding our animal implant study designs. We also thank Prof. Craig Hendrix Ph.D. of Johns Hopkins University., Prof. Peter Anton, M.D of UCLA., Jim Rooney, Ph.D. of Gilead Sciences Inc., and Meredith Clark, Ph.D. of CONRAD, for helpful discussions, and Meagan Watkins of Tulane University, for her assistance with the project. This work was funded by The National Institute of Allergy and Infectious Diseases of the National Institutes of Health under award number UM1 AI120184. This work made use of the IMSERC at Northwestern University, which has received support from the Soft and Hybrid Nanotechnology Experimental (SHyNE) Resource (NSF ECCS-1542205); the State of Illinois and International Institute for Nanotechnology (IIN).

## Supplemental 1. HPLC chromatograms for TFV, monophenyl PMPA, PMPA monoamidate, and TAF parent from a generation B TAF implant on days 5 and 63 of in vitro release testing

**Figure S1.**
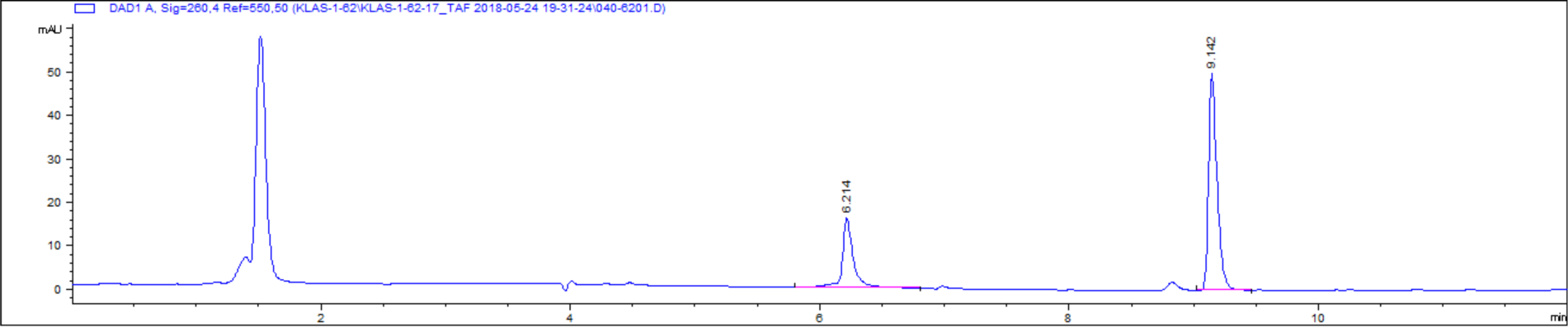
Chromatograph showing relative peaks areas and retention times for PMPA monoamidate (RT=6.21 min) and TAF parent (RT=9.14 min) from a generation B TAF implant on day 5 of *in vitro* release testing

**Figure S2.**
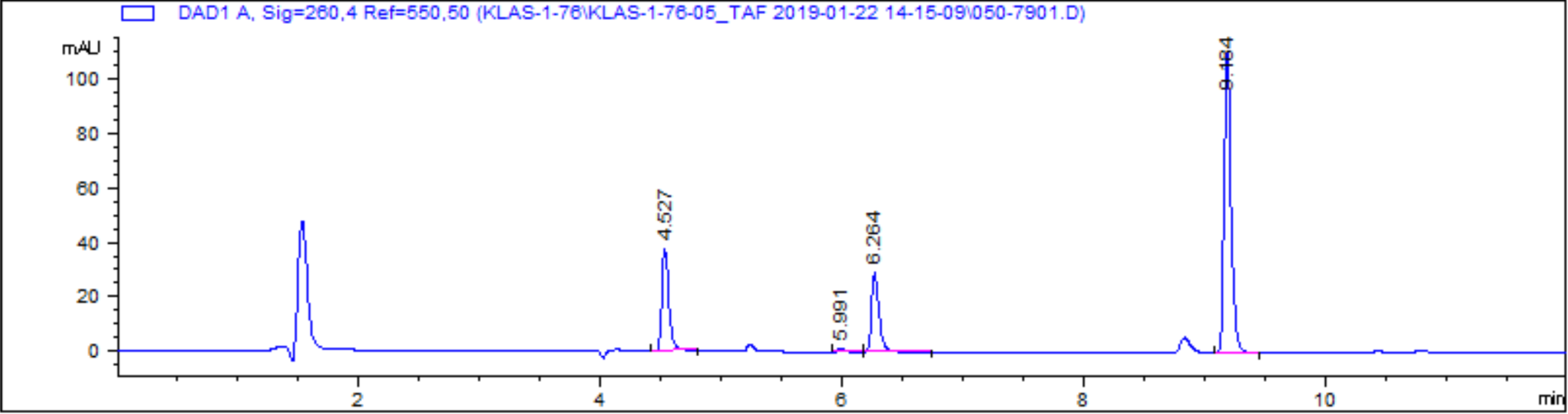
Chromatograph showing relative peak areas and retention times for TFV (RT=4.53 min), monophenyl PMPA (RT=5.99 min), PMPA monoamidate (RT=6.26 min), and TAF parent (RT=9.184 min) from a generation B TAF implant on day 63 of *in vitro* release testing

**Figure S3.**
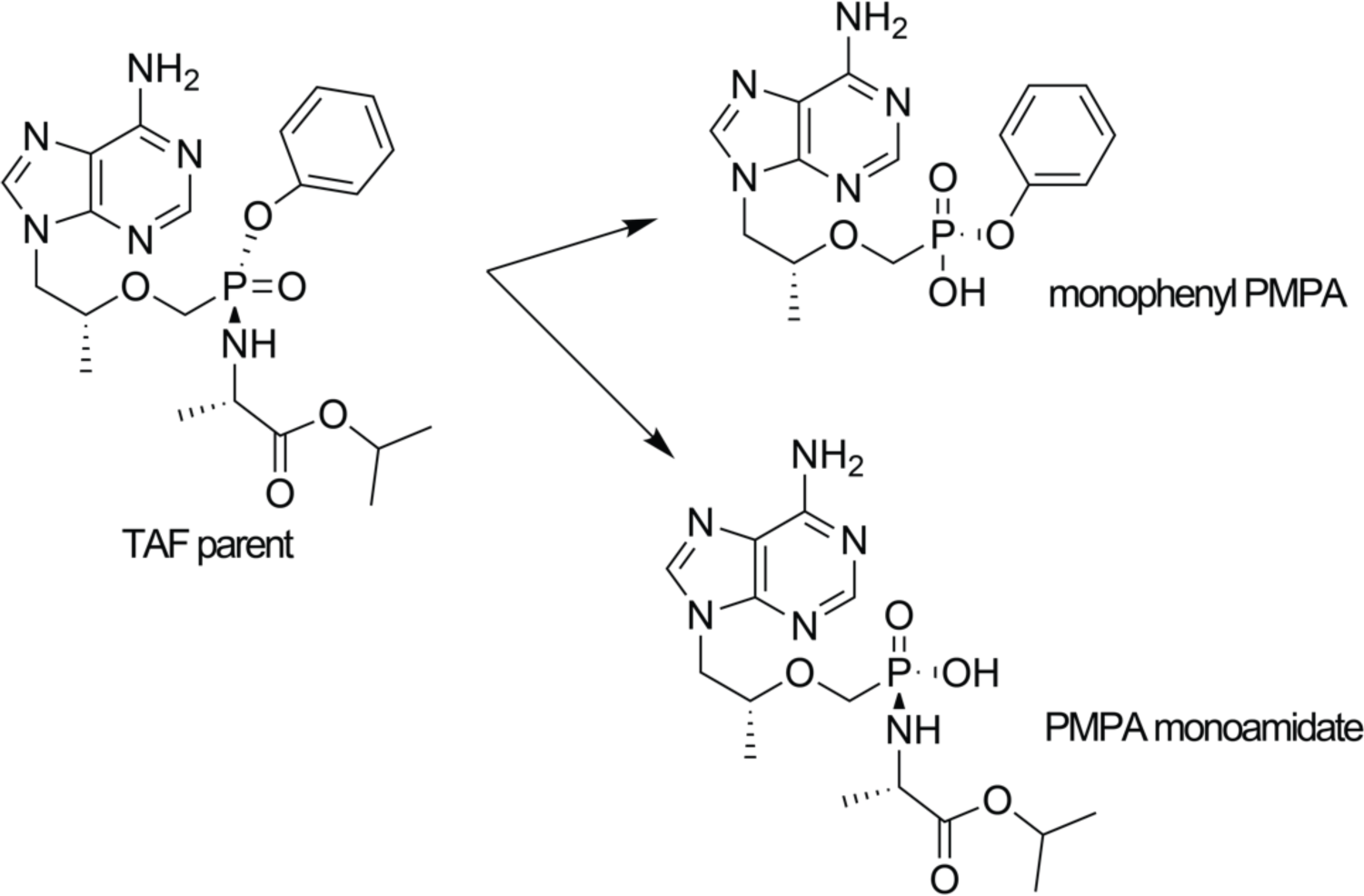
TAF parent is tenofovir alafenamide without the fumarate salt, which undergoes pH-dependent hydrolysis into two main related substances monophenyl PMPA and PMPA monoamidate.

**Figure S4.**
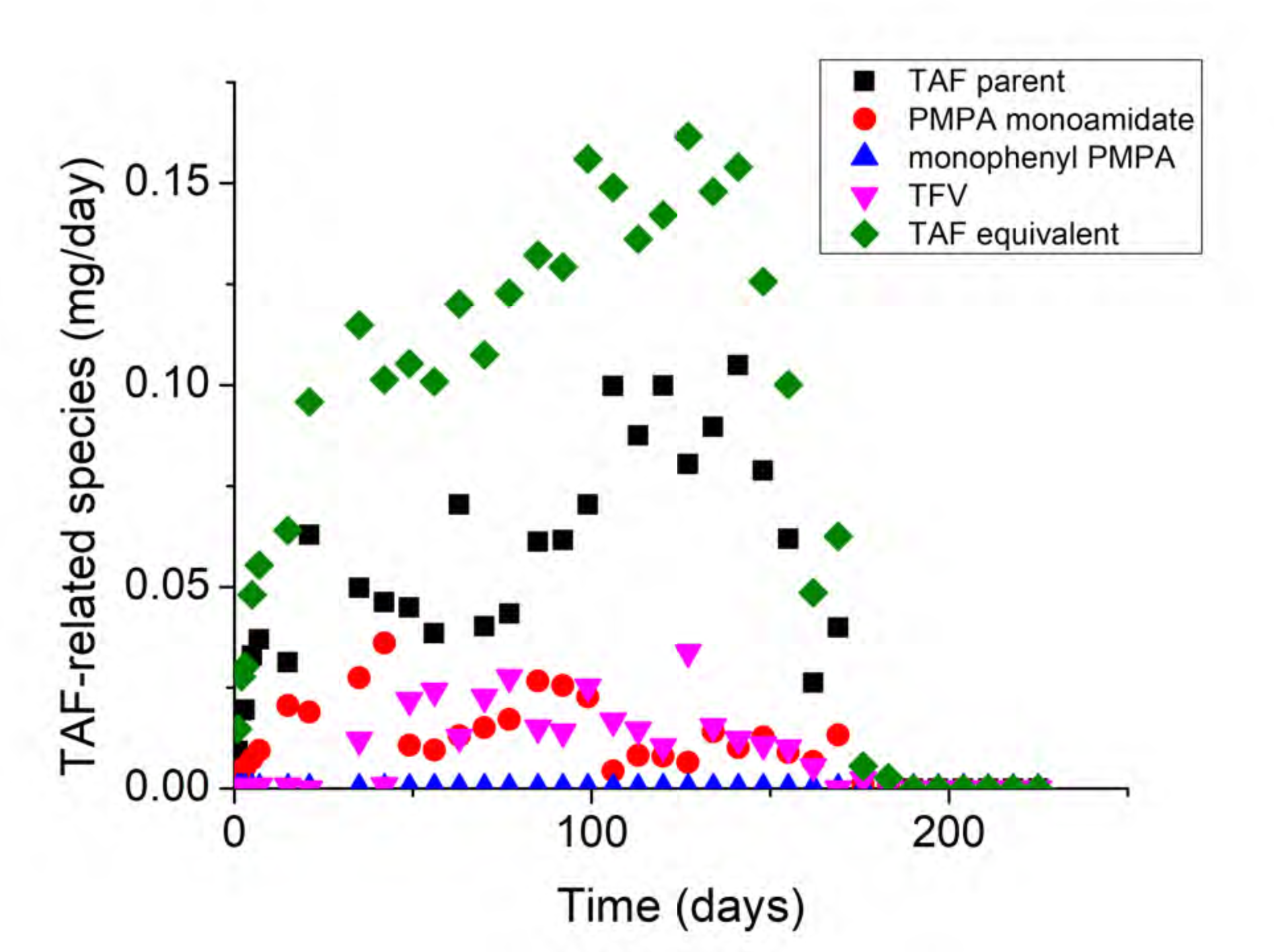
Plot of TAF equivalent and TAF-related species over time.

## Supplemental 2. TFV-DP Tissue and TFV plasma levels in New Zealand White rabbits in generation A implants

**Table S1.** Generation A NZW rabbit TFV-DP tissue levels

| Animal ID | *In vitro*<br>release rate<br>(mg/day) | Time<br>Point<br>(weeks) | Specimen Type | Final [TFVdp]<br>(fmol/mg) |
| --- | --- | --- | --- | --- |
| M29035 | 0.13 | 4 | Implant Tissue Left | 2.06 |
| M29036 | 0.13 | 4 | Implant Tissue Left | BLQ |
| M29037 | 0.26 | 4 | Implant Tissue Left | 3.27 |
| M29038 | 0.26 | 4 | Implant Tissue Left | BLQ |
| M29039 | 0.48 | 4 | Implant Tissue Left | 32.22 |
| M29040 | 0.48 | 4 | Implant Tissue Left | 1537.81 |
| M29041 | 0.72 | 4 | Implant Tissue Left | 11930.48 |
| M29042 | 0.72 | 4 | Implant Tissue Left | 168.26 |
| M29043 | 0.72 | 4 | Implant Tissue Left | 11222.4 |
| M29044 | 0 | 12 | Implant Tissue Left | BLQ |
| M29045 | 0 | 12 | Implant Tissue Left | BLQ |
| M29046 | 0 | 12 | Implant Tissue Left | BLQ |
| M29047 | 0.13 | 12 | Implant Tissue Left | 0.859 |
| M29048 | 0.13 | 12 | Implant Tissue Left | 1.62 |
| M29049 | 0.13 | 12 | Implant Tissue Left | 0.851 |
| M29050 | 0.13 | 12 | Implant Tissue Left | 1.587 |
| M29051 | 0.13 | 12 | Implant Tissue Left | 5.66 |
| M29052 | 0.26 | 12 | Implant Tissue Left | 4102 |
| M29053 | 0.26 | 12 | Implant Tissue Left | 2.36 |
| M29054 | 0.26 | 12 | Implant Tissue Left | 2938 |
| M29055 | 0.26 | 12 | Implant Tissue Left | 10.5 |
| M29056 | 0.26 | 12 | Implant Tissue Left | 114.6 |
| M29057 | 0.48 | 12 | Implant Tissue Left | 44509 |
| M29058 | 0.48 | 12 | Implant Tissue Left | 8.58 |
| M29059 | 0.48 | 12 | Implant Tissue Left | 3427 |
| M29060 | 0.48 | 12 | Implant Tissue Left | 30.7 |
| M29062 | 0.72 | 12 | Implant Tissue Left | 16136 |
| M29063 | 0.72 | 12 | Implant Tissue Left | 10386 |
| M29064 | 0.72 | 12 | Implant Tissue Left | 120 |
| M29036 | 0.13 | 4 | Implant Tissue Right | 2.38 |
| M29037 | 0.13 | 4 | Implant Tissue Right | BLQ |
| M29038 | 0.26 | 4 | Implant Tissue Right | BLQ |
| M29039 | 0.26 | 4 | Implant Tissue Right | 948.60 |
| M29040 | 0.48 | 4 | Implant Tissue Right | 2032.36 |
| M29041 | 0.48 | 4 | Implant Tissue Right | 7291.24 |
| M29042 | 0.72 | 4 | Implant Tissue Right | 305.66 |
| M29044 | 0 | 12 | Implant Tissue Right | BLQ |
| M29045 | 0 | 12 | Implant Tissue Right | BLQ |
| M29046 | 0 | 12 | Implant Tissue Right | BLQ |
| M29047 | 0.13 | 12 | Implant Tissue Right | 214 |
| M29048 | 0.13 | 12 | Implant Tissue Right | 1644 |
| M29049 | 0.13 | 12 | Implant Tissue Right | 126 |
| M29050 | 0.13 | 12 | Implant Tissue Right | 69941 |
| M29051 | 0.13 | 12 | Implant Tissue Right | 6371 |
| M29052 | 0.26 | 12 | Implant Tissue Right | 1.31 |
| M29053 | 0.26 | 12 | Implant Tissue Right | 86.3 |
| M29054 | 0.26 | 12 | Implant Tissue Right | 56.7 |
| M29055 | 0.26 | 12 | Implant Tissue Right | 9132 |
| M29056 | 0.26 | 12 | Implant Tissue Right | 30.3 |
| M29057 | 0.48 | 12 | Implant Tissue Right | 1726 |
| M29058 | 0.48 | 12 | Implant Tissue Right | 27.5 |
| M29059 | 0.48 | 12 | Implant Tissue Right | 8.18 |
| M29060 | 0.48 | 12 | Implant Tissue Right | 7.75 |
| M29061 | 0.72 | 12 | Implant Tissue Right | 12113 |
| M29062 | 0.72 | 12 | Implant Tissue Right | 6422 |
| M29064 | 0.72 | 12 | Implant Tissue Right | 360 |
| M29035 | 0.13 | 4 | Vaginal Tissue | 3.67 |
| M29036 | 0.13 | 4 | Vaginal Tissue | 4.25 |
| M29037 | 0.26 | 4 | Vaginal Tissue | 8.08 |
| M29038 | 0.26 | 4 | Vaginal Tissue | 3.29 |
| M29039 | 0.48 | 4 | Vaginal Tissue | 34.81 |
| M29040 | 0.48 | 4 | Vaginal Tissue | 17.14 |
| M29041 | 0.72 | 4 | Vaginal Tissue | 169.36 |
| M29042 | 0.72 | 4 | Vaginal Tissue | 63.35 |
| M29043 | 0.72 | 4 | Vaginal Tissue | 98.74 |
| M29044 | 0 | 12 | Vaginal Tissue | BLQ |
| M29045 | 0 | 12 | Vaginal Tissue | BLQ |
| M29046 | 0 | 12 | Vaginal Tissue | BLQ |
| M29047 | 0.13 | 12 | Vaginal Tissue | 7.05 |
| M29048 | 0.13 | 12 | Vaginal Tissue | 7.09 |
| M29049 | 0.13 | 12 | Vaginal Tissue | 7.74 |
| M29050 | 0.13 | 12 | Vaginal Tissue | 13.3 |
| M29051 | 0.13 | 12 | Vaginal Tissue | 2.83 |
| M29052 | 0.26 | 12 | Vaginal Tissue | 3.58 |
| M29053 | 0.26 | 12 | Vaginal Tissue | 17 |
| M29054 | 0.26 | 12 | Vaginal Tissue | 19.7 |
| M29055 | 0.26 | 12 | Vaginal Tissue | 14.9 |
| M29056 | 0.26 | 12 | Vaginal Tissue | 8.49 |
| M29057 | 0.48 | 12 | Vaginal Tissue | 62.1 |
| M29058 | 0.48 | 12 | Vaginal Tissue | 14.8 |
| M29059 | 0.48 | 12 | Vaginal Tissue | 24.3 |
| M29060 | 0.48 | 12 | Vaginal Tissue | 22.6 |
| M29061 | 0.48 | 12 | Vaginal Tissue | 51.6 |
| M29062 | 0.72 | 12 | Vaginal Tissue | 73 |
| M29063 | 0.72 | 12 | Vaginal Tissue | 77.6 |
| M29064 | 0.72 | 12 | Vaginal Tissue | 37.1 |
| M29035 | 0.13 | 4 | Rectal Tissue | 2.79 |
| M29036 | 0.13 | 4 | Rectal Tissue | 3.96 |
| M29037 | 0.26 | 4 | Rectal Tissue | 6.33 |
| M29038 | 0.26 | 4 | Rectal Tissue | BLQ |
| M29039 | 0.48 | 4 | Rectal Tissue | 49.95 |
| M29040 | 0.48 | 4 | Rectal Tissue | 10.22 |
| M29042 | 0.72 | 4 | Rectal Tissue | 31.45 |
| M29043 | 0.72 | 4 | Rectal Tissue | 46.97 |
| M29044 | 0 | 12 | Rectal Tissue | BLQ |
| M29045 | 0 | 12 | Rectal Tissue | BLQ |
| M29046 | 0 | 12 | Rectal Tissue | BLQ |
| M29047 | 0.13 | 12 | Rectal Tissue | 3.32 |
| M29048 | 0.13 | 12 | Rectal Tissue | 2 |
| M29049 | 0.13 | 12 | Rectal Tissue | 2.63 |
| M29050 | 0.13 | 12 | Rectal Tissue | 4.26 |
| M29051 | 0.13 | 12 | Rectal Tissue | 3.44 |
| M29052 | 0.26 | 12 | Rectal Tissue | 3.08 |
| M29053 | 0.26 | 12 | Rectal Tissue | 23.4 |
| M29054 | 0.26 | 12 | Rectal Tissue | 8.49 |
| M29055 | 0.26 | 12 | Rectal Tissue | 13.6 |
| M29056 | 0.26 | 12 | Rectal Tissue | 7.08 |
| M29057 | 0.48 | 12 | Rectal Tissue | 43.9 |
| M29058 | 0.48 | 12 | Rectal Tissue | 30.1 |
| M29059 | 0.48 | 12 | Rectal Tissue | 18.9 |
| M29060 | 0.48 | 12 | Rectal Tissue | 6.51 |
| M29061 | 0.48 | 12 | Rectal Tissue | 25.3 |
| M29062 | 0.72 | 12 | Rectal Tissue | 23.5 |
| M29063 | 0.72 | 12 | Rectal Tissue | 47.1 |
| M29064 | 0.72 | 12 | Rectal Tissue | 29.6 |

**Table S2.** Gen A plasma TFV concentrations in NZW rabbits.

| **Specimen ID** | ***In vitro* release rate (mg/day)** | **Time Point (Week)** | **Calculated Concentration (ng/mL)** |
| --- | --- | --- | --- |
| M29035 | 0.13 | Week 0 | BLQ |
| M29035 | 0.13 | Week 1 | BLQ |
| M29035 | 0.13 | Week 2 | 1.73 |
| M29035 | 0.13 | Week 3 | BLQ |
| M29035 | 0.13 | Week 4 | BLQ |
| M29036 | 0.13 | Week 0 | BLQ |
| M29036 | 0.13 | Week 1 | BLQ |
| M29036 | 0.13 | Week 2 | BLQ |
| M29036 | 0.13 | Week 3 | BLQ |
| M29036 | 0.13 | Week 4 | BLQ |
| M29037 | 0.26 | Week 0 | BLQ |
| M29037 | 0.26 | Week 1 | 4.33 |
| M29037 | 0.26 | Week 2 | 2.16 |
| M29037 | 0.26 | Week 3 | 2.8 |
| M29037 | 0.26 | Week 4 | 4.36 |
| M29038 | 0.26 | Week 0 | BLQ |
| M29038 | 0.26 | Week 1 | 1.98 |
| M29038 | 0.26 | Week 2 | BLQ |
| M29038 | 0.26 | Week 3 | 1.89 |
| M29038 | 0.26 | Week 4 | 1.65 |
| M29039 | 0.48 | Week 0 | BLQ |
| M29039 | 0.48 | Week 1 | 4.77 |
| M29039 | 0.48 | Week 2 | 4.97 |
| M29039 | 0.48 | Week 3 | 4.76 |
| M29039 | 0.48 | Week 4 | 4.79 |
| M29040 | 0.48 | Week 0 | BLQ |
| M29040 | 0.48 | Week 1 | 4.49 |
| M29040 | 0.48 | Week 2 | 3.51 |
| M29040 | 0.48 | Week 3 | 4.7 |
| M29040 | 0.48 | Week 4 | 5.16 |
| M29041 | 0.72 | Week 0 | 5.36 |
| M29041 | 0.72 | Week 1 | 4.95 |
| M29041 | 0.72 | Week 2 | 4.93 |
| M29041 | 0.72 | Week 3 | 6.45 |
| M29041 | 0.72 | Week 4 | 5.62 |
| M29042 | 0.72 | Week 0 | BLQ |
| M29042 | 0.72 | Week 1 | 4.07 |
| M29042 | 0.72 | Week 2 | 4.42 |
| M29042 | 0.72 | Week 3 | 5.43 |
| M29042 | 0.72 | Week 4 | 5.91 |
| M29043 | 0.72 | Week 0 | BLQ |
| M29043 | 0.72 | Week 1 | 4.9 |
| M29043 | 0.72 | Week 2 | 6.01 |
| M29043 | 0.72 | Week 3 | 6.75 |
| M29043 | 0.72 | Week 4 | 5.78 |
| M29044 | Placebo | Week 0 | BLQ |
| M29044 | Placebo | Week 1 | BLQ |
| M29044 | Placebo | Week 2 | BLQ |
| M29044 | Placebo | Week 3 | BLQ |
| M29044 | Placebo | Week 4 | BLQ |
| M29044 | Placebo | Week 5 | BLQ |
| M29044 | Placebo | Week 6 | BLQ |
| M29044 | Placebo | Week 7 | BLQ |
| M29044 | Placebo | Week 8 | BLQ |
| M29044 | Placebo | Week 9 | BLQ |
| M29044 | Placebo | Week 10 | BLQ |
| M29044 | Placebo | Week 11 | BLQ |
| M29044 | Placebo | Week 12 | BLQ |
| M29045 | Placebo | Week 0 | BLQ |
| M29045 | Placebo | Week 1 | BLQ |
| M29045 | Placebo | Week 2 | BLQ |
| M29045 | Placebo | Week 3 | BLQ |
| M29045 | Placebo | Week 4 | 11.5 |
| M29045 | Placebo | Week 6 | BLQ |
| M29045 | Placebo | Week 7 | BLQ |
| M29045 | Placebo | Week 8 | BLQ |
| M29045 | Placebo | Week 10 | BLQ |
| M29045 | Placebo | Week 11 | BLQ |
| M29045 | Placebo | Week 12 | BLQ |
| M29046 | Placebo | Week 0 | BLQ |
| M29046 | Placebo | Week 1 | BLQ |
| M29046 | Placebo | Week 2 | BLQ |
| M29046 | Placebo | Week 3 | BLQ |
| M29046 | Placebo | Week 4 | BLQ |
| M29046 | Placebo | Week 5 | BLQ |
| M29046 | Placebo | Week 6 | BLQ |
| M29046 | Placebo | Week 7 | BLQ |
| M29046 | Placebo | Week 8 | BLQ |
| M29046 | Placebo | Week 9 | BLQ |
| M29046 | Placebo | Week 10 | BLQ |
| M29046 | Placebo | Week 11 | BLQ |
| M29046 | Placebo | Week 12 | BLQ |
| M29047 | 0.13 | Week 0 | BLQ |
| M29047 | 0.13 | Week 1 | BLQ |
| M29047 | 0.13 | Week 2 | 2.15 |
| M29047 | 0.13 | Week 3 | 1.67 |
| M29047 | 0.13 | Week 4 | BLQ |
| M29047 | 0.13 | Week 5 | BLQ |
| M29047 | 0.13 | Week 6 | BLQ |
| M29047 | 0.13 | Week 7 | 3.29 |
| M29047 | 0.13 | Week 8 | 2.29 |
| M29047 | 0.13 | Week 9 | 2.31 |
| M29047 | 0.13 | Week 10 | 2.22 |
| M29047 | 0.13 | Week 11 | 1.79 |
| M29047 | 0.13 | Week 12 | BLQ |
| M29048 | 0.13 | Week 0 | BLQ |
| M29048 | 0.13 | Week 1 | BLQ |
| M29048 | 0.13 | Week 2 | BLQ |
| M29048 | 0.13 | Week 3 | BLQ |
| M29048 | 0.13 | Week 4 | BLQ |
| M29048 | 0.13 | Week 5 | 2.06 |
| M29048 | 0.13 | Week 6 | 2.09 |
| M29048 | 0.13 | Week 7 | 1.92 |
| M29048 | 0.13 | Week 8 | BLQ |
| M29048 | 0.13 | Week 9 | BLQ |
| M29048 | 0.13 | Week 10 | 5.23 |
| M29048 | 0.13 | Week 11 | 1.67 |
| M29048 | 0.13 | Week 12 | BLQ |
| M29049 | 0.13 | Week 0 | BLQ |
| M29049 | 0.13 | Week 1 | BLQ |
| M29049 | 0.13 | Week 2 | BLQ |
| M29049 | 0.13 | Week 3 | BLQ |
| M29049 | 0.13 | Week 4 | BLQ |
| M29049 | 0.13 | Week 5 | 1.55 |
| M29049 | 0.13 | Week 6 | 3.61 |
| M29049 | 0.13 | Week 7 | BLQ |
| M29049 | 0.13 | Week 8 | BLQ |
| M29049 | 0.13 | Week 9 | 3.3 |
| M29049 | 0.13 | Week 10 | 3.2 |
| M29049 | 0.13 | Week 11 | BLQ |
| M29049 | 0.13 | Week 12 | BLQ |
| M29050 | 0.13 | Week 0 | BLQ |
| M29050 | 0.13 | Week 1 | BLQ |
| M29050 | 0.13 | Week 2 | 1.68 |
| M29050 | 0.13 | Week 3 | BLQ |
| M29050 | 0.13 | Week 4 | BLQ |
| M29050 | 0.13 | Week 5 | 2.73 |
| M29050 | 0.13 | Week 6 | 3.02 |
| M29050 | 0.13 | Week 7 | 1.82 |
| M29050 | 0.13 | Week 8 | BLQ |
| M29050 | 0.13 | Week 9 | 1.65 |
| M29050 | 0.13 | Week 10 | 19.7 |
| M29050 | 0.13 | Week 11 | BLQ |
| M29050 | 0.13 | Week 12 | BLQ |
| M29051 | 0.13 | Week 0 | BLQ |
| M29051 | 0.13 | Week 1 | BLQ |
| M29051 | 0.13 | Week 2 | 1.93 |
| M29051 | 0.13 | Week 3 | 1.53 |
| M29051 | 0.13 | Week 4 | BLQ |
| M29051 | 0.13 | Week 5 | BLQ |
| M29051 | 0.13 | Week 6 | 1.59 |
| M29051 | 0.13 | Week 7 | BLQ |
| M29051 | 0.13 | Week 8 | BLQ |
| M29051 | 0.13 | Week 9 | BLQ |
| M29051 | 0.13 | Week 10 | 10.1 |
| M29051 | 0.13 | Week 11 | 2.48 |
| M29051 | 0.13 | Week 12 | BLQ |
| M29052 | 0.26 | Week 0 | BLQ |
| M29052 | 0.26 | Week 1 | BLQ |
| M29052 | 0.26 | Week 2 | 4.82 |
| M29052 | 0.26 | Week 3 | 2.55 |
| M29052 | 0.26 | Week 4 | 2.34 |
| M29052 | 0.26 | Week 5 | 4.47 |
| M29052 | 0.26 | Week 6 | 4.36 |
| M29052 | 0.26 | Week 7 | 3.55 |
| M29052 | 0.26 | Week 8 | 2.85 |
| M29052 | 0.26 | Week 9 | 3.39 |
| M29052 | 0.26 | Week 10 | 4.38 |
| M29052 | 0.26 | Week 11 | 2.64 |
| M29052 | 0.26 | Week 12 | BLQ |
| M29053 | 0.26 | Week 0 | BLQ |
| M29053 | 0.26 | Week 1 | 2.02 |
| M29053 | 0.26 | Week 2 | 2.38 |
| M29053 | 0.26 | Week 3 | 3.08 |
| M29053 | 0.26 | Week 4 | 1.94 |
| M29053 | 0.26 | Week 5 | 2.95 |
| M29053 | 0.26 | Week 6 | 3.32 |
| M29053 | 0.26 | Week 7 | 2.19 |
| M29053 | 0.26 | Week 8 | 1.84 |
| M29053 | 0.26 | Week 9 | 2.78 |
| M29053 | 0.26 | Week 10 | 7.32 |
| M29053 | 0.26 | Week 11 | 3.66 |
| M29053 | 0.26 | Week 12 | 2.47 |
| M29054 | 0.26 | Week 0 | BLQ |
| M29054 | 0.26 | Week 1 | 2.01 |
| M29054 | 0.26 | Week 2 | 4.04 |
| M29054 | 0.26 | Week 3 | 2.96 |
| M29054 | 0.26 | Week 4 | 2.78 |
| M29054 | 0.26 | Week 5 | 2.86 |
| M29054 | 0.26 | Week 6 | 5.72 |
| M29054 | 0.26 | Week 7 | 2.29 |
| M29054 | 0.26 | Week 8 | 2.52 |
| M29054 | 0.26 | Week 9 | 2.98 |
| M29054 | 0.26 | Week 10 | 7.99 |
| M29054 | 0.26 | Week 11 | 3.55 |
| M29054 | 0.26 | Week 12 | 1.53 |
| M29055 | 0.26 | Week 0 | BLQ |
| M29055 | 0.26 | Week 1 | 3.93 |
| M29055 | 0.26 | Week 2 | 4.28 |
| M29055 | 0.26 | Week 3 | 2.42 |
| M29055 | 0.26 | Week 5 | 4.84 |
| M29055 | 0.26 | Week 6 | 3.77 |
| M29055 | 0.26 | Week 7 | 2.41 |
| M29055 | 0.26 | Week 9 | 2.29 |
| M29055 | 0.26 | Week 10 | 1.59 |
| M29055 | 0.26 | Week 12 | 2.53 |
| M29056 | 0.26 | Week 0 | BLQ |
| M29056 | 0.26 | Week 1 | 2.53 |
| M29056 | 0.26 | Week 2 | 3 |
| M29056 | 0.26 | Week 3 | 3.03 |
| M29056 | 0.26 | Week 4 | 2.68 |
| M29056 | 0.26 | Week 5 | 7.26 |
| M29056 | 0.26 | Week 6 | 4.32 |
| M29056 | 0.26 | Week 7 | 2.5 |
| M29056 | 0.26 | Week 8 | 2.78 |
| M29056 | 0.26 | Week 9 | 3.17 |
| M29056 | 0.26 | Week 11 | 3.15 |
| M29056 | 0.26 | Week 12 | 3.54 |
| M29057 | 0.48 | Week 0 | BLQ |
| M29057 | 0.48 | Week 1 | 7.5 |
| M29057 | 0.48 | Week 2 | 6.72 |
| M29057 | 0.48 | Week 3 | 6.06 |
| M29057 | 0.48 | Week 4 | 5.75 |
| M29057 | 0.48 | Week 5 | 5.76 |
| M29057 | 0.48 | Week 6 | 7.59 |
| M29057 | 0.48 | Week 7 | 5.7 |
| M29057 | 0.48 | Week 8 | 5.8 |
| M29057 | 0.48 | Week 9 | 7.26 |
| M29057 | 0.48 | Week 10 | 6.46 |
| M29057 | 0.48 | Week 11 | 9.23 |
| M29057 | 0.48 | Week 12 | 5.28 |
| M29058 | 0.48 | Week 0 | BLQ |
| M29058 | 0.48 | Week 1 | 5.52 |
| M29058 | 0.48 | Week 2 | 3.58 |
| M29058 | 0.48 | Week 3 | 4.21 |
| M29058 | 0.48 | Week 4 | 3.65 |
| M29058 | 0.48 | Week 5 | 4.49 |
| M29058 | 0.48 | Week 6 | 7.43 |
| M29058 | 0.48 | Week 7 | 6.27 |
| M29058 | 0.48 | Week 8 | 6.69 |
| M29058 | 0.48 | Week 9 | 6.51 |
| M29058 | 0.48 | Week 10 | 6.26 |
| M29058 | 0.48 | Week 11 | 7.64 |
| M29058 | 0.48 | Week 12 | 7.55 |
| M29059 | 0.48 | Week 1 | 4.63 |
| M29059 | 0.48 | Week 3 | 7.22 |
| M29059 | 0.48 | Week 4 | 7 |
| M29059 | 0.48 | Week 5 | 6.56 |
| M29059 | 0.48 | Week 6 | 8.94 |
| M29059 | 0.48 | Week 7 | 5.27 |
| M29059 | 0.48 | Week 8 | 5.93 |
| M29059 | 0.48 | Week 9 | 8.54 |
| M29059 | 0.48 | Week 10 | BLQ |
| M29059 | 0.48 | Week 12 | 1.9 |
| M29060 | 0.48 | Week 0 | BLQ |
| M29060 | 0.48 | Week 1 | 2.74 |
| M29060 | 0.48 | Week 2 | 2.45 |
| M29060 | 0.48 | Week 3 | 4.17 |
| M29060 | 0.48 | Week 4 | 3.43 |
| M29060 | 0.48 | Week 5 | 4.96 |
| M29060 | 0.48 | Week 6 | 5.92 |
| M29060 | 0.48 | Week 7 | 4.37 |
| M29060 | 0.48 | Week 8 | 4.44 |
| M29060 | 0.48 | Week 9 | 5.65 |
| M29060 | 0.48 | Week 10 | 12.2 |
| M29060 | 0.48 | Week 11 | 6.08 |
| M29060 | 0.48 | Week 12 | 5.03 |
| M29061 | 0.48 | Week 0 | BLQ |
| M29061 | 0.48 | Week 1 | 3.49 |
| M29061 | 0.48 | Week 2 | 7.17 |
| M29061 | 0.48 | Week 3 | 5.99 |
| M29061 | 0.48 | Week 4 | 6.76 |
| M29061 | 0.48 | Week 5 | 7.88 |
| M29061 | 0.48 | Week 6 | 9.38 |
| M29061 | 0.48 | Week 7 | 6.09 |
| M29061 | 0.48 | Week 8 | 7.47 |
| M29061 | 0.48 | Week 9 | 10.2 |
| M29061 | 0.48 | Week 10 | 4 |
| M29061 | 0.48 | Week 11 | 6.67 |
| M29061 | 0.48 | Week 12 | 8.28 |
| M29062 | 0.72 | Week 0 | BLQ |
| M29062 | 0.72 | Week 1 | 5.29 |
| M29062 | 0.72 | Week 2 | 7.24 |
| M29062 | 0.72 | Week 3 | 6.47 |
| M29062 | 0.72 | Week 4 | 8.2 |
| M29062 | 0.72 | Week 5 | 11.3 |
| M29062 | 0.72 | Week 6 | 8.1 |
| M29062 | 0.72 | Week 7 | 8.04 |
| M29062 | 0.72 | Week 8 | 6.55 |
| M29062 | 0.72 | Week 9 | 5.35 |
| M29062 | 0.72 | Week 10 | 10.4 |
| M29062 | 0.72 | Week 11 | 7.33 |
| M29062 | 0.72 | Week 12 | 5.71 |
| M29063 | 0.72 | Week 0 | BLQ |
| M29063 | 0.72 | Week 1 | 4.48 |
| M29063 | 0.72 | Week 2 | 5.5 |
| M29063 | 0.72 | Week 3 | 5.63 |
| M29063 | 0.72 | Week 4 | 8.68 |
| M29063 | 0.72 | Week 5 | 8.4 |
| M29063 | 0.72 | Week 6 | 11.1 |
| M29063 | 0.72 | Week 7 | 7.29 |
| M29063 | 0.72 | Week 8 | 8.74 |
| M29063 | 0.72 | Week 9 | 6.82 |
| M29063 | 0.72 | Week 10 | 7.95 |
| M29063 | 0.72 | Week 11 | 6.32 |
| M29063 | 0.72 | Week 12 | 7.97 |
| M29064 | 0.72 | Week 0 | BLQ |
| M29064 | 0.72 | Week 1 | 4.22 |
| M29064 | 0.72 | Week 2 | 5.81 |
| M29064 | 0.72 | Week 3 | 6.83 |
| M29064 | 0.72 | Week 4 | 6.08 |
| M29064 | 0.72 | Week 5 | 11.5 |
| M29064 | 0.72 | Week 6 | 7.72 |
| M29064 | 0.72 | Week 7 | 5.58 |
| M29064 | 0.72 | Week 8 | 7.92 |
| M29064 | 0.72 | Week 9 | 7.74 |
| M29064 | 0.72 | Week 10 | 7.23 |
| M29064 | 0.72 | Week 11 | 5.41 |
| M29064 | 0.72 | Week 12 | 3.46 |

## Supplemental 3. 12 week histology reports in New Zealand White rabbits in generation A implants

### Placebos

**Table S3.** Histological characteristic scores from four placebo implants in rabbits M29044 and M29045.

|  | Response |  |  |  |
| --- | --- | --- | --- | --- |
| Animal | M29044 |  | M29045 |  |
| Implant | Left | Right | Left | Right |
| Polymorphonuclear cells | 0 | 0 | 0 | 0 |
| Lymphocytes | 0 | 0 | 0 | 0 |
| Plasma cells | 0 | 0 | 0 | 0 |
| Macrophages | 0 | 0 | 0 | 0 |
| Giant cells | 0 | 0 | 0 | 0 |
| Necrosis | 0 | 0 | 0 | 0 |
| Capsule thickness | 2 | 2 | 1 | 1 |
| Tissue infiltrate | 0 | 0 | 0 | 0 |
| Other |  |  |  |  |
| **Subtotal** | **2** | **2** | **1** | **1** |

**Figure S5.**
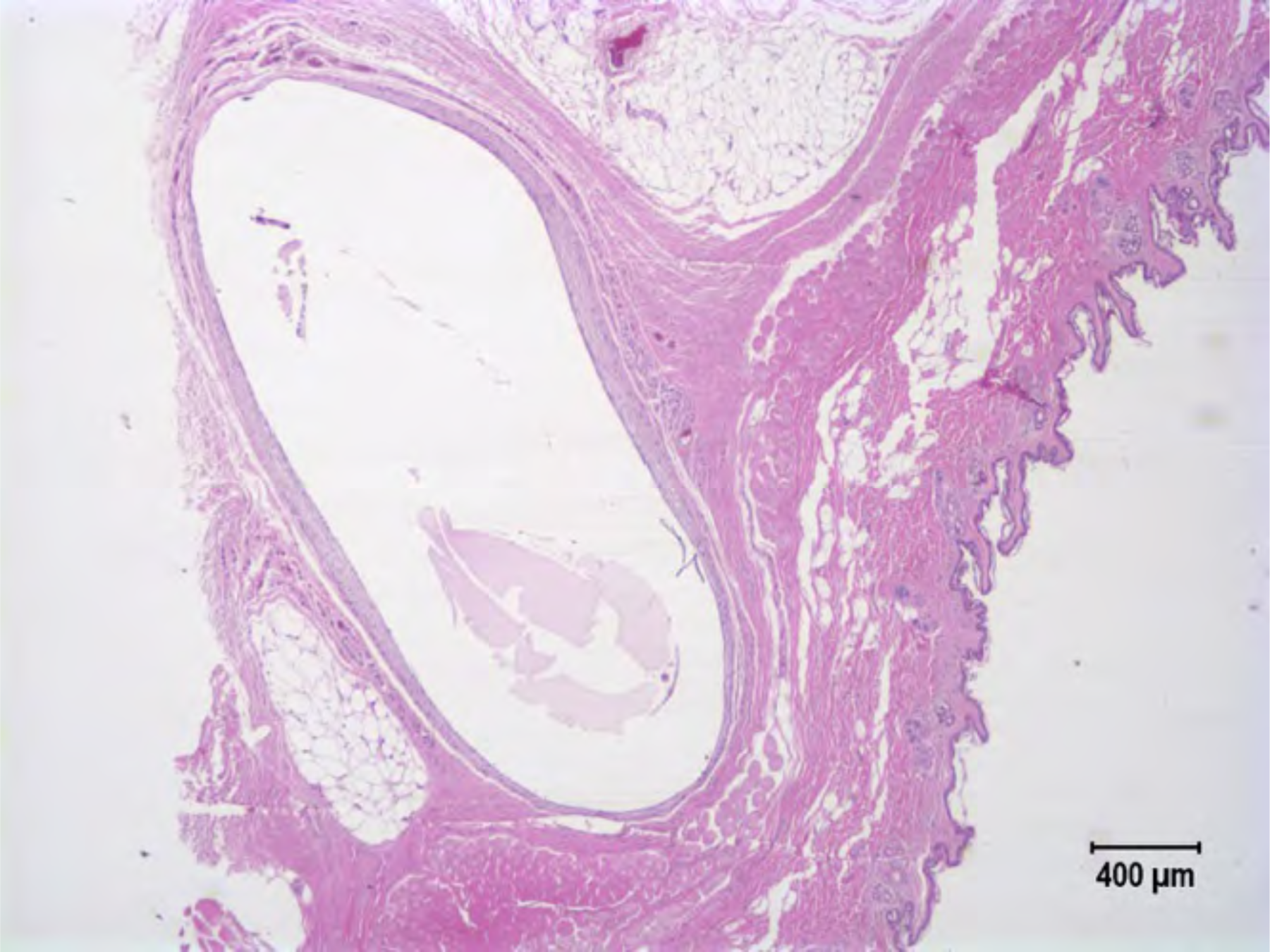
NZW Rabbit M29044. Placebo implant. Minimal inflammation detected in or around either implant. There is a thin fibrous tissue capsule around implants, but no inflammation observed in any of 8 sections. No significant lesions (NSL) are observed in the kidney, liver, spleen, lung, vagina or rectum.

**Figure S6.**
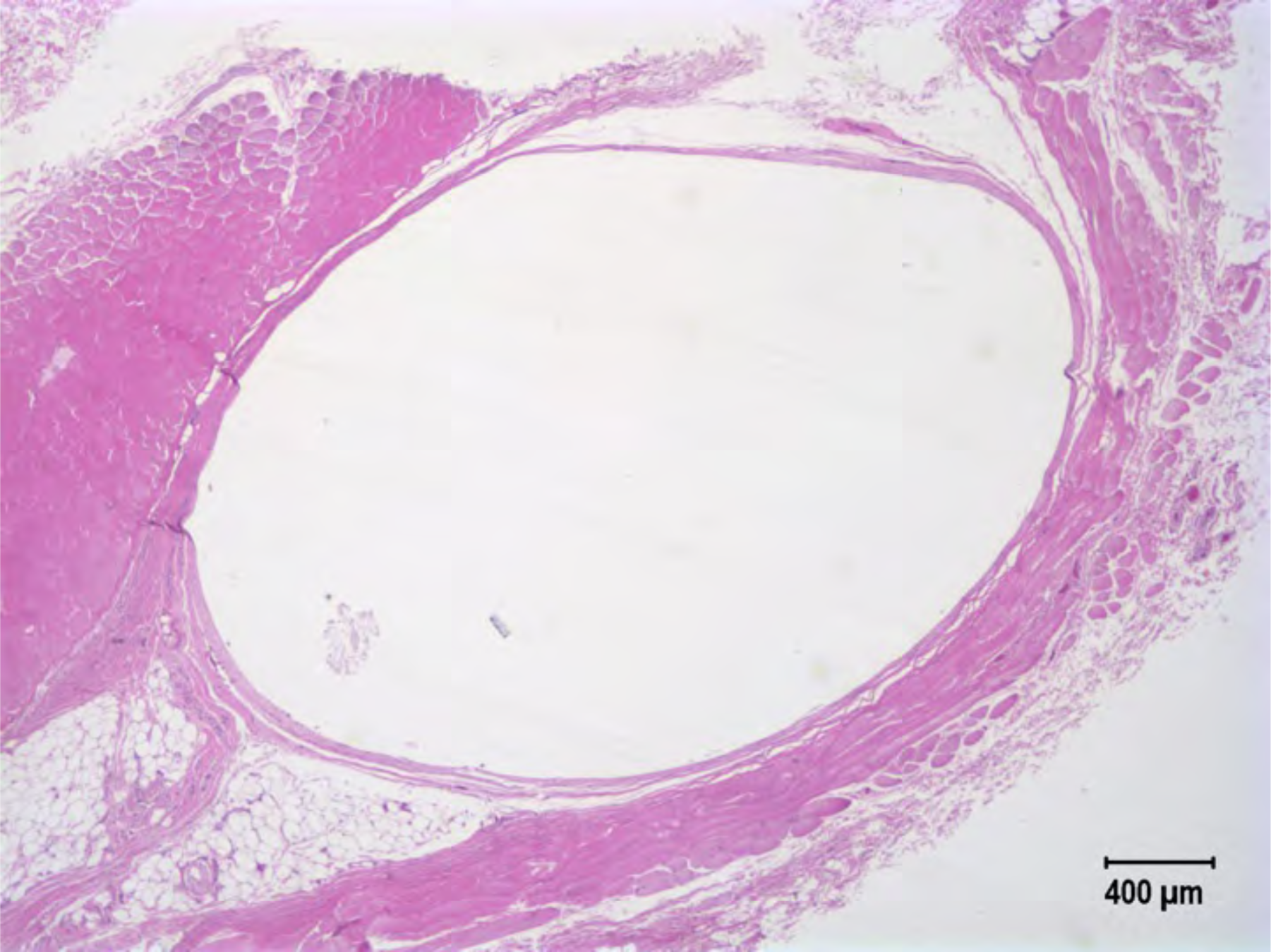
NZW Rabbit M29045. Placebo implant. Minimal inflammation in either implant. Thin fibrous tissue capsule around implants but no inflammation in any of 8 sections. NSL in kidney, liver, spleen, lung, vagina or rectum.

### Group 1: *In vitro* release rate 0.13 mg/day

**Table S4.** Histological characteristic scores from rabbits M29047 – M29051. Each rabbit received a 0.8 cm active implant (right) and a contralateral placebo (left).

|  | Response |  |  |  |  |  |  |  |
| --- | --- | --- | --- | --- | --- | --- | --- | --- |
| Animal | M29047 |  | M29048 | M29049 |  | M29050 | M29051 |  |
| Implant | Left | Right | Left | Left | Right | Left | Left | Right |
| Polymorphonuclear cells | 0 | 2 | 1 | 0 | 0 | 0 | 0 | 2 |
| Lymphocytes | 0 | 3 | 3 | 0 | 4 | 0 | 3 | 3 |
| Plasma cells | 0 | 3 | 3 | 0 | 3 | 0 | 1 | 3 |
| Macrophages | 1 | 3 | 3 | 0 | 3 | 0 | 1 | 3 |
| Giant cells | 0 | 0 | 0 | 0 | 1 | 0 | 0 | 1 |
| Necrosis | 0 | 1 | 3 | 0 | 3 | 0 | 0 | 2 |
| Capsule thickness | 2 | 4 | 3 | 2 | 4 | 2 | 1 | 2 |
| Tissue infiltrate | 0 | 2 | 1 | 0 | 2 | 0 | 1 | 1 |
| Other |  |  |  |  |  |  |  |  |
| **Overall total** | **3** | **18** | **17** | **2** | **18** | **2** | **7** | **17** |

**Figure S7.**
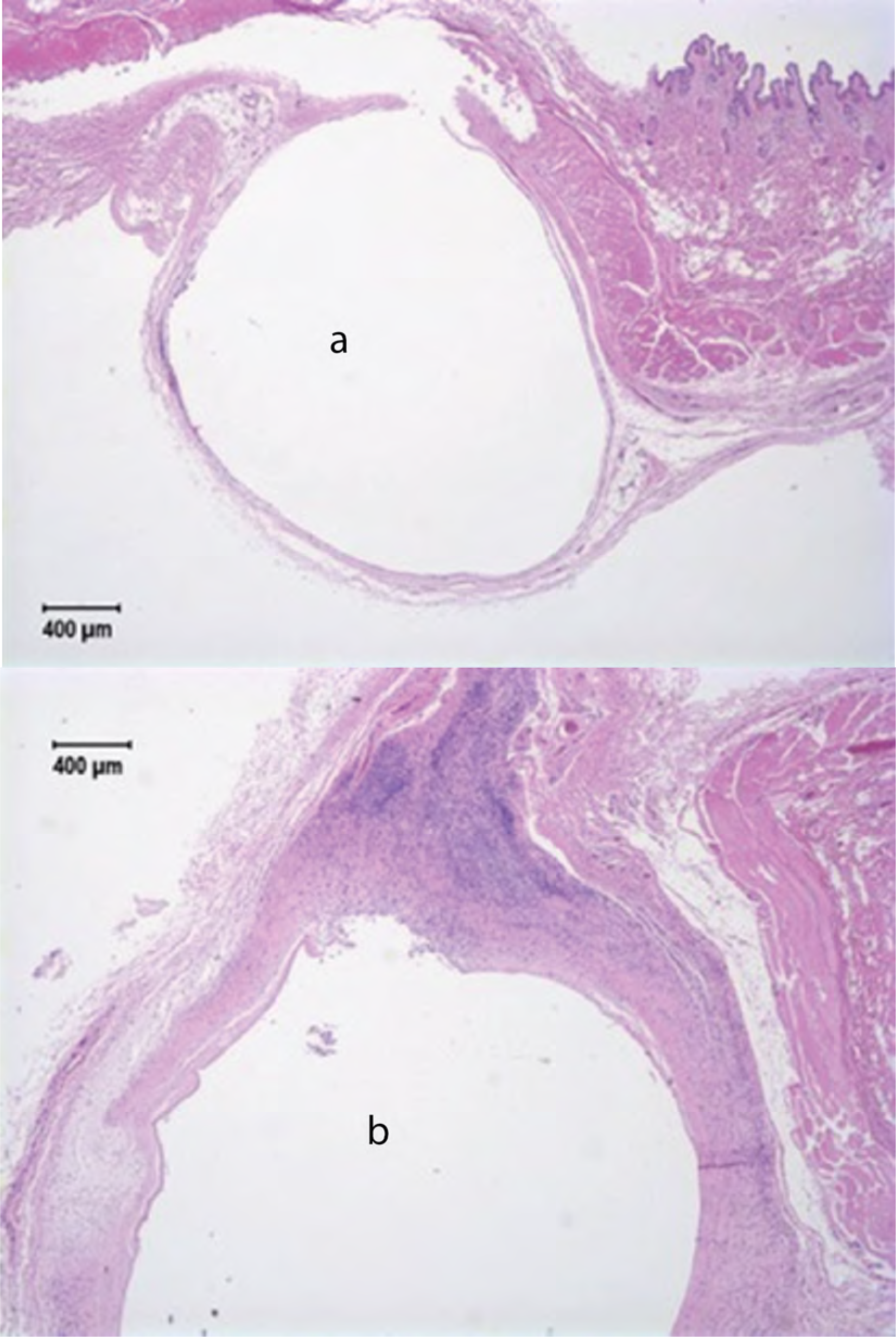
NZW Rabbit M29047. In vitro release rate 0.13 mg/day. Sections from left (a) usually have no inflammation although one section has mild to moderate inflammation. All sections from the right (b) show moderate inflammation. NSL in kidney, liver, spleen, lung, vagina or rectum.

**Figure S8.**
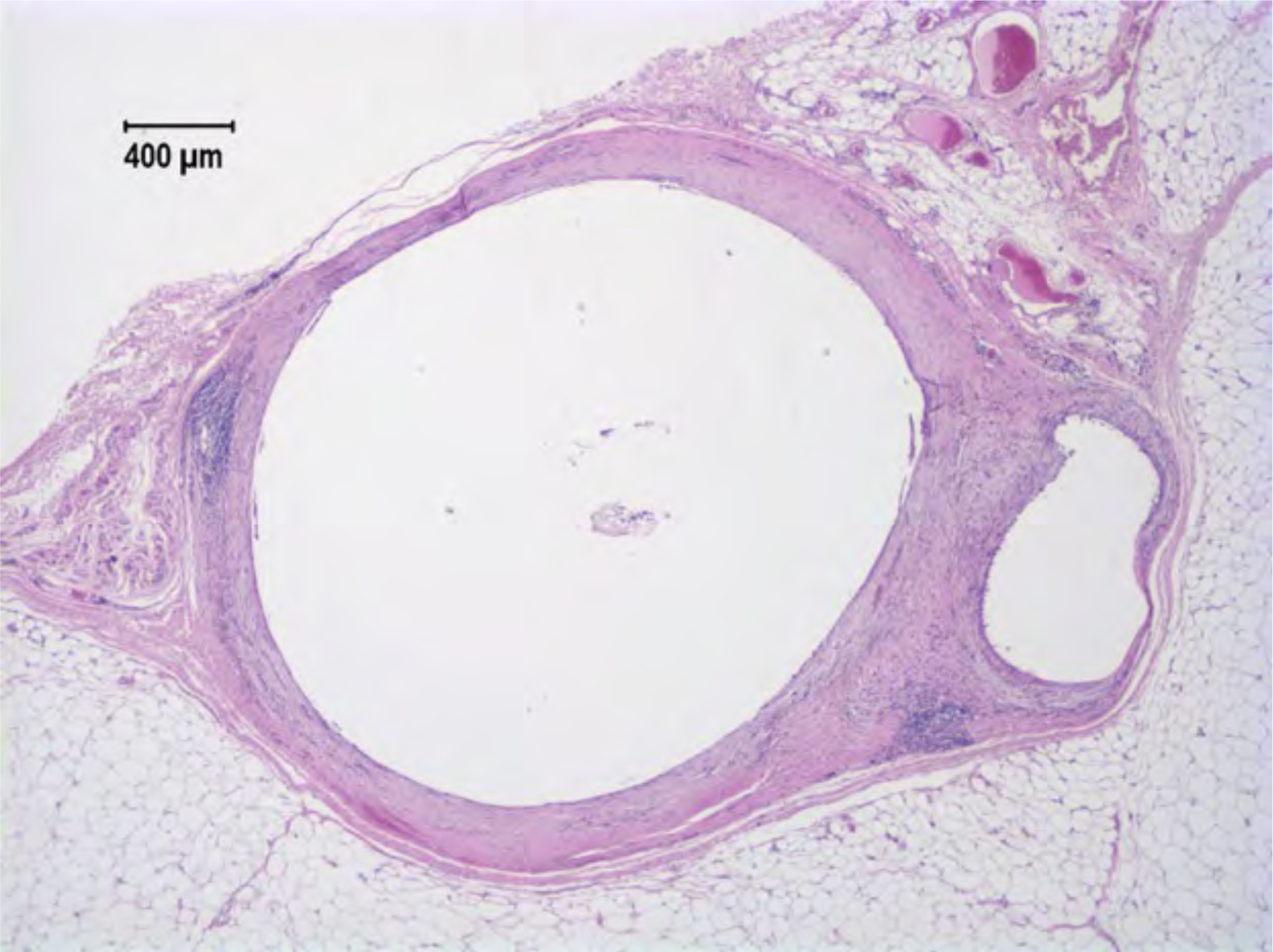
NZW Rabbit M29048. In vitro release rate 0.13 mg/day. Sections from the left implant show moderate chronic inflammation. There are NSL in kidney, liver, spleen, lung, vagina or rectum.

**Figure S9.**
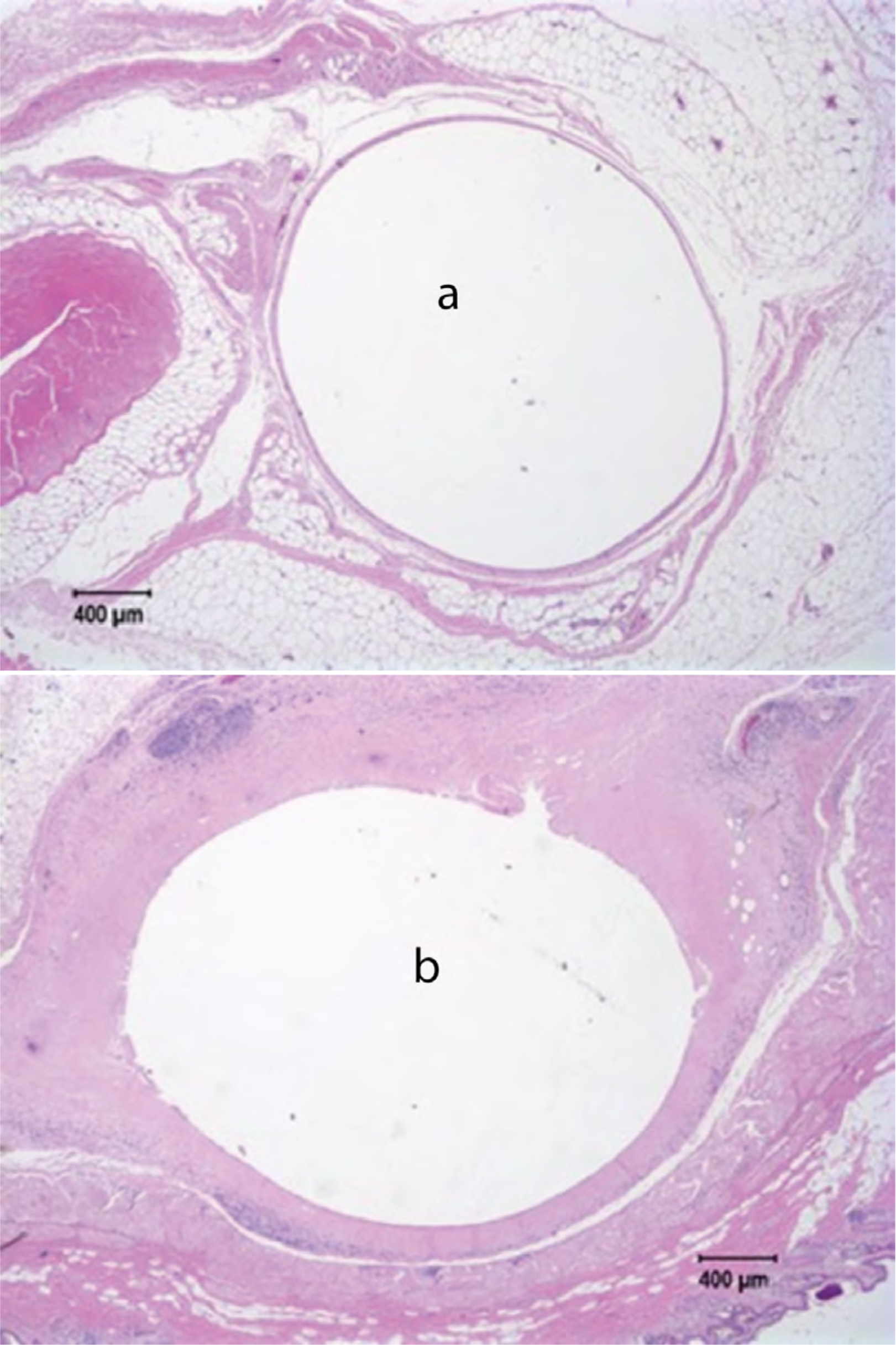
NZW Rabbit M29049. In vitro release rate 0.13 mg/day. The left implant (a) shows minimal inflammation, but the right (b) has chronic granulomatous inflammation and necrosis. There is also mild to moderate multifocal periportal lymphocytic hepatitis of unknown etiology. NSL in kidney, spleen, lung, vagina or rectum.

**Figure S10.**
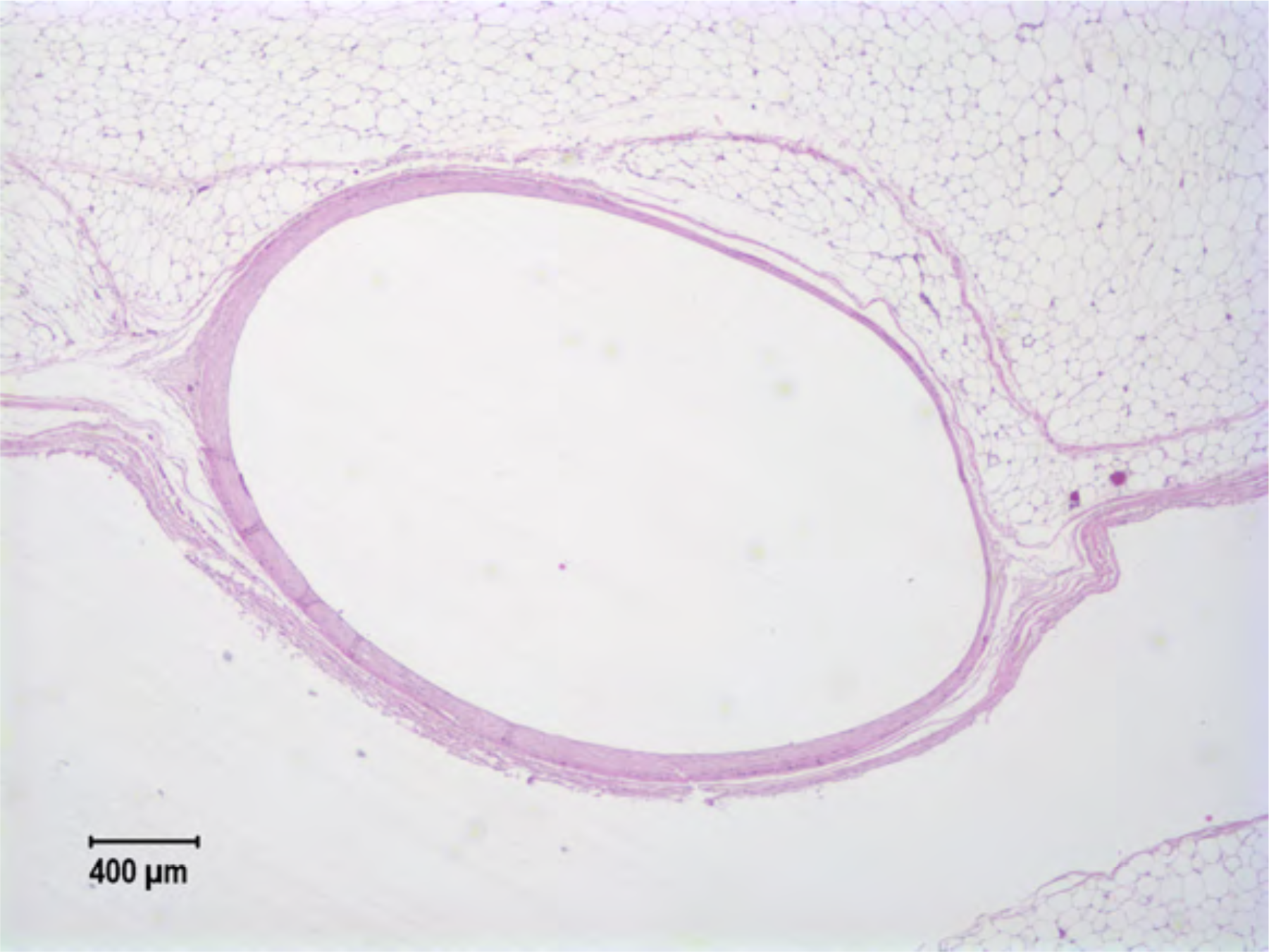
NZW Rabbit M29050. In vitro release rate 0.13 mg/day. There is minimal inflammation associated with the implant. There is mild lymphocytic periportal hepatitis of unknown etiology. NSL are observed in the kidney, spleen, lung, vagina or rectum.

**Figure S11.**
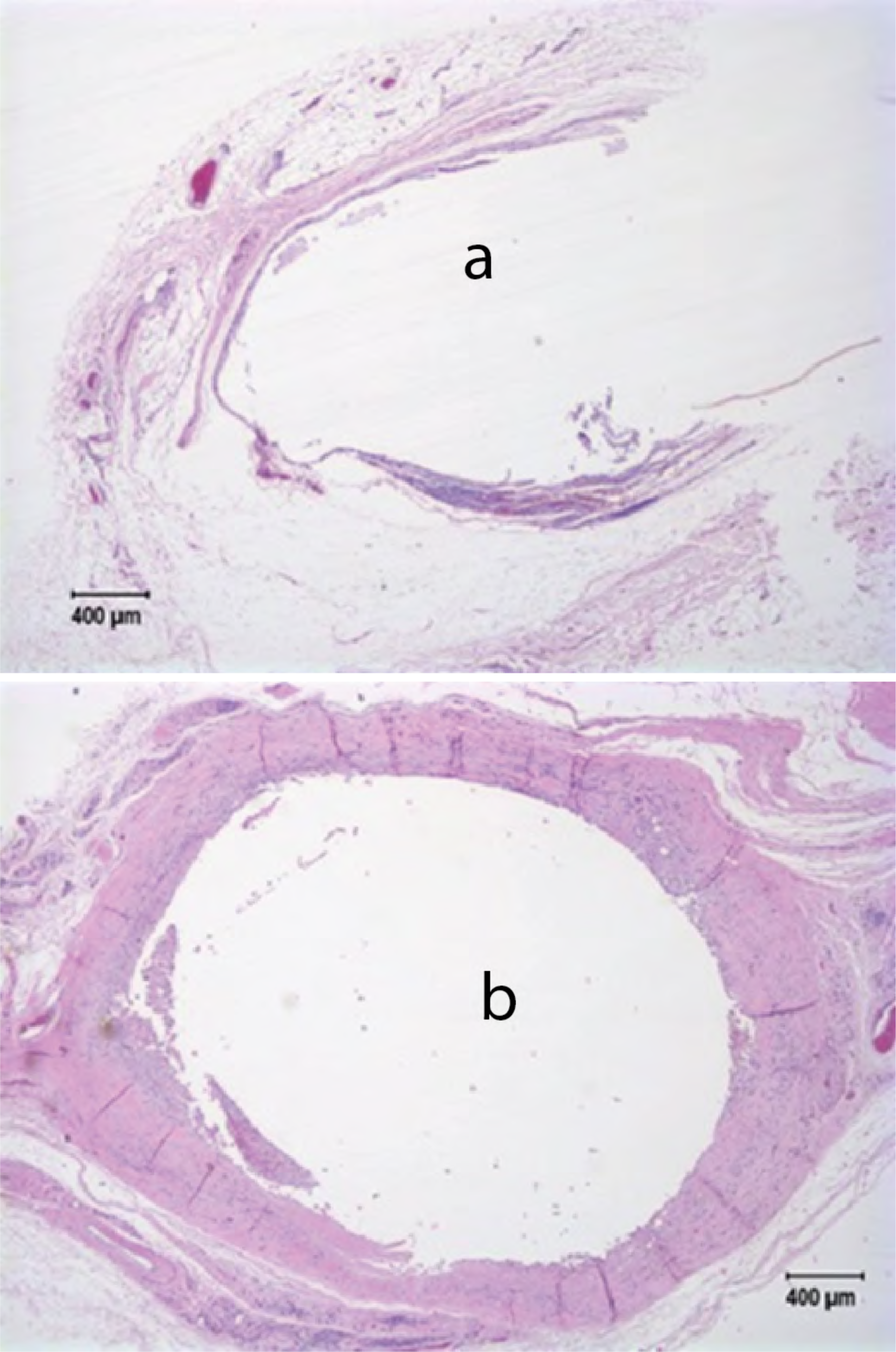
NZW rabbit M29051. In vitro release rate 0.13 mg/day. The right implant (b) has moderate to marked chronic inflammation around the implant, but the left (a) has only mild mononuclear cell infiltrations. NSL are observed in the liver, kidney, spleen, lung, vagina or rectum.

### Group 2: *In vitro* release rate 0.26 mg/day

**Table S5.** Histological characteristic scores from rabbits M29052 – M29056. Each rabbit <u>received two active 0.8 cm</u> implants.

| Animal | Response |  |  |  |  |
| --- | --- | --- | --- | --- | --- |
|  | M29052 | M29053 | M29054 | M29055 | M29056 |
| Implant | Right | Left | Left | Left | Right |
| Polymorphonuclear cells | 4 | 3 | 2 | 2 | 4 |
| Lymphocytes | 4 | 4 | 4 | 2 | 4 |
| Plasma cells | 4 | 4 | 4 | 2 | 4 |
| Macrophages | 4 | 4 | 4 | 2 | 4 |
| Giant cells | 0 | 0 | 0 | 0 | 1 |
| Necrosis | 3 | 3 | 3 | 1 | 3 |
| Capsule thickness | 3 | 4 | 3 | 3 | 3 |
| Tissue infiltrate | 0 | 3 | 0 | 2 | 3 |
| Other |  |  |  |  |  |
| **Overall total** | **22** | **25** | **20** | **14** | **26** |

**Figure S12.**
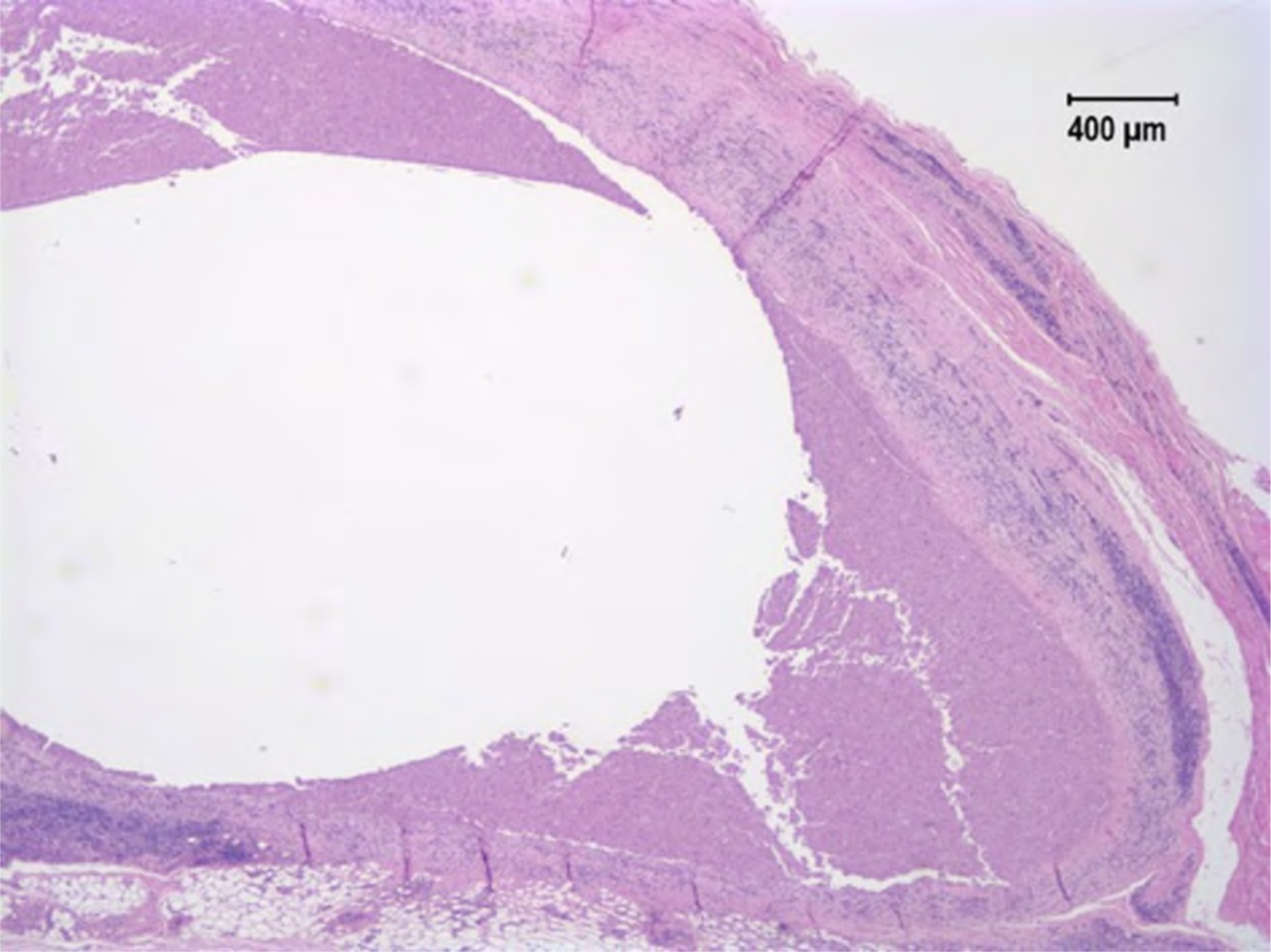
NZW Rabbit M29052. *In vitro* release rate 0.26 mg/day. All sections of the right implant have marked inflammation around the implant and necrotic cells in the center. NSL are observed in the liver, kidney, spleen, lung, vagina or rectum.

**Figure S13.**
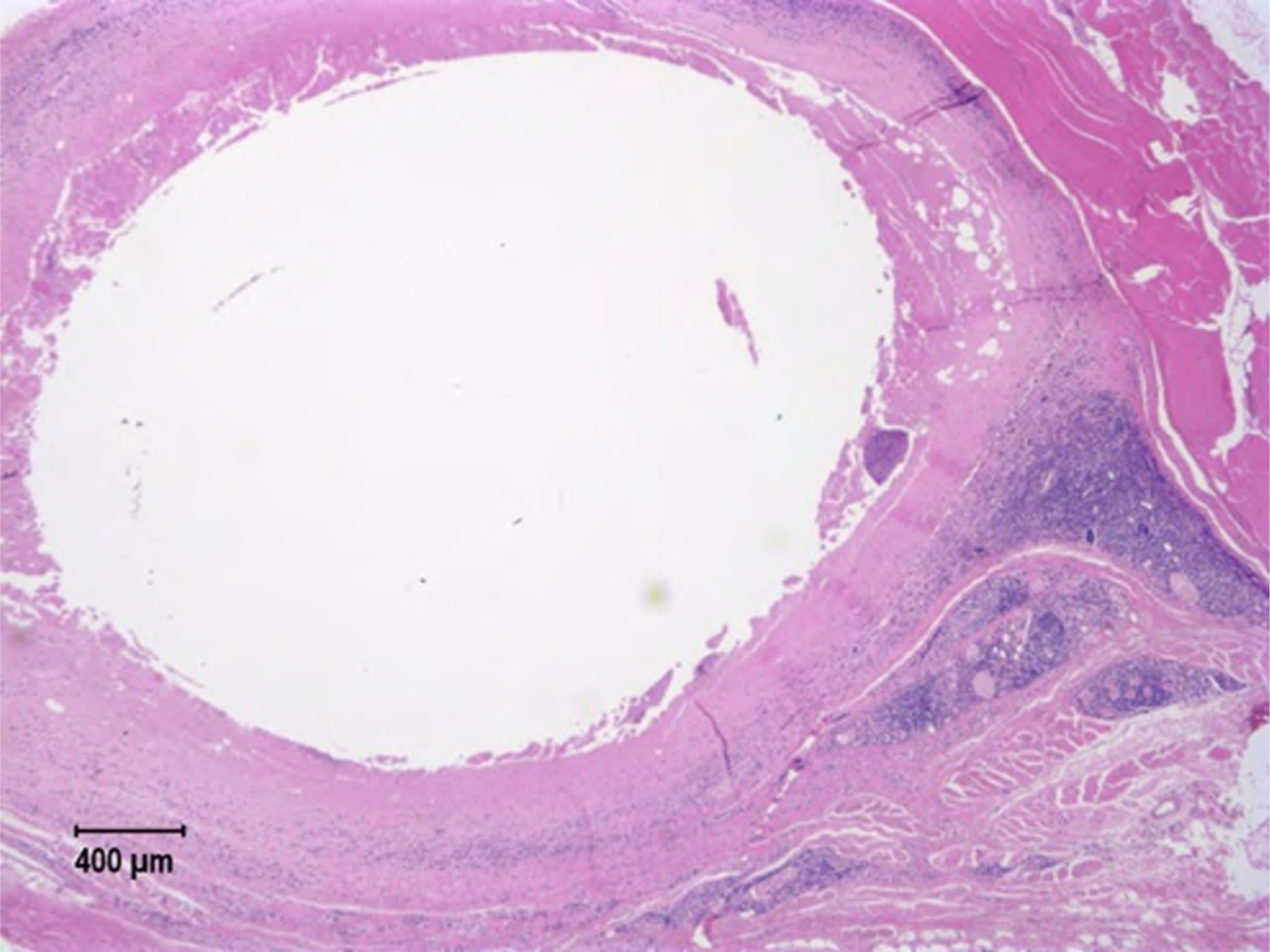
NZW Rabbit M29053. *In vitro* release rate 0.26 mg/day. All sections of the left implant have marked inflammation associated with the implant and necrotic tissue and cells in the center. NSL are observed in the liver, kidney, spleen, lung, vagina or rectum.

**Figure S14.**
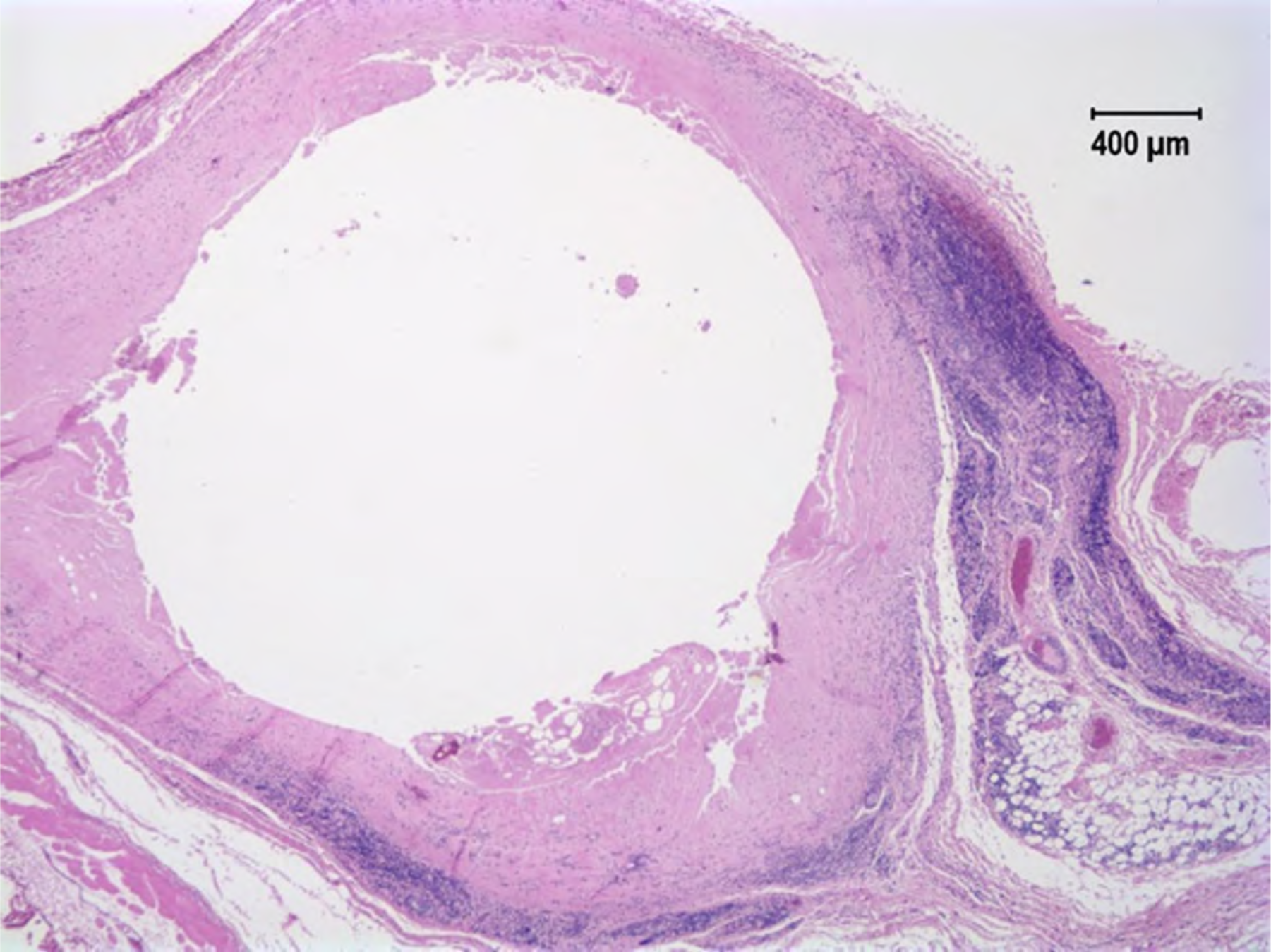
NZW Rabbit M29054. *In vitro* release rate 0.26 mg/day. All sections of the right implant have moderate to marked inflammation associated with the implant and necrotic tissue and cells in the center. NSL are observed in the liver, kidney, spleen, lung, vagina or rectum.

**Figure S15.**
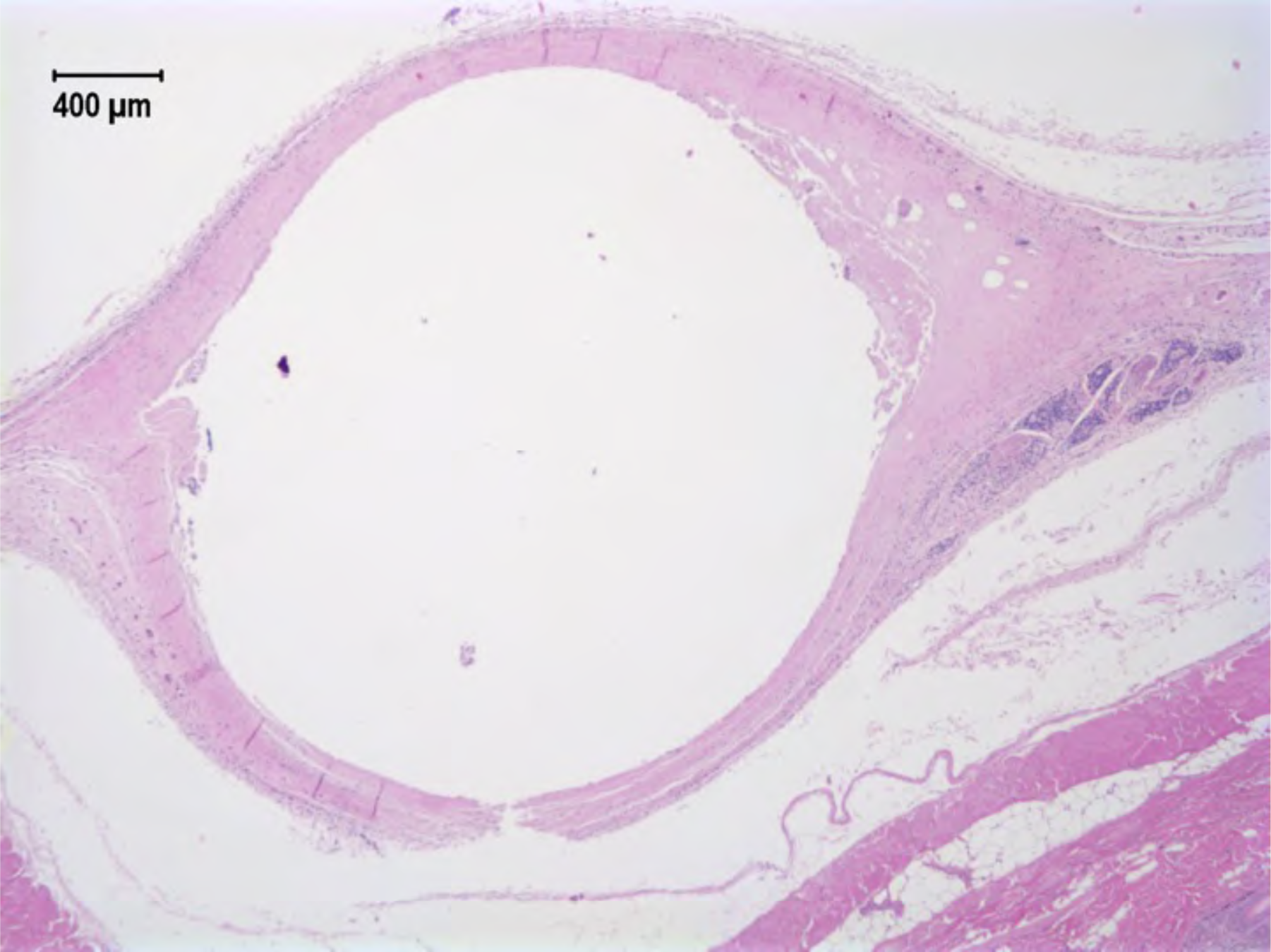
NZW Rabbit M29055. *In vitro* release rate 0.26 mg/day. There is moderate inflammation associated with the implant. There is mild lymphocytic periportal hepatitis. NSL are observed in the kidney, spleen, lung, vagina or rectum.

**Figure S16.**
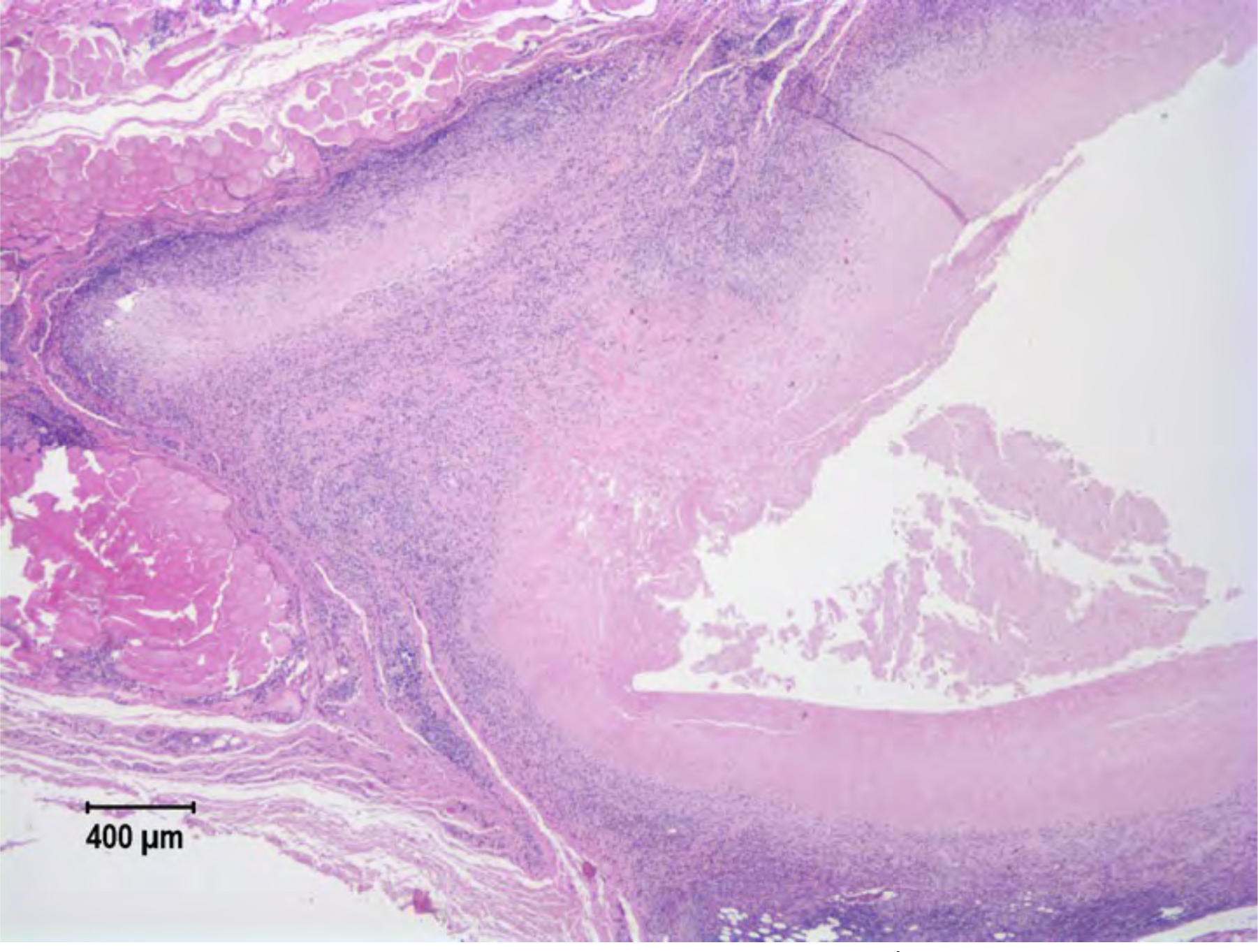
NZW Rabbit M29056. *In vitro* release rate 0.26 mg/day. There is marked inflammation and moderate necrosis associated with the implant. There is mild lymphocytic periportal hepatitis. NSL are observed in the kidney, spleen, lung, vagina or rectum.

### Group 3: *In vitro* release rate 0.48 mg/day

**Table S6.** Histological characteristic scores for animals M29057 – M29061. Each animal <u>received two active 1.6 cm implants.</u>

|  | Response |  |  |  |  |
| --- | --- | --- | --- | --- | --- |
| Animal | M29057 | M29058 | M29059 | M29060 | M29061 |
| Implant | Right | Left | Left | Left | Right |
| Polymorphonuclear cells | 2 | 4 | 4 | 4 | 4 |
| Lymphocytes | 3 | 4 | 4 | 4 | 4 |
| Plasma cells | 3 | 4 | 4 | 4 | 4 |
| Macrophages | 3 | 4 | 4 | 4 | 4 |
| Giant cells | 0 | 0 | 0 | 0 | 0 |
| Necrosis | 3 | 3 | 3 | 4 | 3 |
| Capsule thickness | 0 | 3 | 3 | 3 | 3 |
| Tissue infiltrate | 0 | 3 | 3 | 3 | 3 |
| Other |  |  |  |  |  |
| Overall total | 14 | 25 | 25 (marked inflammation) | 26 | 25 (marked inflammation) |

**Figure S17.**
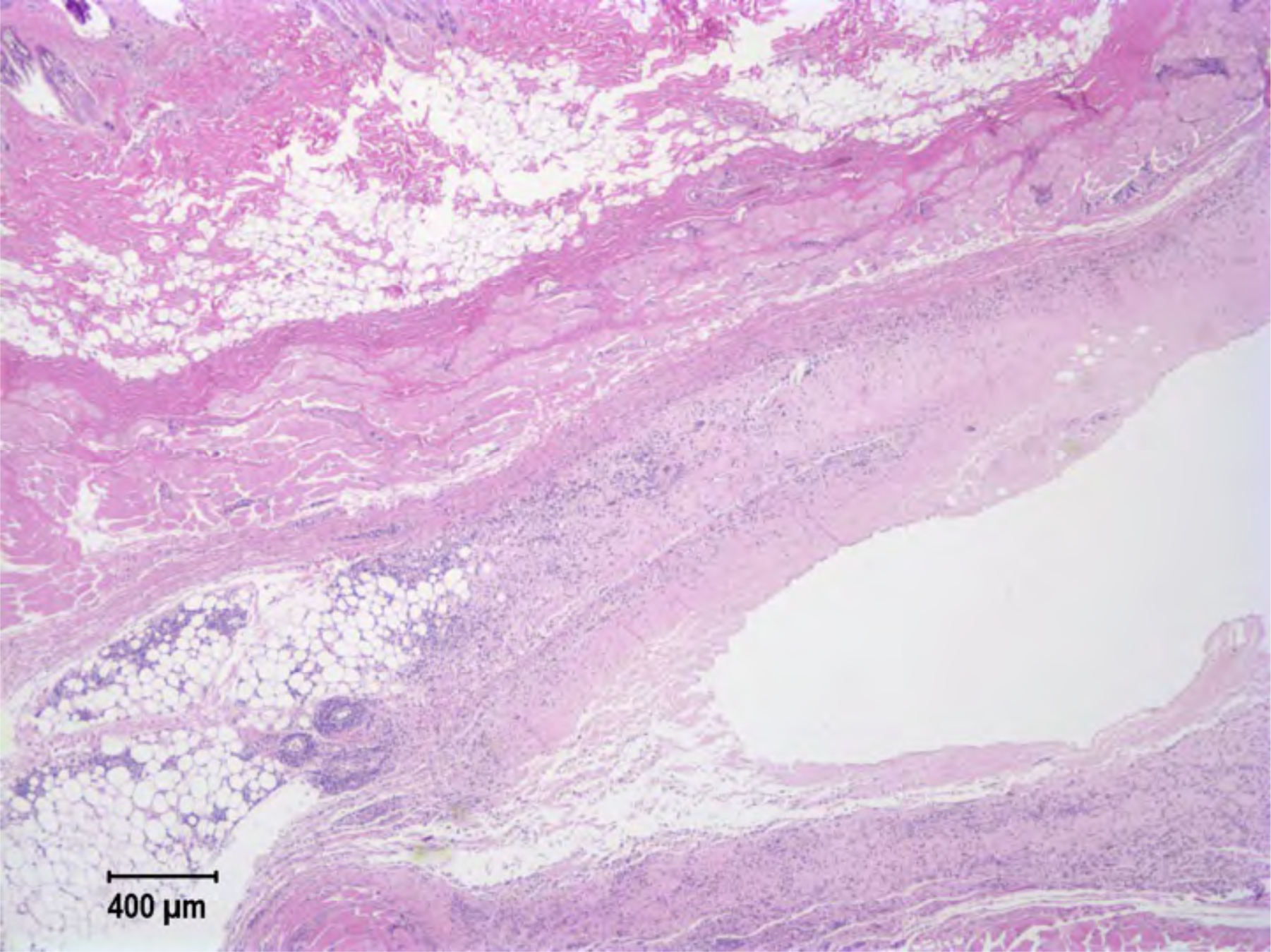
NZW Rabbit M29057. *In vitro* release rate 0.48 mg/day. There is moderate to marked inflammation and moderate necrosis associated with the implant. NSL are observed in the liver, kidney, spleen, lung, vagina or rectum

**Figure S18.**
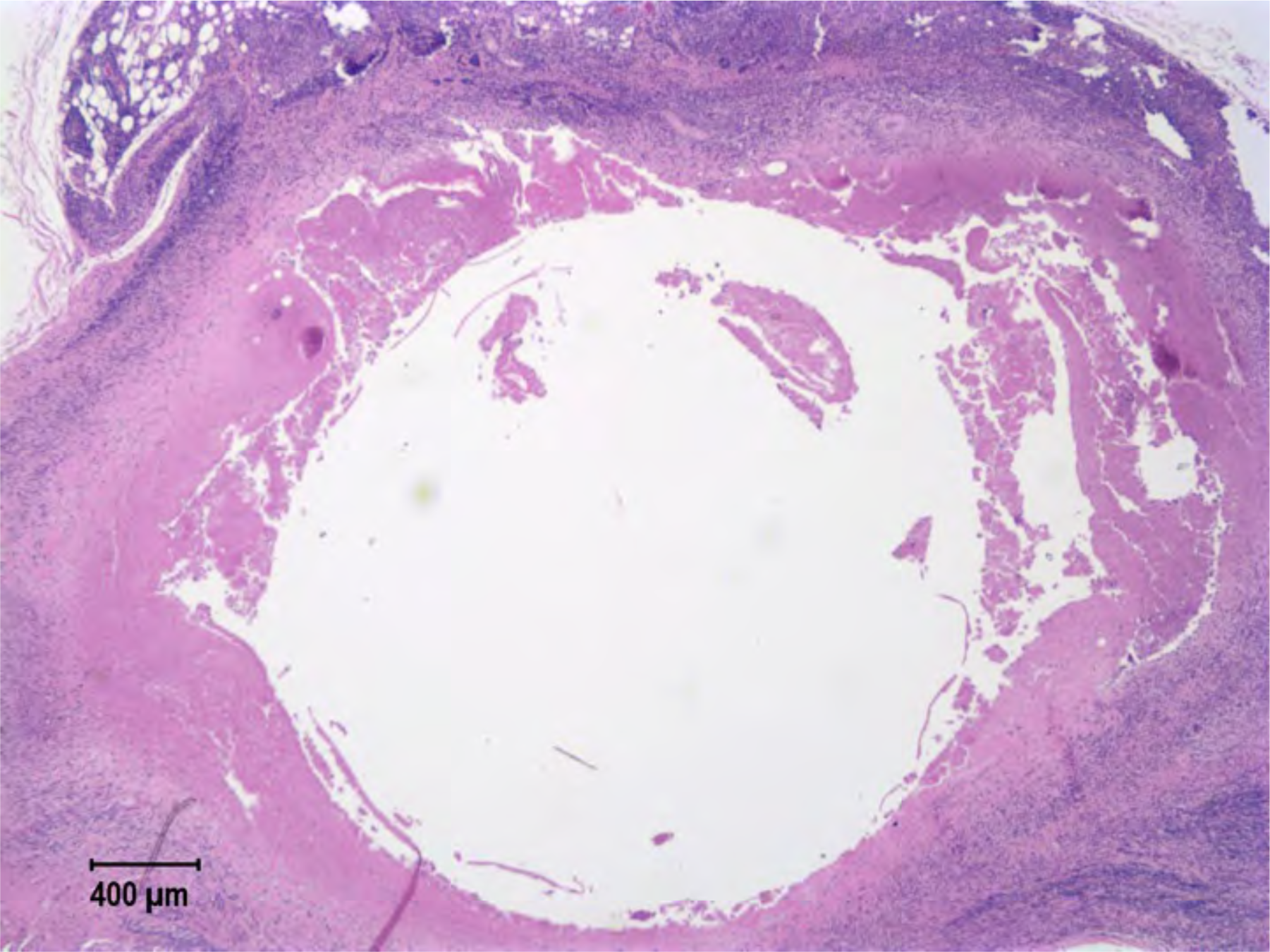
NZW Rabbit M29058. *In vitro* release rate 0.48 mg/day. All sections from the left implant show marked inflammation and moderate necrosis associated with the implant. NSL are observed in the liver, kidney, spleen, lung, vagina or rectum

**Figure S19.**
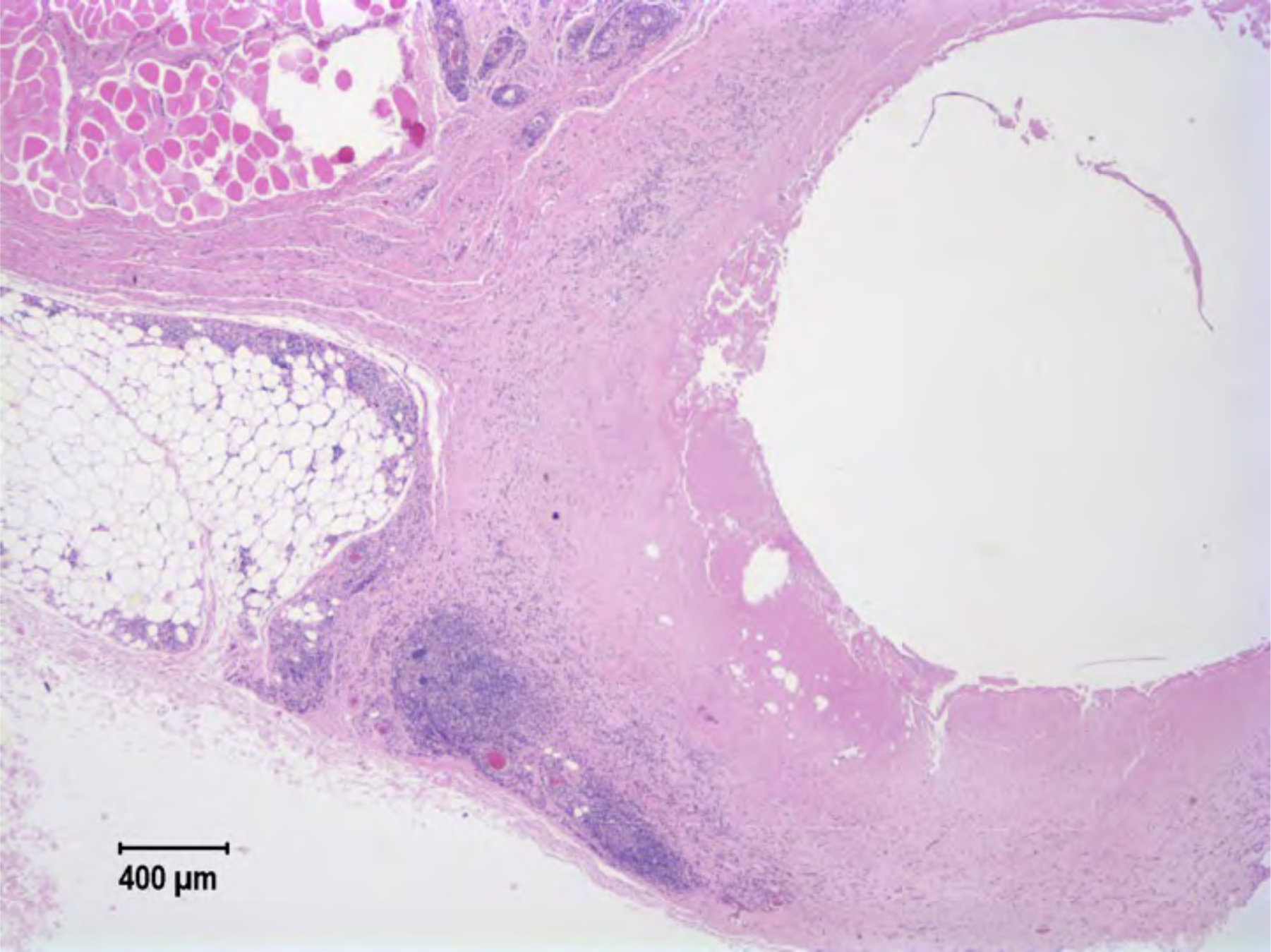
NZW Rabbit M29059. *In vitro* release rate 0.48 mg/day. All sections from the right implant show marked inflammation and moderate necrosis associated with the implant. NSL are observed in the liver, kidney, spleen, lung, vagina or rectum

**Figure S20.**
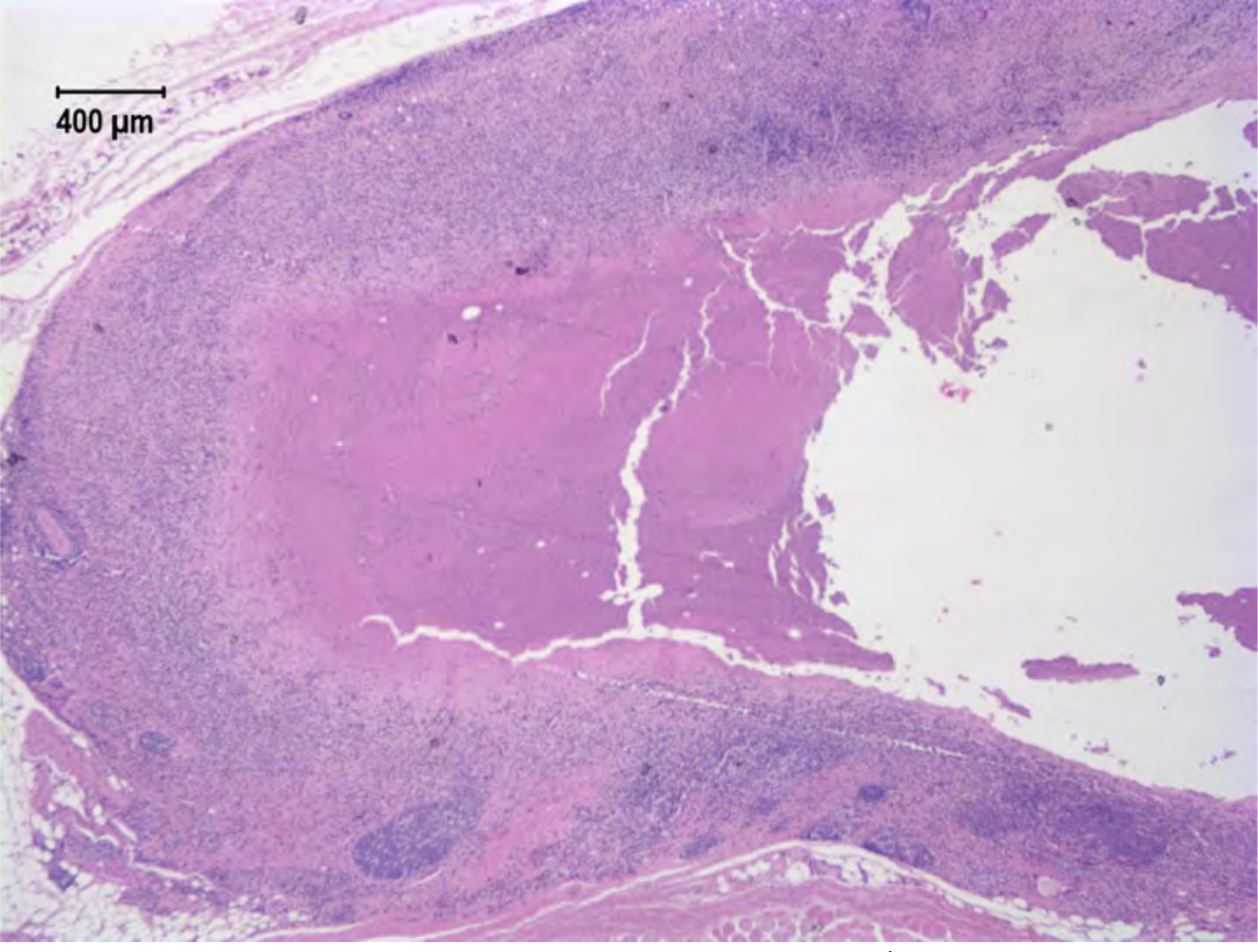
NZW Rabbit M29060. *In vitro* release rate 0.48 mg/day. All sections from the left implant show marked inflammation and moderate necrosis associated with the implant. NSL are observed in the liver, kidney, spleen, lung, vagina or rectum

**Figure S21.**
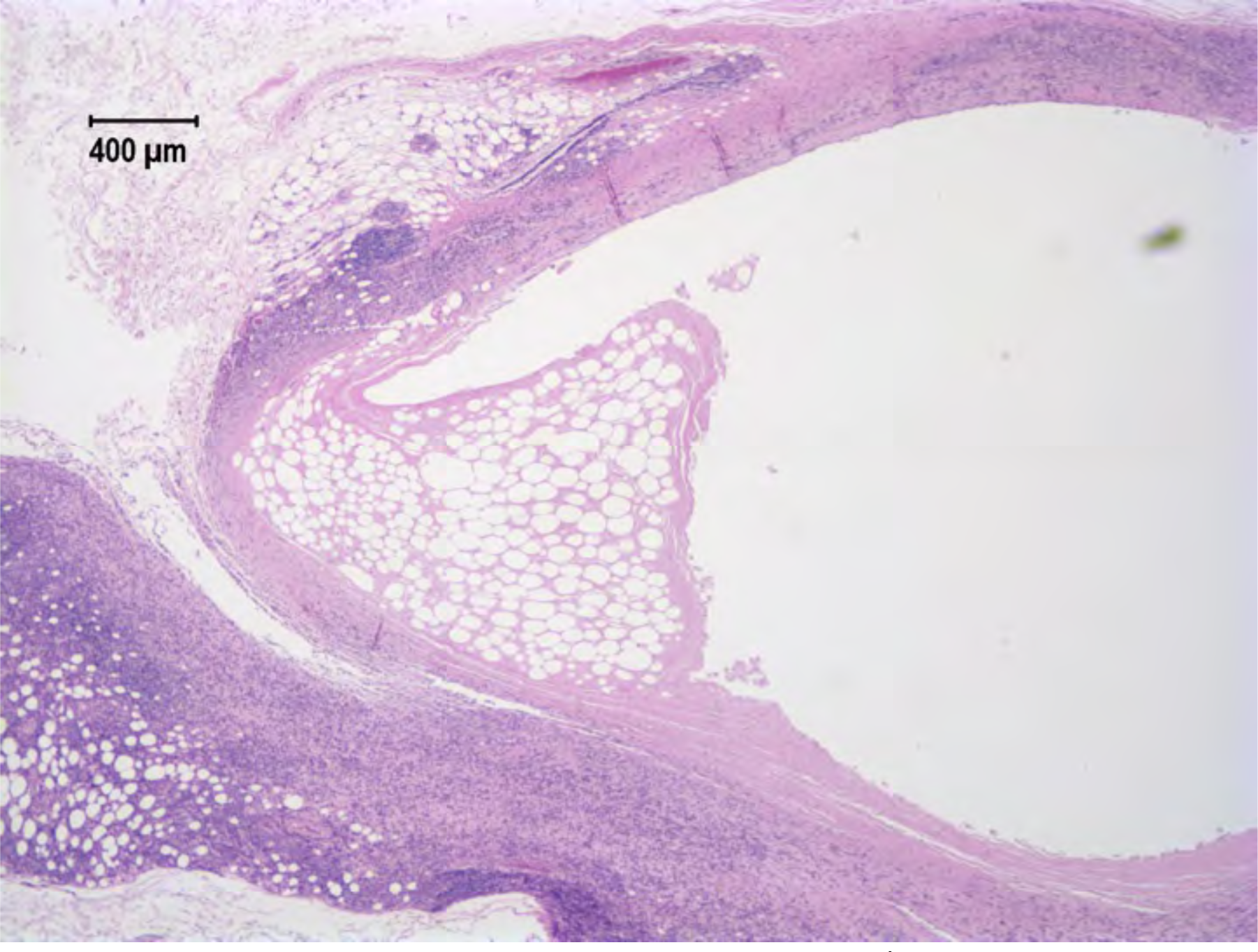
NZW Rabbit M29061. *In vitro* release rate 0.48 mg/day. All sections from the left implant show marked inflammation and moderate necrosis associated with the implant. NSL are observed in the liver, kidney, spleen, lung, vagina or rectum

### Group 4: *In vitro* release rate of 0.72 mg/day

**Table S7.**
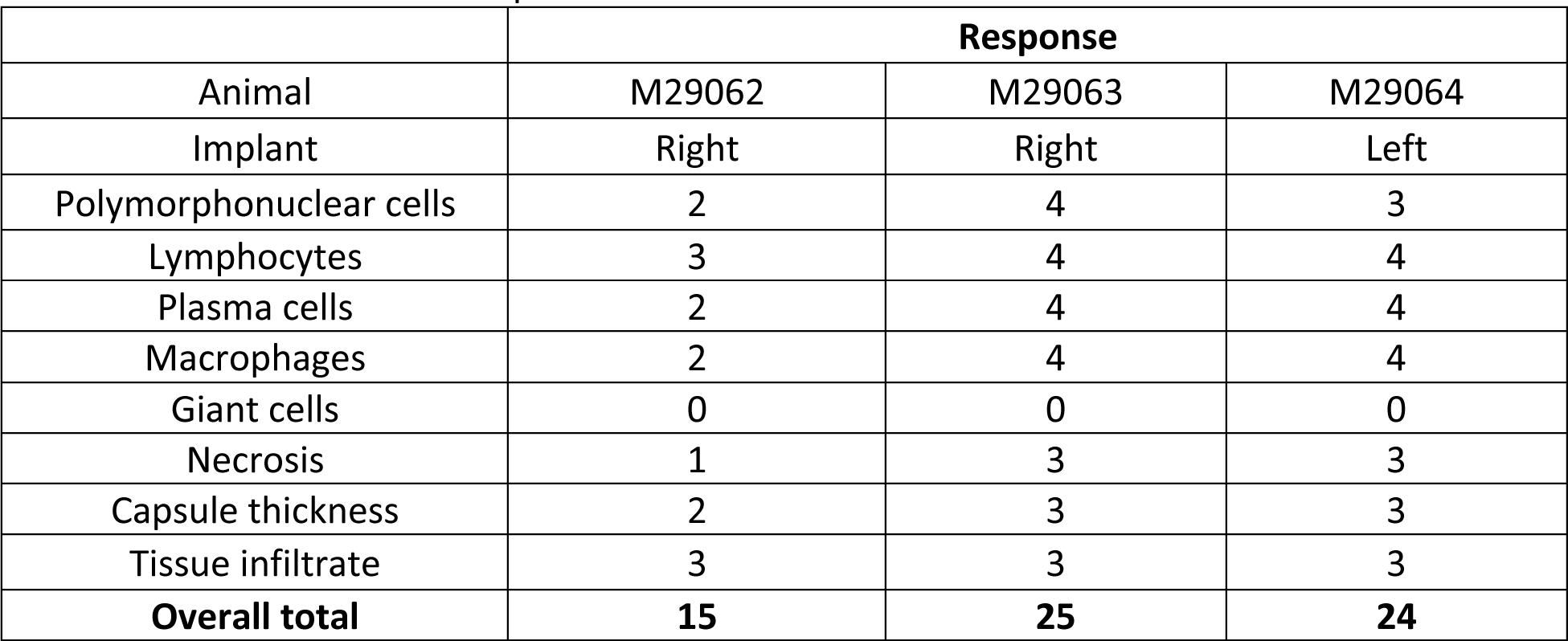
Histological characteristic scores for animals M29062 – M29064. Each animal received three active 1.6 cm implants.

| Animal | Response |  |  |
| --- | --- | --- | --- |
|  | M29062 | M29063 | M29064 |
| Implant | Right | Right | Left |
| Polymorphonuclear cells | 2 | 4 | 3 |
| Lymphocytes | 3 | 4 | 4 |
| Plasma cells | 2 | 4 | 4 |
| Macrophages | 2 | 4 | 4 |
| Giant cells | 0 | 0 | 0 |
| Necrosis | 1 | 3 | 3 |
| Capsule thickness | 2 | 3 | 3 |
| Tissue infiltrate | 3 | 3 | 3 |
| **Overall total** | **15** | **25** | **24** |

**Figure S22.**
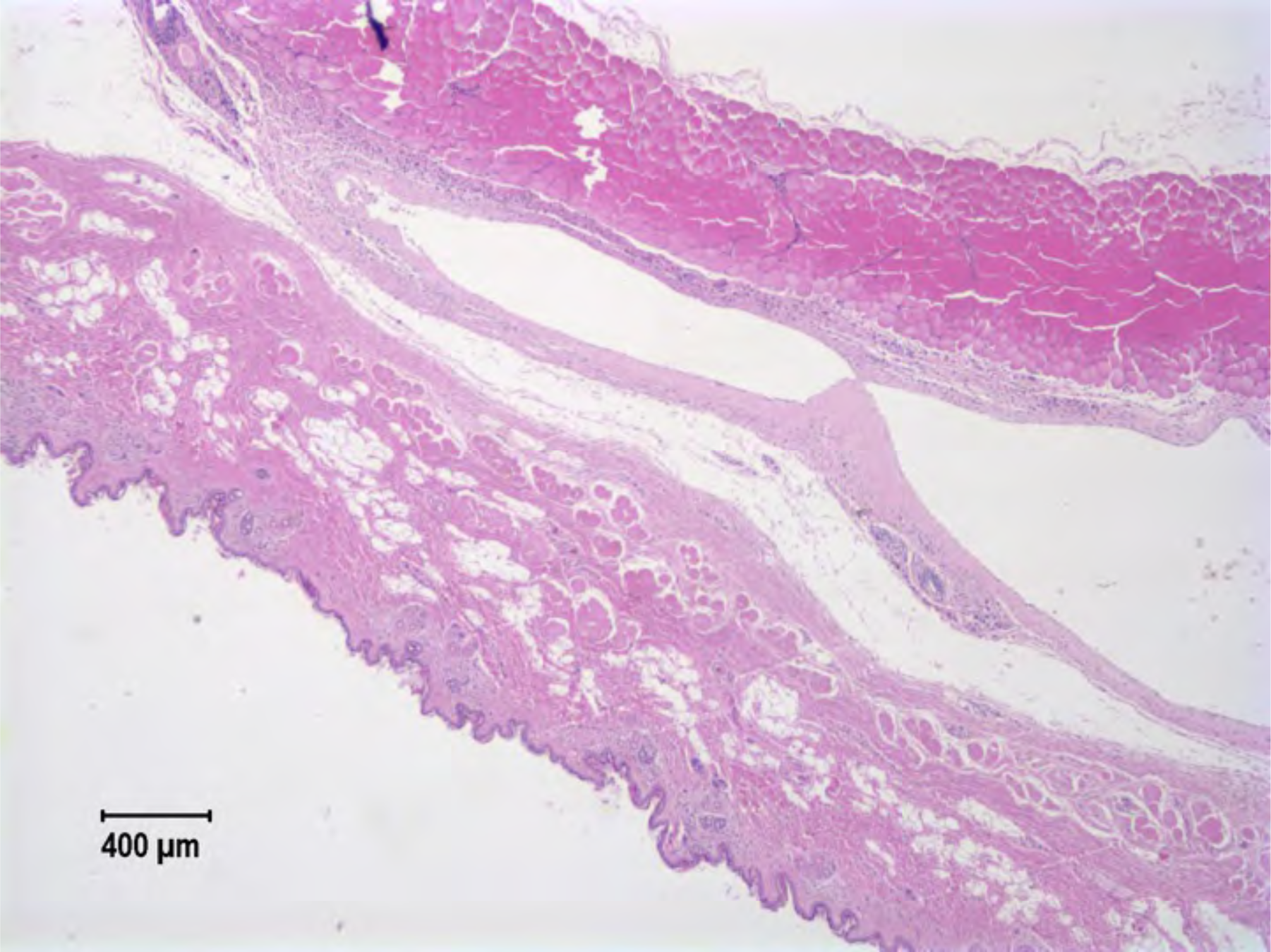
NZW Rabbit M29062. *In vitro* release rate 0.72 mg/day. All sections from the left implant show marked inflammation and moderate necrosis associated with the implant. NSL are observed in the liver, kidney, spleen, lung, vagina or rectum

**Figure S23.**
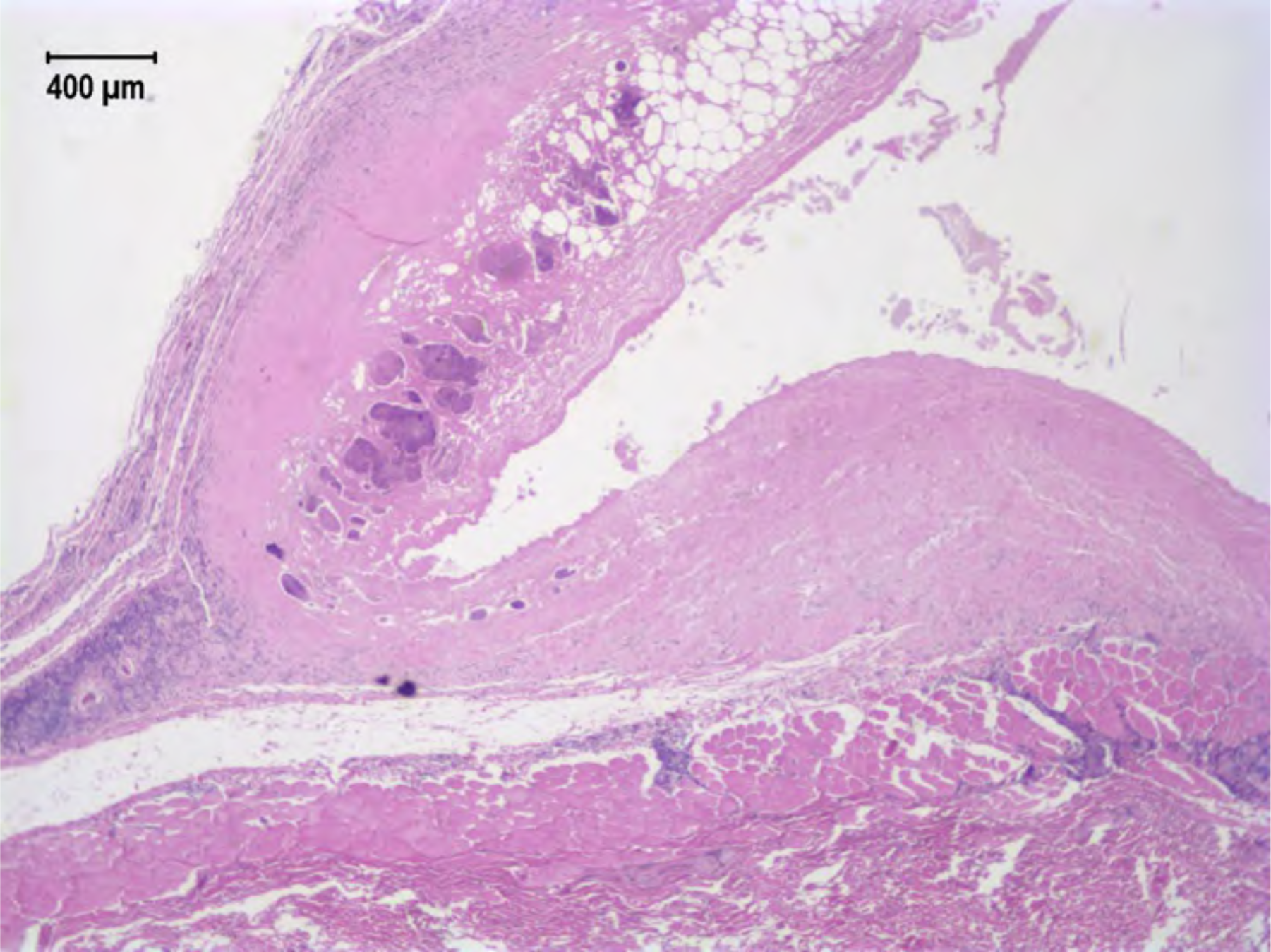
NZW Rabbit M29063. *In vitro* release rate 0.72 mg/day. All sections from the right implant show marked inflammation and moderate necrosis associated with the implant. NSL are observed in the liver, kidney, spleen, lung, vagina or rectum

**Figure S24.**
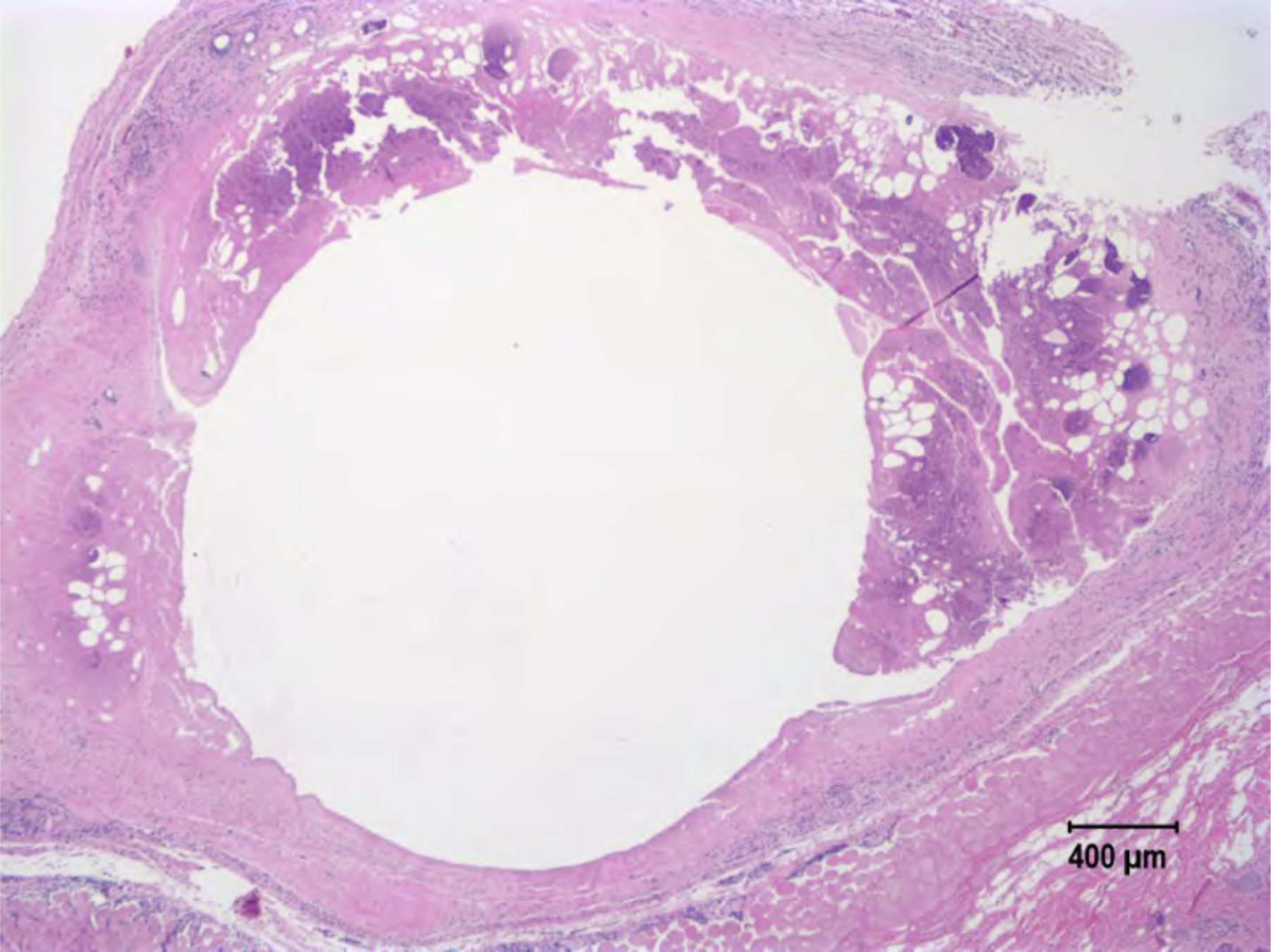
NZW Rabbit M29064. *In vitro* release rate 0.72 mg/day. All sections from the left implant show marked inflammation and moderate necrosis associated with the implant. NSL are observed in the liver, kidney, spleen, lung, vagina or rectum.

## Supplemental 4. 4 week and 12-week histology report of TAF generation B in rhesus macaques

Note: Each rhesus macaque received two 2 cm implants: one placebo (right), and one active (left).

**Four-week histology reports:**

**Table S8.** Histological characteristic scores for rhesus macaque DP15. Implant placed 2/22/2018; euthanized 3/22/2018.

| Animal# DP15<br>Necropsy# 18A100 | Response |  |
| --- | --- | --- |
|  | Left (Active) (1-4) | Right (Placebo) (1-4) |
| Implant |  |  |
| Polymorphonuclear cells | 3 | 1 |
| Lymphocytes | 3 | 1 |
| Plasma cells | 3 | 1 |
| Macrophages | 3 | 2 |
| Giant cells | 2* | 0 |
| Necrosis | 2 | 0 |
| Capsule thickness | 3 | 2 |
| Tissue infiltrate | 3 | 1 |
| Other |  |  |
| **Overall total** | **22** | **8** |

\*NB: There are significant multifocal areas of granulomatous inflammation associated with suture material which often coalesces with the implant margins, and most multinucleated giant cells (MNGC) appear to be associated with these rather than the implant.

**Figure S25.**
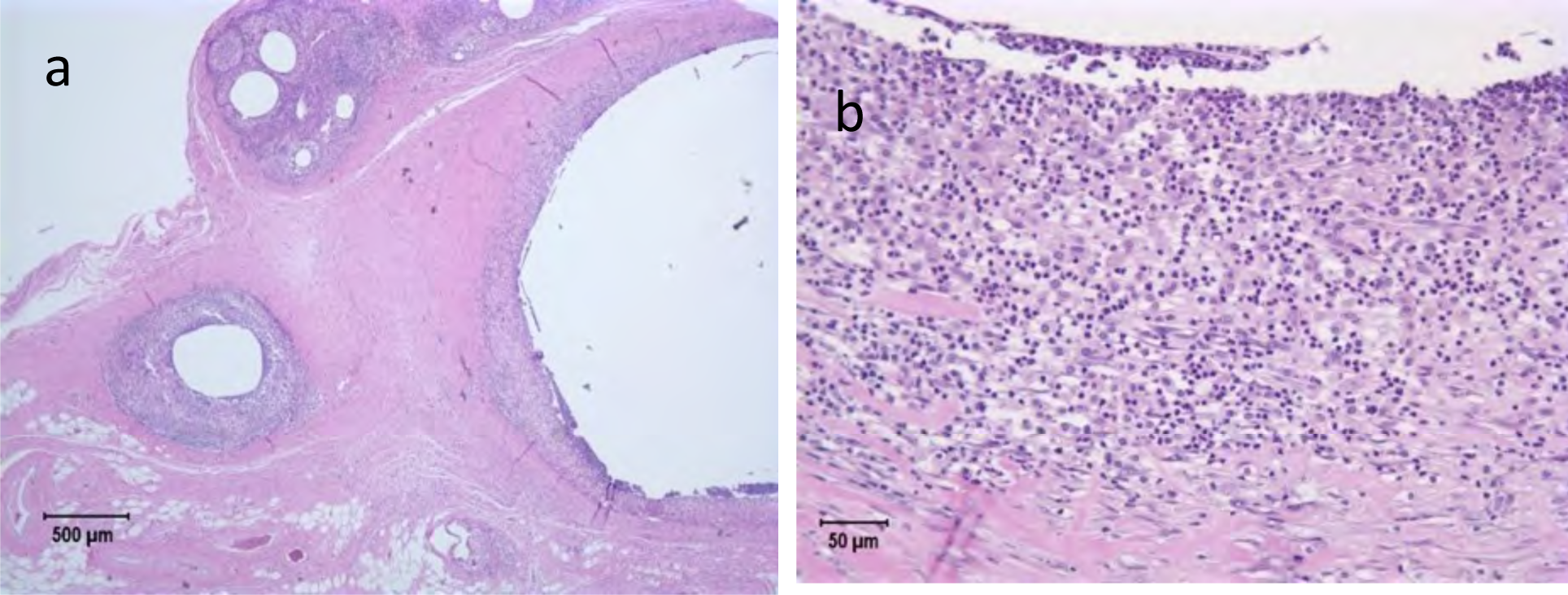
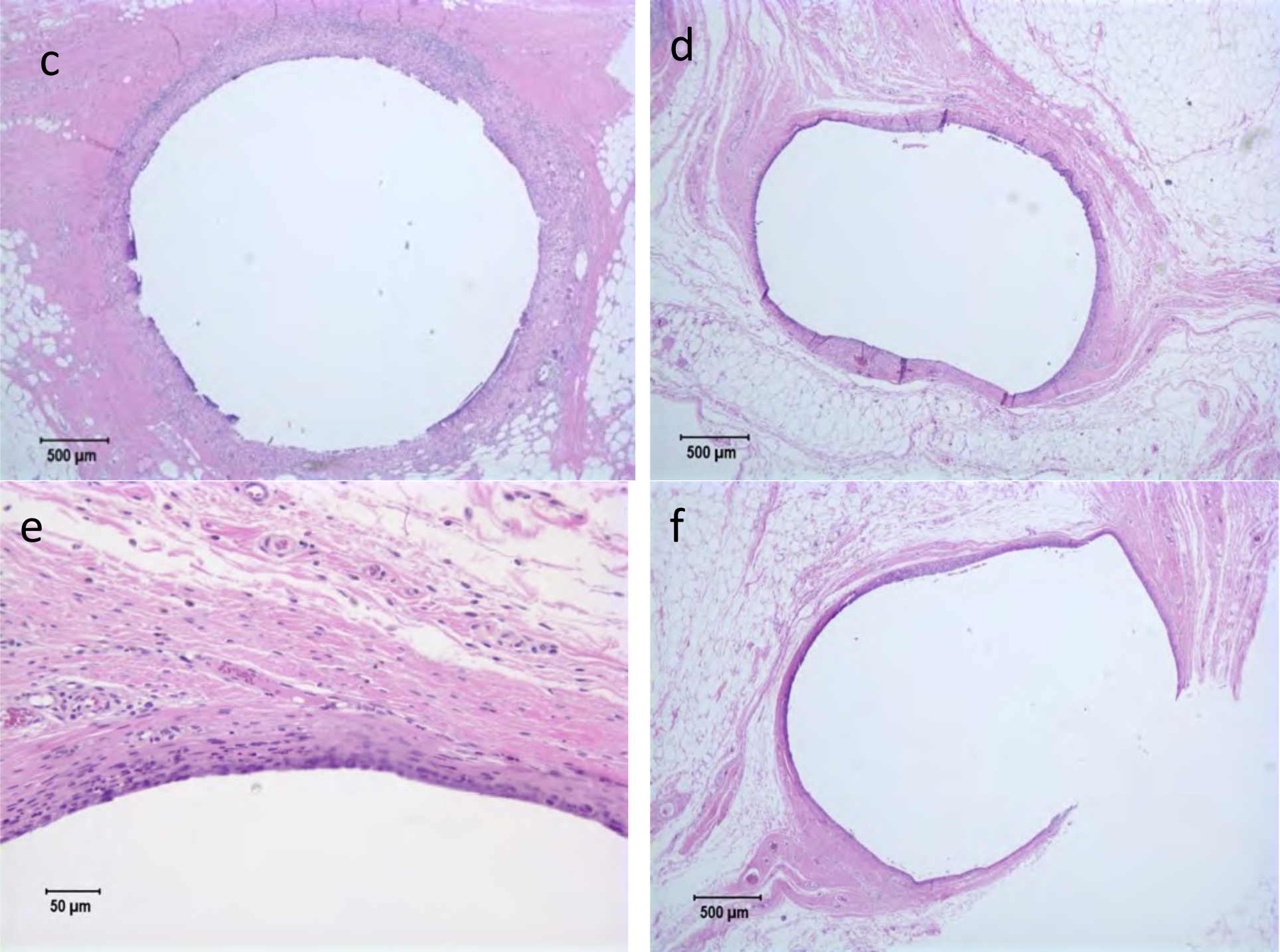
Rhesus macaque DP15. *In vitro* release rate 0.13 mg/day. The left (active) implant (a-c) shows a moderately thick band of heterophils, macrophages, lymphocytes, and plasma cells surrounding implant interspersed with moderate numbers of multinucleated giant cells in peripheral sections. Fibrous tissue and inflammation extend into adjacent connective tissue. The right (placebo) implant (d-f) clearly has less inflammation, yet adjacent connective tissues of both implants have some fibrosis and minimal inflammation. *NB: There are significant multifocal areas of granulomatous inflammation associated with suture material which often coalesces with the implant margins, and most MNGC appear to be associated with these rather than the implant.

**Table S9.** Histological characteristic scores for rhesus macaque GH84. Implant placed 2/22/2018; euthanized 3/22/2018.

| Animal# GH84<br>Necropsy# 18A101 | Response |  |
| --- | --- | --- |
|  | Left (Active) 1-4 | Right (Placebo) 1-4 |
| Implant |  |  |
| Polymorphonuclear cells | 3 | 0 |
| Lymphocytes | 3 | 1 |
| Plasma cells | 2 | 1 |
| Macrophages | 2 | 1 |
| Giant cells | 1* | 0 |
| Necrosis | 1 | 0 |
| Capsule thickness | 3 | 1 |
| Tissue infiltrate | 2 | 1 |
| Other |  |  |
| Overall total | 17 | 5 |

**\*NB:** There are significant multifocal areas of granulomatous inflammation associated with suture material which often coalesces with the implant margins, and most MNGC appear to be associated with these rather than the implant.

**Figure S26.**
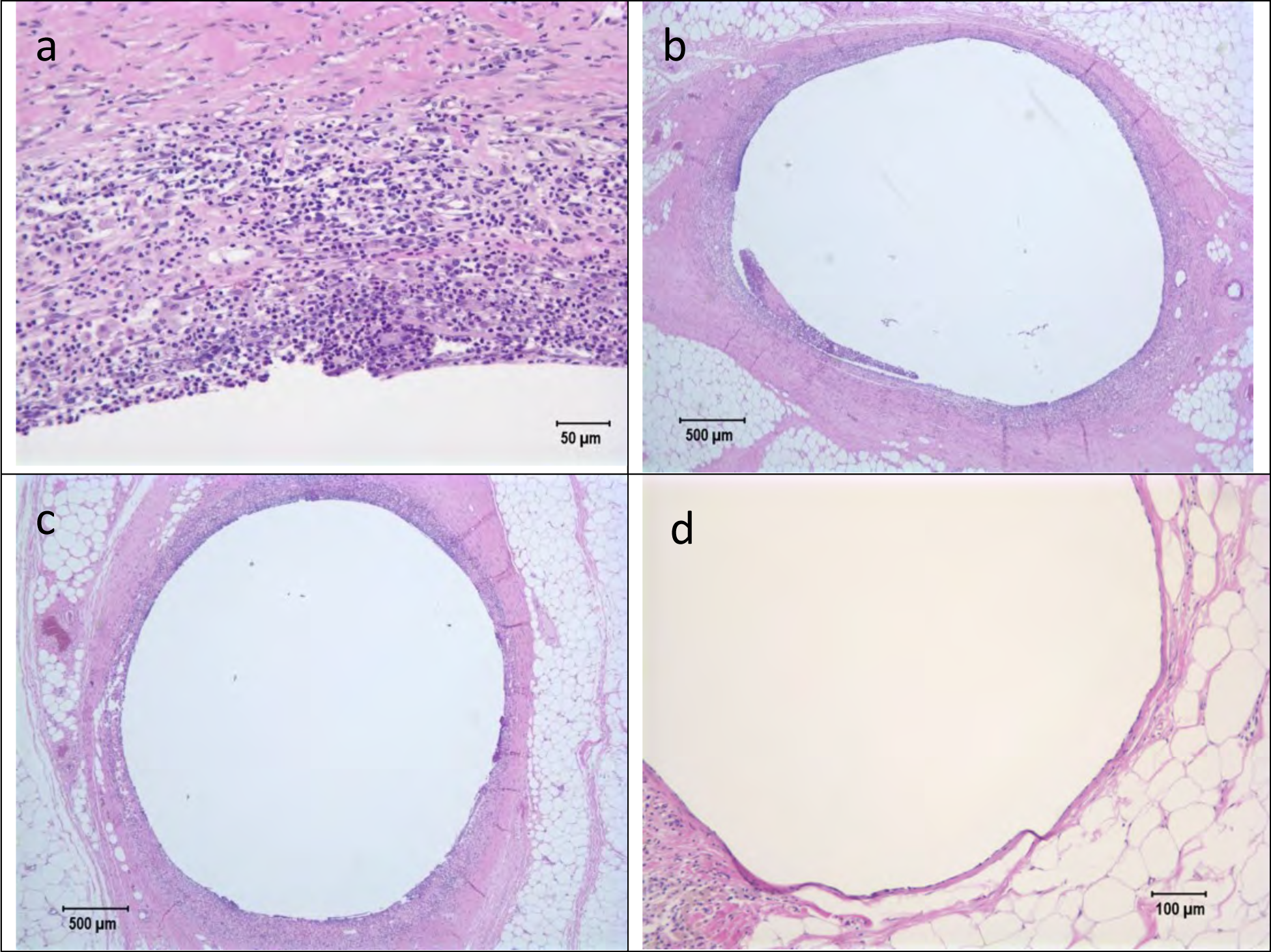
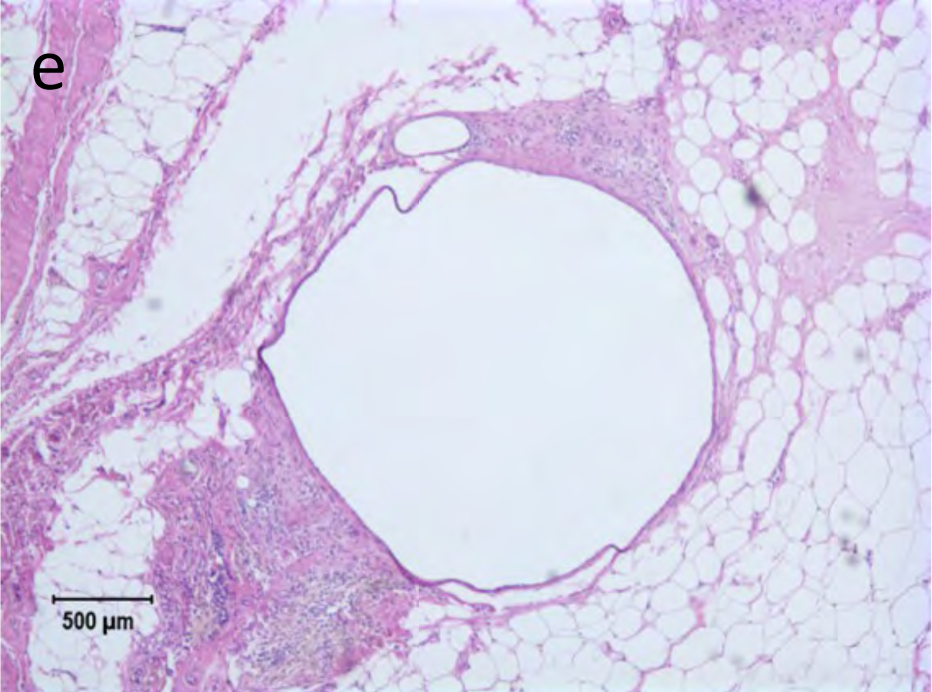
Rhesus macaque GH84. *In vitro* release rate 0.13 mg/day. The left (active) implant (a-c) shows a moderately thick band of heterophils, macrophages, lymphocytes, and plasma cells surrounding implant. Fibrous tissue and inflammation extend into adjacent connective tissue. Right (placebo) implant (d-e)has less inflammation, yet adjacent connective tissues of both implants have some fibrosis and minimal inflammation.

**12-week histology reports:**

**Table S10.** Histological characteristic scores for rhesus macaque FC48. An implant placed 2/22/2018; euthanized 5/17/2018.

| Animal# FC48<br>Necropsy# 18A219 | Response |  |
| --- | --- | --- |
| Implant | Left (Active) (1-4) | Right (Placebo) (1-4) |
| Polymorphonuclear cells | 4 | 2 |
| Lymphocytes | 4 | 3 |
| Plasma cells | 4 | 2 |
| Macrophages | 3 | 3 |
| Giant cells | 0 | 0 |
| Necrosis | 2 | 1 |
| Capsule thickness | 3 | 2 |
| Tissue infiltrate | 4 | 3 |
| Other |  |  |
| **Overall total** | **24** | **16** |

**Figure S27.**
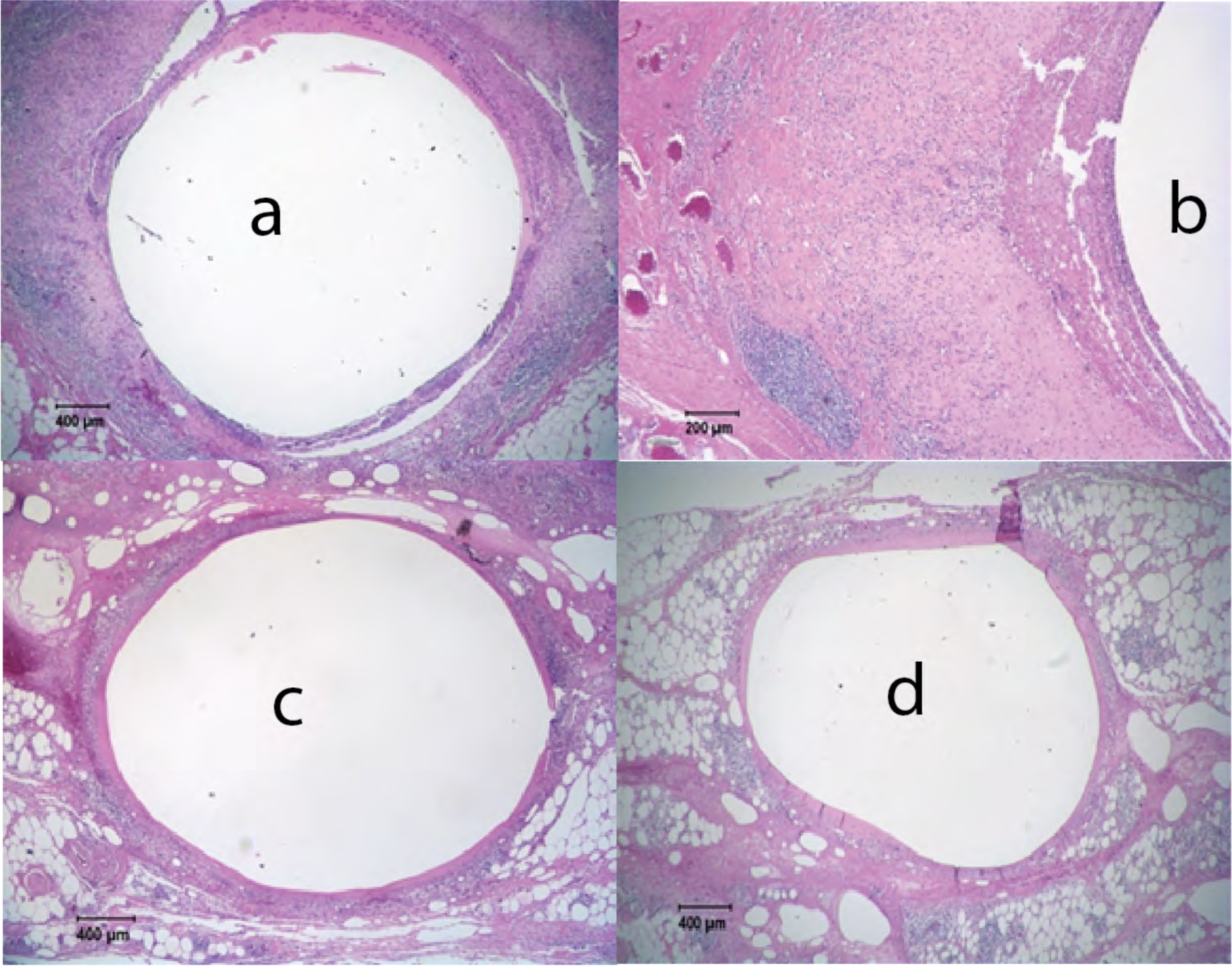
Rhesus macaque FC48. *In vitro* release rate 0.13 mg/day. Right implant (placebo) (c,d) has a thin capsule, and mild pericapsular infiltrates of lymphocytes and plasma cells, but deeper CT has multifocal aggregates of lymphocytes, plasma cells, edema, and hemorrhage. Left (active) implant (a,b) has thick capsule filled with proteinaceous fluid, heterophils, plasma cells, macrophages. Extensive fibrosis and lymphoplasmacytic inflammation extend into adjacent tissues

**Table S11.** Histological characteristic scores for rhesus macaque EC74. Implant placed 2/22/2018; euthanized 5/17/2018.

| Animal# EC74<br>Necropsy# 18A220 | Response |  |
| --- | --- | --- |
|  | *Left (Active) (1-4) | Right (Placebo) (1-4) |
| Implant |  |  |
| Polymorphonuclear cells | 4 | 1 |
| Lymphocytes | 4 | 3 |
| Plasma cells | 4 | 2 |
| Macrophages | 4 | 4 |
| Giant cells | 4 | 3 |
| Necrosis | 4 | 1 |
| Capsule thickness | 3 | 3 |
| Tissue infiltrate | 4 | 3 |
| Other |  |  |
| **Overall total** | **31** | **20** |

**Figure S28.**
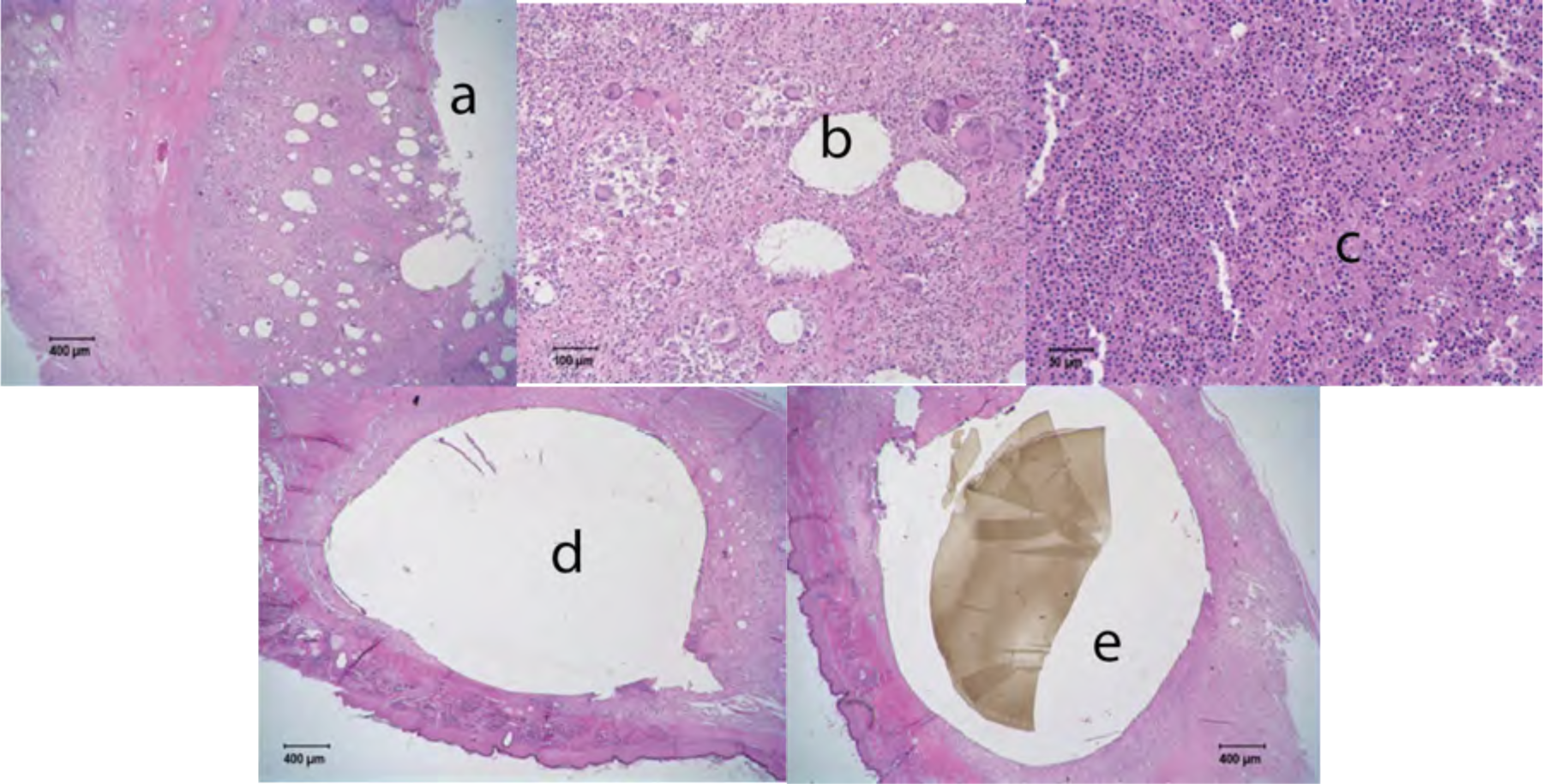
Rhesus macaque EC74. *In vitro* release rate 0.13 mg/day. *The left implant (active) (a, b, c) could not be located, but there was a purulent hemorrhagic abscess with fibrosis in the region and histology shows marked necrotic cellular material surrounded by severe granulomatous inflammation with abundant multinucleated giant cells. The left implant was lost through abscess formation before necropsy. The right implant (placebo) (d, e) has marked accumulations of macrophages and lymphocytes with moderate numbers of giant cells around the implant but minimal heterophils, plasma cells, or necrosis. Panel e shows a remnant of a sectioned piece of the membrane wall the occupies the implant volume after sectioning.

**Table S12.** Histological characteristic scores for rhesus macaque EE22. Implant placed 2/22/2018; euthanized 5/24/2018.

| Animal# EE22<br>Necropsy# 18A233 | Response |  |
| --- | --- | --- |
|  | Left (Active) (1-4) | Right (Placebo) (1-4) |
| Implant |  |  |
| Polymorphonuclear cells | 4 | 0 |
| Lymphocytes | 4 | 1 |
| Plasma cells | 4 | 0 |
| Macrophages | 4 | 0 |
| Giant cells | 0 | 0 |
| Necrosis | 3 | 0 |
| Capsule thickness | 3 | 1 |
| Tissue infiltrate | 4 | 0 |
| Other |  |  |
| **Overall total** | **26** | **2** |

**Figure S29.**
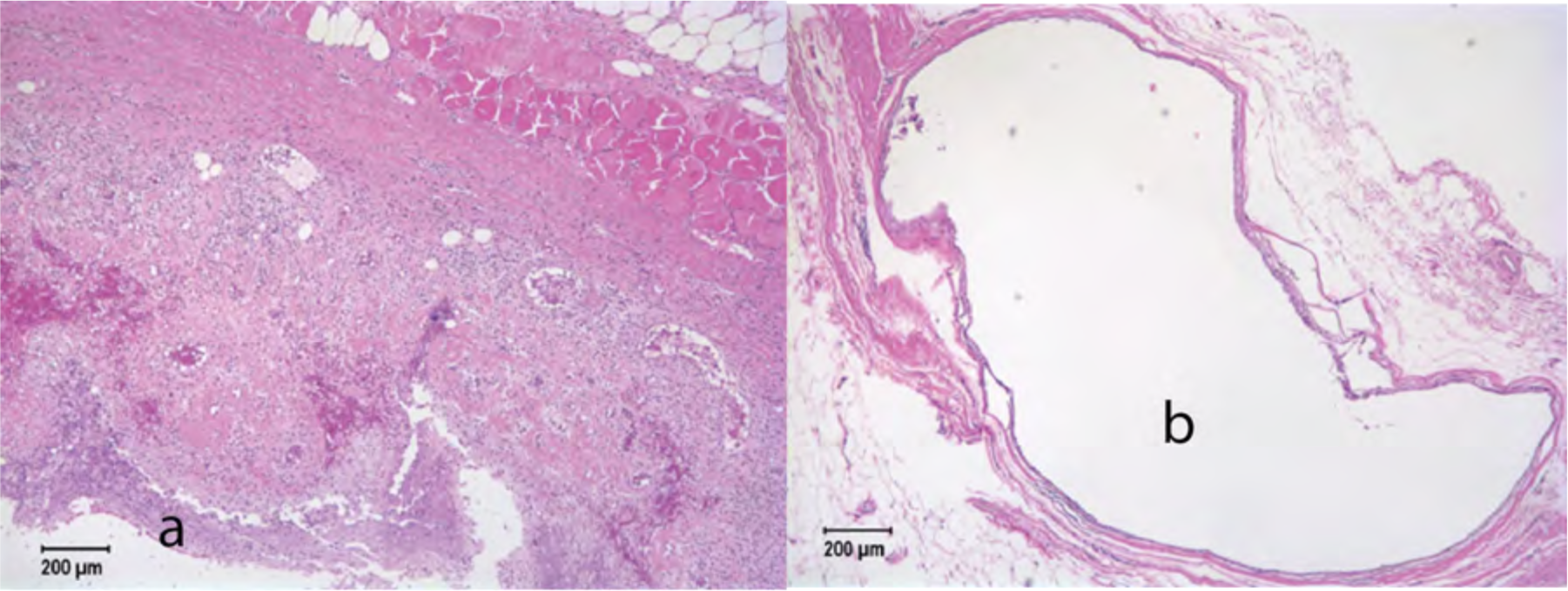
Rhesus macaque EE22. *In vitro* release rate 0.13 mg/day. Grossly there is an abscess containing serosanguinous watery fluid above the left scapular region. Histologically the left (active) implant (a) is surrounded by a thick capsule and thick layer of granulomatous inflammation with necrotic cellular material in the lumen. The right (placebo) implant (b) is surrounded by a thin capsule with minimal inflammation.

**Table S13.** Histological characteristic scores for rhesus macaque HC11. Implant placed 2/22/2018; euthanized 5/24/2018.

| **Animal# HC11**<br>**Necropsy# 18A234** | **Response** |  |
| --- | --- | --- |
|  | **Left (Active) (1-4)** | **Right (Placebo) (1-4)** |
| Implant |  |  |
| Polymorphonuclear cells | 4 | 1 |
| Lymphocytes | 4 | 2* |
| Plasma cells | 4 | 1 |
| Macrophages | 4 | 0 |
| Giant cells | 3 | 0 |
| Necrosis | 3 | 0 |
| Capsule thickness | 3 | 2 |
| Tissue infiltrate | 3 | 1* |
| Other |  |  |
| **Overall total** | **28** | **7=AV** |

**Figure S30.**
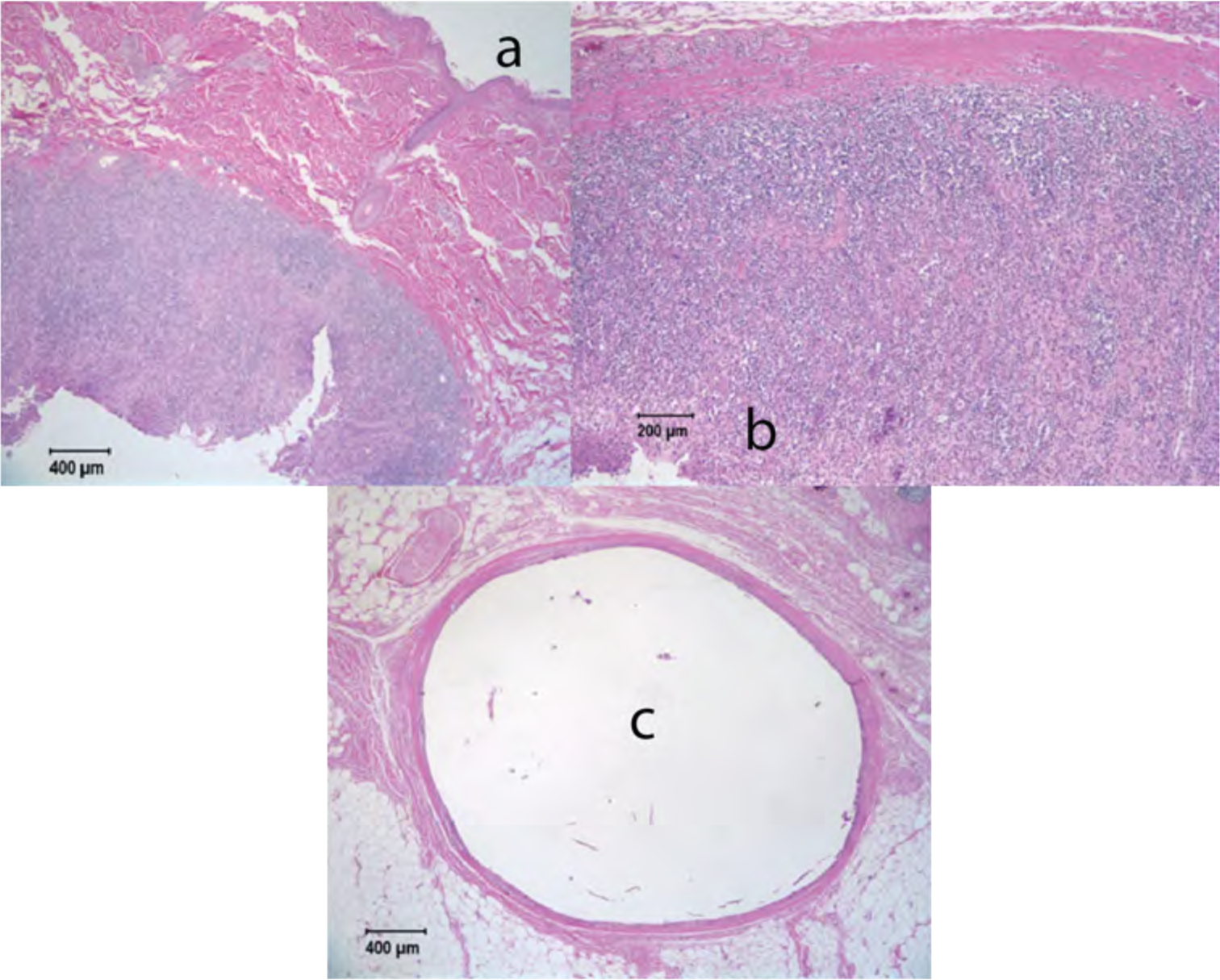
Rhesus macaque HC11. *In vitro* release rate 0.13 mg/day. Sections from left (active) implant (a,b) show thick fibrous capsule filled with heterophils, plasma cells, necrotic cellular debris, proteinaceous fluid, and occasional multinucleated giant cells. Little necrosis outside of capsule but multifocal aggregates of densely packed lymphocytes are in surrounding tissues. Right (placebo) implant (c) has minimal fibrosis or inflammation. A few heterophils and plasma cells in the lumen of the implant. Deeper tissues show mild infiltrate of lymphocytes and plasma cells but may be extensions from another implant.

## Supplemental 5. Implant reactivity grade summary

**Table S14.** Implant reactivity scores and placebo-adjusted implant reactivity scores.

| Animal | Implant Type | Time (week) | Number of implants tested | Average implant reactivity score (0-56) | Average placebo-adjusted implant reactivity score ($\bar{S}_{pair}$)<sup>a</sup> | Reactivity Grade |
| --- | --- | --- | --- | --- | --- | --- |
| Rabbit | Placebo Gen A<br>0.8 cm | 12 | 9 | 6 |  |  |
| Rabbit | Active Gen A<br>0.8 cm | 12 | 8 | 37 | 31 | Severe reaction |
| Rabbit | Active Gen A<br>1.6 cm | 12 | 8 | 40 | 34 | Severe reaction |
| Macaque | Placebo Gen B<br>2.0 cm | 4 | 2 | 11 |  |  |
| Macaque | Active Gen B<br>2.0 cm | 4 | 2 | 34 | 23 | Severe reaction |
| Macaque | Placebo Gen B<br>2.0 cm | 12 | 4 | 16 |  |  |
| Macaque | Active Gen B<br>2.0 cm | 12 | 4 | 48 | 32 | Severe reaction |

a. $\bar{S}_{pair}$ from 0.0 up to 2.9 no reaction, $\bar{S}_{pair}$ from 3.0 up to 8.9 a slight reaction, $\bar{S}_{pair}$ from 9.0 up to 15.0 a moderate reaction, and $\bar{S}_{pair} > 15.1$ is assigned severe reaction grade as per the standard (30).

## Supplemental 6. Molecular weight analysis of polymer wall material before and after in vivo exposure

Method: The core of the implants, recovered after the animal studies, was extracted for residual drug analysis. The tubing left over after the extraction procedure was processed to determine molecular weight. The tubing was dried in vacuum for 24 hours at room temperature. The dried tubing was dissolved in tetrahydrofuran (THF) and filtered through a 0.2 µm PTFE syringe filter. Gel permeation chromatography (GPC) was then used to measure the molecular weight of the tubing. A GPC system (Wyatt Technology, Santa Barbara, CA) consisting of a multi-angle light scattering detector (DAWN HELEOS-II 8 angle), and refractive index detector (Optilab T-rEX) was used. Software provided by the GPC instrument manufacturer (ASTRA software, Version 7.0) was used to process data. The GPC column used was a PLgel Mixed-B GPC column (10 µm, 7.5 x 300 mm) in combination with PLgel guard column (10 µm, 7.5 x 50 mm). The flow rate of the mobile phase, THF, was set at 0.7 mL/min. The dn/dc value of EG85A was 0.0756 mL/g.

The measured weight average molecular weights in TAF implant controls and implants from macaques for 12 weeks were 98.3±4.0 kDa and 95.9±2.9 kDa, respectively (Table S15. The measured weight average molecular weights of the TAF implant controls and implants from macaques for 12 weeks were 97.0±0.7 kDa and 91.3±2.5 kDa, respectively (Table S16).

**Table S15.** The number and weight average molecular weights (M_n_, M_w_) and polydispersity index values (M_w_/M_n_) of tubing recovered from Generation A implants in NZW rabbits after 12 weeks

| ID Number | $M_n$ (kDa) | $M_w$ (kDa) | Polydispersity ($M_w/M_n$) |
| --- | --- | --- | --- |
| TAF Implant Controls |  |  |  |
| Control 1 | 51.5 | 98.4 | 1.911 |
| Control 2 | 50.5 | 96.7 | 1.914 |
| Control 3 | 48.4 | 92.4 | 1.910 |
| Control 4 | 52.0 | 102.7 | 1.974 |
| Control 5 | 53.3 | 101.1 | 1.897 |
| Average (n=5) | 51.2±1.8 | 98.3±4.0 | 1.921±0.030 |
| TAF Implants in NZW Rabbits (12 weeks) |  |  |  |
| Implant 1 | 52.7 | 97.9 | 1.857 |
| Implant 2 | 48.7 | 91.2 | 1.875 |
| Implant 3 | 50.0 | 93.8 | 1.875 |
| Implant 4 | 50.6 | 95.2 | 1.880 |
| Implant 5 | 49.1 | 92.0 | 1.874 |
| Implant 6 | 50.2 | 92.4 | 1.839 |
| Implant 7 | 49.6 | 93.6 | 1.886 |
| Implant 8 | 51.4 | 95.2 | 1.853 |
| Implant 9 | 52.0 | 95.9 | 1.842 |
| Implant 10 | 50.4 | 98.2 | 1.949 |
| Implant 11 | 49.3 | 95.0 | 1.926 |
| Implant 12 | 52.5 | 100.0 | 1.904 |
| Implant 13 | 52.0 | 98.1 | 1.888 |
| Implant 14 | 52.9 | 100.8 | 1.904 |
| Implant 15 | 51.0 | 93.5 | 1.834 |
| Average (n=15) | 50.8±1.4 | 95.5±2.9 | 1.879±0.033 |

**Table S16.** The number and weight average molecular weights (M_n_, M_w_) and polydispersity index values (M_w_/M_n_) of tubing recovered from Generation A implants in rhesus macaques after 12 weeks

| ID Number | M<sub>n</sub> (kDa) | M<sub>w</sub> (kDa) | Polydispersity (M<sub>w</sub>/M<sub>n</sub>) |
| --- | --- | --- | --- |
| TAF Implant Controls |  |  |  |
| Control 1 | 48.3 | 96.8 | 2.003 |
| Control 2 | 47.2 | 97.0 | 2.054 |
| Control 3 | 46.1 | 97.8 | 2.120 |
| Control 4 | 47.8 | 96.2 | 2.015 |
| Average (n=4) | 47.4±0.9 | 97.0±0.7 | 2.048±0.053 |
| TAF Implants in Macaques (12 weeks) |  |  |  |
| Implant 1 | 44.7 | 92.4 | 2.069 |
| Implant 2 | 45.9 | 89.5 | 1.948 |
| Implant 3 | 41.9 | 92.3 | 2.203 |
| Implant 4 | 40.4 | 89.0 | 2.202 |
| Implant 5 | 45.5 | 91.0 | 2.002 |
| Implant 6 | 42.7 | 88.6 | 2.074 |
| Implant 7 | 46.1 | 91.5 | 1.986 |
| Implant 8 | 45.2 | 90.1 | 1.992 |
| Implant 9 | 48.5 | 96.9 | 1.998 |
| Average (n=9) | 44.5±2.5 | 91.3±2.5 | 2.053±0.094 |

## Supplemental 7. TFV-DP tissue and TAF & TFV plasma levels in rhesus macaques with generation B implants

**Table S17.** Gen B rhesus macaque TFV-DP tissue levels. TFV-DP tissue levels for BLQ for TFV-DP was <5 fmol/sample.

| Animal ID | Time (weeks) | Specimen Type | Final [TFV-DP] (fmol/mg) |
| --- | --- | --- | --- |
| EC74 | 12 | Implant | 26.8 |
| FC48 | 12 | Implant | 1.06 |
| EE22 | 13 | Implant | 9.13 |
| HC11 | 13 | Implant | BLQ |
| EJ87 | 0 | Rectal | BLQ |
| GR10 | 0 | Rectal | BLQ |
| EJ87 | 4 | Rectal | BLQ |
| GR10 | 4 | Rectal | BLQ |
| EC74 | 12 | Rectal | BLQ |
| EC74 | 12 | Rectal | BLQ |
| EE22 | 12 | Rectal | BLQ |
| FC48 | 12 | Rectal | 11.8 |
| FC48 | 12 | Rectal | 9.12 |
| HC11 | 12 | Rectal | BLQ |
| IT23 | 12 | Rectal | 2.18 |
| JC07 | 12 | Rectal | 2.10 |
| EE22 | 13 | Rectal | BLQ |
| EE22 | 13 | Rectal | BLQ |
| HC11 | 13 | Rectal | 1.21 |
| HC11 | 13 | Rectal | 1.40 |
| IT23 | 13 | Rectal | 1.80 |
| JC07 | 13 | Rectal | 7.20 |
| EJ87 | 0 | Vagina | BLQ |
| GR10 | 0 | Vagina | BLQ |
| EJ87 | 4 | Vagina | BLQ |
| GR10 | 4 | Vagina | BLQ |
| EC74 | 12 | Vagina | BLQ |
| EC74 | 12 | Vagina | BLQ |
| EE22 | 12 | Vagina | BLQ |
| FC48 | 12 | Vagina | BLQ |
| FC48 | 12 | Vagina | 0.994 |
| EE22 | 13 | Vagina | BLQ |
| EE22 | 13 | Vagina | BLQ |

**Table S18.** Gen B rhesus macaque TAF and TFV plasma levels.

| Specimen ID | Time Point | Final [TAF] (ng/mL) | Final [TFV] (ng/mL) |
| --- | --- | --- | --- |
| DP15 | Week 0 | BLQ | BLQ |
| DP15 | Week 1 | 0.050 | 0.358 |
| DP15 | Week 2 | 0.106 | 0.415 |
| DP15 | Week 3 | 0.111 | 0.487 |
| DP15 | Week 4 | 0.078 | 0.531 |
| EC74 | Week 0 | BLQ | BLQ |
| EC74 | Week 1 | 0.064 | 0.373 |
| EC74 | Week 10 | BLQ | 0.034 |
| EC74 | Week 12 | BLQ | BLQ |
| EC74 | Week 12 | BLQ | BLQ |
| EC74 | Week 2 | BLQ | 0.371 |
| EC74 | Week 3 | 0.035 | 0.621 |
| EC74 | Week 4 | 0.041 | 0.459 |
| EC74 | Week 5 | 0.803 | 0.07 |
| EC74 | Week 6 | 0.55 | 0.097 |
| EC74 | Week 7 | 0.674 | 0.059 |
| EC74 | Week 8 | 0.79 | 0.066 |
| EC74 | Week 9 | BLQ | 0.046 |
| EE22 | Week 0 | BLQ | BLQ |
| EE22 | Week 1 | 0.043 | 0.606 |
| EE22 | Week 10 | BLQ | BLQ |
| EE22 | Week 11 | BLQ | BLQ |
| EE22 | Week 12 | BLQ | BLQ |
| EE22 | Week 13 | BLQ | BLQ |
| EE22 | Week 13 | BLQ | BLQ |
| EE22 | Week 2 | 0.069 | 0.313 |
| EE22 | Week 3 | BLQ | 6.95 |
| EE22 | Week 4 | 0.048 | 0.696 |
| EE22 | Week 5 | 0.596 | 0.11 |
| EE22 | Week 6 | 0.529 | 0.109 |
| EE22 | Week 7 | 0.51 | 0.075 |
| EE22 | Week 8 | 0.624 | BLQ |
| EE22 | Week 9 | BLQ | BLQ |
| FC48 | Week 0 | BLQ | BLQ |
| FC48 | Week 1 | 0.040 | 1.05 |
| FC48 | Week 10 | 2.55 | 0.108 |
| FC48 | Week 12 | 0.819 | 0.124 |
| FC48 | Week 12 | 0.706 | 0.052 |
| FC48 | Week 2 | 0.053 | 0.941 |
| FC48 | Week 3 | 0.042 | 0.488 |
| FC48 | Week 4 | 0.062 | 0.560 |
| FC48 | Week 5 | 0.414 | 0.07 |
| FC48 | Week 6 | 0.56 | 0.115 |
| FC48 | Week 7 | 0.489 | 0.078 |
| FC48 | Week 8 | 0.726 | 0.101 |
| FC48 | Week 9 | 1.08 | 0.129 |
| GH84 | Week 0 | BLQ | BLQ |
| GH84 | Week 1 | 0.051 | BLQ |
| GH84 | Week 2 | 0.095 | BLQ |
| GH84 | Week 3 | 0.064 | BLQ |
| GH84 | Week 4 | 0.080 | 0.345 |
| HC11 | Week 0 | BLQ | BLQ |
| HC11 | Week 1 | 0.044 | BLQ |
| HC11 | Week 10 | BLQ | BLQ |
| HC11 | Week 11 | 0.663 | BLQ |
| HC11 | Week 12 | BLQ | BLQ |
| HC11 | Week 13 | BLQ | BLQ |
| HC11 | Week 13 | BLQ | BLQ |
| HC11 | Week 2 | BLQ | BLQ |
| HC11 | Week 3 | BLQ | BLQ |
| HC11 | Week 4 | BLQ | 0.578 |
| HC11 | Week 5 | BLQ | BLQ |
| HC11 | Week 6 | BLQ | BLQ |
| HC11 | Week 7 | 0.343 | BLQ |
| HC11 | Week 8 | 0.512 | BLQ |
| HC11 | Week 9 | 0.361 | BLQ |
| IT23 | Week 0 | BLQ | BLQ |
| IT23 | Week 1 | 0.045 | BLQ |
| IT23 | Week 10 | 0.491 | 0.045 |
| IT23 | Week 11 | BLQ | 0.038 |
| IT23 | Week 12 | 0.701 | 0.063 |
| IT23 | Week 13 | 0.73 | 0.048 |
| IT23 | Week 14 | 1.01 | 0.117 |
| IT23 | Week 2 | 0.031 | BLQ |
| IT23 | Week 3 | BLQ | 0.519 |
| IT23 | Week 4 | 0.038 | 0.485 |
| IT23 | Week 5 | 0.339 | BLQ |
| IT23 | Week 6 | 0.392 | 0.058 |
| IT23 | Week 7 | 0.437 | 0.054 |
| IT23 | Week 8 | 0.629 | 0.051 |
| IT23 | Week 9 | 3.66 | 0.038 |
| JC07 | Week 0 | BLQ | BLQ |
| JC07 | Week 1 | BLQ | BLQ |
| JC07 | Week 10 | 0.597 | 0.09 |
| JC07 | Week 11 | 0.444 | 0.059 |
| JC07 | Week 12 | BLQ | 0.057 |
| JC07 | Week 13 | BLQ | 0.074 |
| JC07 | Week 14 | 0.624 | 0.052 |
| JC07 | Week 2 | BLQ | BLQ |
| JC07 | Week 3 | BLQ | BLQ |
| JC07 | Week 4 | BLQ | 0.341 |
| JC07 | Week 5 | 0.481 | BLQ |
| JC07 | Week 6 | BLQ | BLQ |
| JC07 | Week 7 | 0.552 | BLQ |
| JC07 | Week 8 | 0.445 | BLQ |
| JC07 | Week 9 | 0.456 | 0.069 |

## Supplemental 8. Use of a trocar for implantation in a PK and safety study of the Generation B implant in rhesus macaques

**Figure S31.**
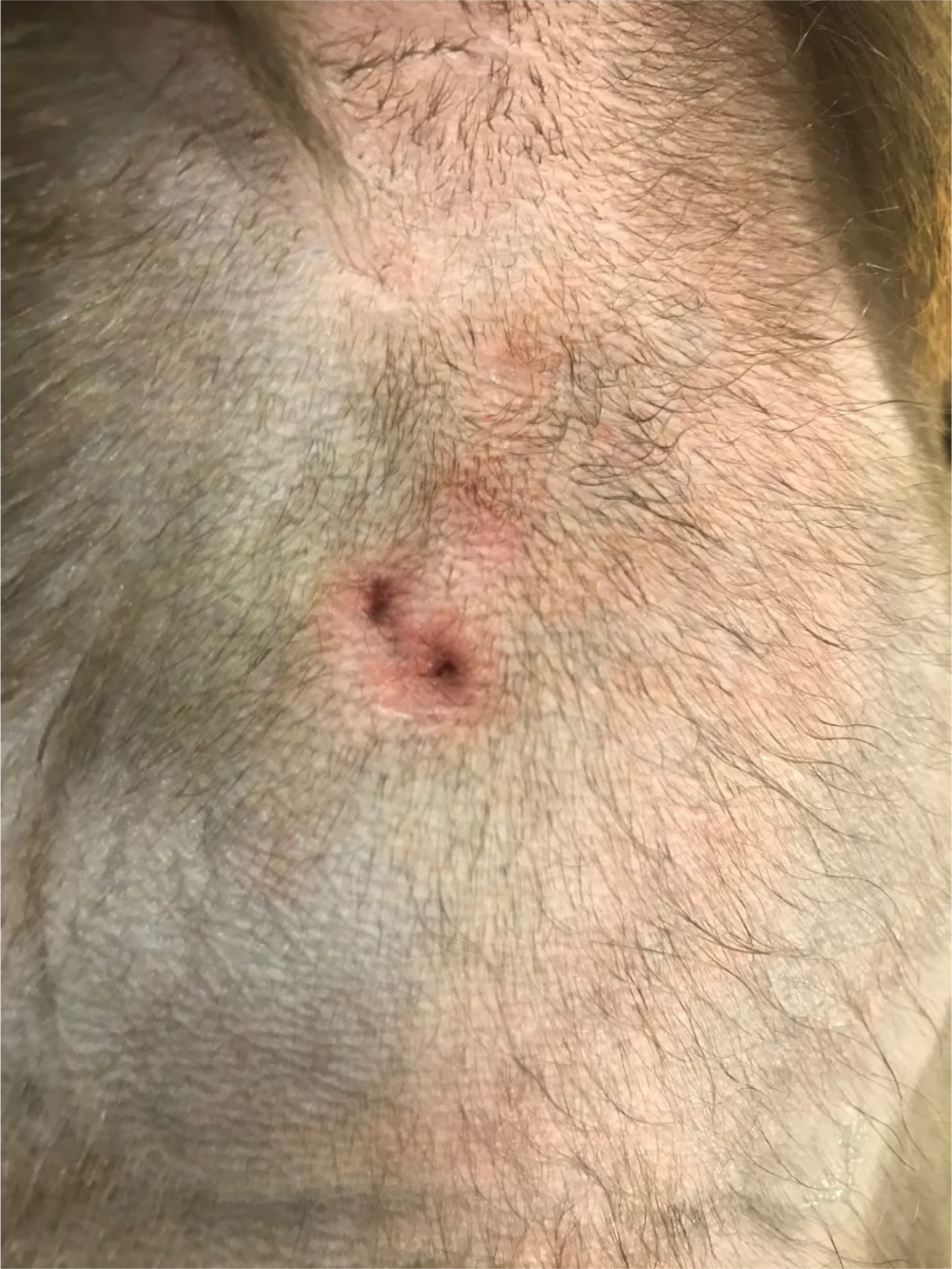
Trocar inserted implant resulted in a puncture like lesion. IT19 had 2 puncture-like lesions (with a small amount of bloody purulent discharge in the area of the left implant. The animal was started on antibiotics.

**Figure S32.**
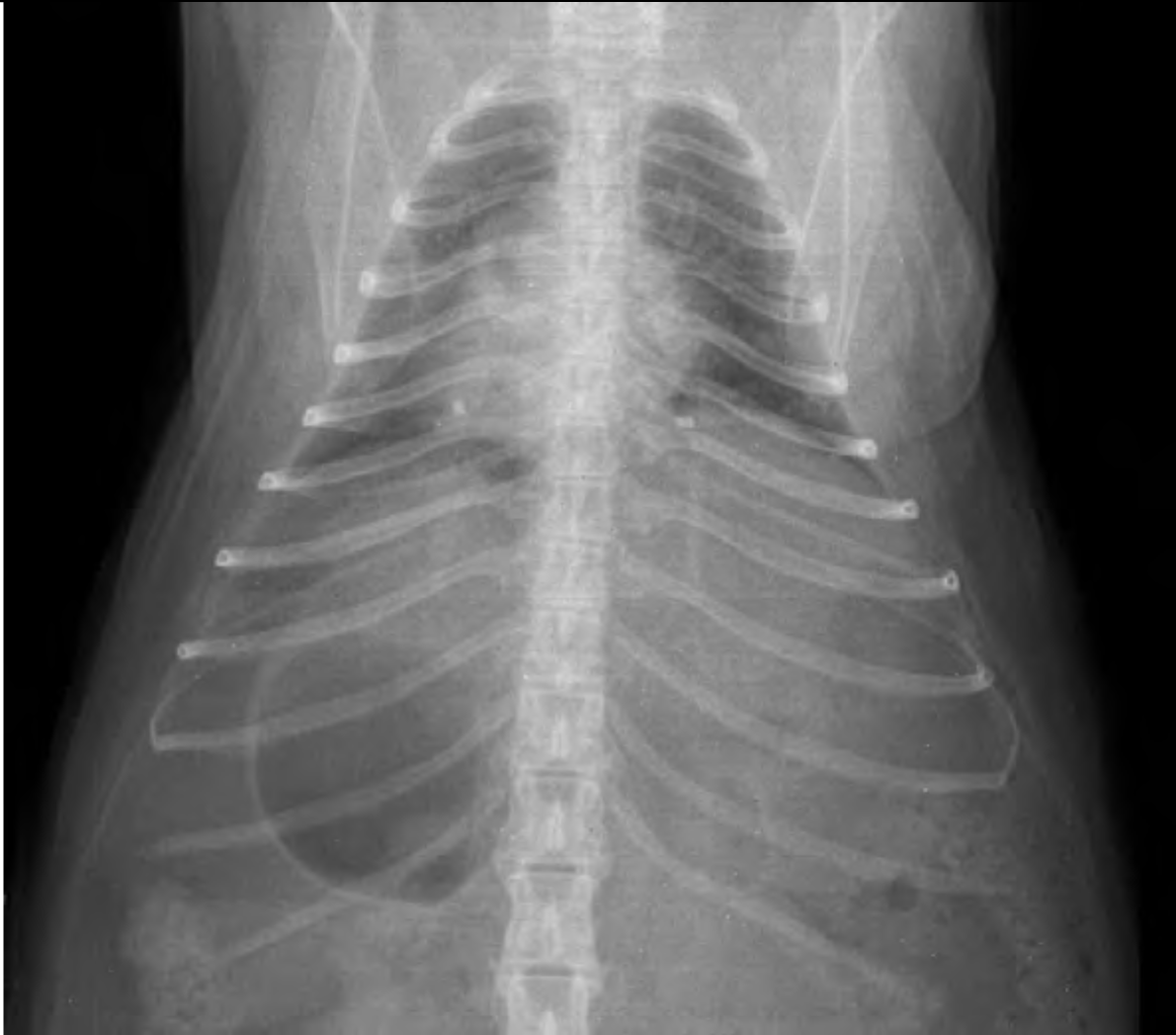
X-ray image of rhesus macaque with trocar inserted implants. An x-ray opaque barium sulfate pellet was added to each implant.

## REFERENCES

1. McGowan I. 2015. Injectable and implantable antiretroviral strategies for HIV prevention. Future Virology 10:1163–1176.

2. Baeten JM, Donnell D, Ndase P, Mugo NR, Campbell JD, Wangisi J, Tappero JW, Bukusi EA, Cohen CR, Katabira E, Ronald A, Tumwesigye E, Were E, Fife KH, Kiarie J, Farquhar C, John-Stewart G, Kakia A, Odoyo J, Mucunguzi A, Nakku-Joloba E, Twesigye R, Ngure K, Apaka C, Tamooh H, Gabona F, Mujugira A, Panteleeff D, Thomas KK, Kidoguchi L, Krows M, Revall J, Morrison S, Haugen H, Emmanuel-Ogier M, Ondrejcek L, Coombs RW, Frenkel L, Hendrix C, Bumpus NN, Bangsberg D, Haberer JE, Stevens WS, Lingappa JR, Celum C, Partners Pr EPST. 2012. Antiretroviral prophylaxis for HIV prevention in heterosexual men and women. N Engl J Med 367:399–410.

3. Grant RM, Lama JR, Anderson PL, McMahan V, Liu AY, Vargas L, Goicochea P, Casapia M, Guanira-Carranza JV, Ramirez-Cardich ME, Montoya-Herrera O, Fernandez T, Veloso VG, Buchbinder SP, Chariyalertsak S, Schechter M, Bekker LG, Mayer KH, Kallas EG, Amico KR, Mulligan K, Bushman LR, Hance RJ, Ganoza C, Defechereux P, Postle B, Wang F, McConnell JJ, Zheng JH, Lee J, Rooney JF, Jaffe HS, Martinez AI, Burns DN, Glidden DV, iPrEx Study T. 2010. Preexposure chemoprophylaxis for HIV prevention in men who have sex with men. N Engl J Med 363:2587–99.

4. Choopanya K, Martin M, Suntharasamai P, Sangkum U, Mock PA, Leethochawalit M, Chiamwongpaet S, Kitisin P, Natrujirote P, Kittimunkong S, Chuachoowong R, Gvetadze RJ, McNicholl JM, Paxton LA, Curlin ME, Hendrix CW, Vanichseni S, Bangkok Tenofovir Study G. 2013. Antiretroviral prophylaxis for HIV infection in injecting drug users in Bangkok, Thailand (the Bangkok Tenofovir Study): a randomised, double-blind, placebo-controlled phase 3 trial. Lancet 381:2083–90.

5. Thigpen MC, Kebaabetswe PM, Paxton LA, Smith DK, Rose CE, Segolodi TM, Henderson FL, Pathak SR, Soud FA, Chillag KL, Mutanhaurwa R, Chirwa LI, Kasonde M, Abebe D, Buliva E, Gvetadze RJ, Johnson S, Sukalac T, Thomas VT, Hart C, Johnson JA, Malotte CK, Hendrix CW, Brooks JT, Group TDFS. 2012. Antiretroviral preexposure prophylaxis for heterosexual HIV transmission in Botswana. N Engl J Med 367:423–34.

6. Marrazzo JM, Ramjee G, Richardson BA, Gomez K, Mgodi N, Nair G, Palanee T, Nakabiito C, van der Straten A, Noguchi L, Hendrix CW, Dai JY, Ganesh S, Mkhize B, Taljaard M, Parikh UM, Piper J, Masse B, Grossman C, Rooney J, Schwartz JL, Watts H, Marzinke MA, Hillier SL, McGowan IM, Chirenje ZM, Team VS. 2015. Tenofovir-based preexposure prophylaxis for HIV infection among African women. N Engl J Med 372:509–18.

7. Haberer JE. 2016. Current concepts for PrEP adherence in the PrEP revolution: from clinical trials to routine practice. Curr Opin HIV AIDS 11:10–7.

8. van der Straten A, Brown ER, Marrazzo JM, Chirenje MZ, Liu K, Gomez K, Marzinke MA, Piper JM, Hendrix CW, Network M-VPTfMT. 2016. Divergent adherence estimates with pharmacokinetic and behavioural measures in the MTN-003 (VOICE) study. J Int AIDS Soc 19:20642.

9. Matthews R. 2019. First-in-Human Trial of MK-8591-Eluting Implants Demonstrates Concentrations Suitable for HIV Propylaxis for at Least One Year, 10th IAS Conference on HIV Science, Mexico City.

10. Sundaram A, Vaughan B, Kost K, Bankole A, Finer L, Singh S, Trussell J. 2017. Contraceptive Failure in the United States: Estimates from the 2006-2010 National Survey of Family Growth. Perspect Sex Reprod Health 49:7–16.

11. Trussell J. 2011. Contraceptive failure in the United States. Contraception 83:397–404.

12. Trussell J, Kost K. 1987. Contraceptive failure in the United States: a critical review of the literature. Stud Fam Plann 18:237–83.

13. Trussell J, Henry N, Hassan F, Prezioso A, Law A, Filonenko A. 2013. Burden of unintended pregnancy in the United States: potential savings with increased use of long-acting reversible contraception. Contraception 87:154–61.

14. Ray AS, Fordyce MW, Hitchcock MJ. 2016. Tenofovir alafenamide: A novel prodrug of tenofovir for the treatment of Human Immunodeficiency Virus. Antiviral Res 125:63–70.

15. Bam RA, Birkus G, Babusis D, Cihlar T, Yant SR. 2014. Metabolism and antiretroviral activity of tenofovir alafenamide in CD4+ T-cells and macrophages from demographically diverse donors. Antivir Ther 19:669–77.

16. Lee WA, He GX, Eisenberg E, Cihlar T, Swaminathan S, Mulato A, Cundy KC. 2005. Selective intracellular activation of a novel prodrug of the human immunodeficiency virus reverse transcriptase inhibitor tenofovir leads to preferential distribution and accumulation in lymphatic tissue. Antimicrob Agents Chemother 49:1898–906.

17. Hawkins T, Veikley W, St Claire RL, Guyer B, Clark N, Kearney BP. 2005. Intracellular pharmacokinetics of tenofovir diphosphate, carbovir triphosphate, and lamivudine triphosphate in patients receiving triple-nucleoside regimens. Jaids-Journal of Acquired Immune Deficiency Syndromes 39:406–411.

18. Massud I, Mitchell J, Babusis D, Deyounks F, Ray AS, Rooney JF, Heneine W, Miller MD, Garcia-Lerma JG. 2016. Chemoprophylaxis With Oral Emtricitabine and Tenofovir Alafenamide Combination Protects Macaques From Rectal Simian/Human Immunodeficiency Virus Infection. J Infect Dis 214:1058–62.

19. Hare CB, Coll J, Ruane P, Molina J-M, Mayer KH, Jessen H, Grant RM, Wet JJD, Thompson M, DeJesus E, Ebrahimi R, Giler RM, Das M, Brainard D, McCallister S. THE PHASE 3 DISCOVER STUDY: DAILY F/TAF OR F/TDF FOR HIV PREEXPOSURE PROPHYLAXIS, p. *In* (ed),

20. Gunawardana M, Remedios-Chan M, Miller CS, Fanter R, Yang F, Marzinke MA, Hendrix CW, Beliveau M, Moss JA, Smith TJ, Baum MM. 2015. Pharmacokinetics of long-acting tenofovir alafenamide (GS-7340) subdermal implant for HIV prophylaxis. Antimicrob Agents Chemother 59:3913–9.

21. Schlesinger E, Johengen D, Luecke E, Rothrock G, McGowan I, van der Straten A, Desai T. 2016. A Tunable, Biodegradable, Thin-Film Polymer Device as a Long-Acting Implant Delivering Tenofovir Alafenamide Fumarate for HIV Pre-exposure Prophylaxis. Pharm Res 33:1649–56.

22. Johnson LM, Krovi SA, Li L, Girouard N, Demkovich ZR, Myers D, Creelman B, van der Straten A. 2019. Characterization of a Reservoir-Style Implant for Sustained Release of Tenofovir Alafenamide (TAF) for HIV Pre-Exposure Prophylaxis (PrEP). Pharmaceutics 11.

23. Chua CYX, Jain P, Ballerini A, Bruno G, Hood RL, Gupte M, Gao S, Di Trani N, Susnjar A, Shelton K, Bushman LR, Folci M, Filgueira CS, Marzinke MA, Anderson PL, Hu M, Nehete P, Arduino RC, Sastry JK, Grattoni A. 2018. Transcutaneously refillable nanofluidic implant achieves sustained level of tenofovir diphosphate for HIV pre-exposure prophylaxis. J Control Release 286:315–325.

24. Gatto G, Girouard N, Brand RM, Johnson L, Marzinke M, Rowshan S, Engstrom JC, McGowan I, Demkovich Z, Lueke E, van der Straten A. 2019. Pharmacokinetics of tenofovir alafenamide by subcutaneous implant for HIV PREP., abstr Conference on Retroviruses and Opportunistic Infections, Seattle, WA, March 4–7.

25. Anderson JM, Rodriguez A, Chang DT. 2008. Foreign body reaction to biomaterials. Semin Immunol 20:86–100.

26. Greco RS. 1994. Implantation biology: the host response and biomedical devices. CRC Press, Boca Raton.

27. Golla VM, Kurmi M, Shaik K, Singh S. 2016. Stability behaviour of antiretroviral drugs and their combinations. 4: Characterization of degradation products of tenofovir alafenamide fumarate and comparison of its degradation and stability behaviour with tenofovir disoproxil fumarate. J Pharm Biomed Anal 131:146–155.

28. Anderson PL, Glidden DV, Liu A, Buchbinder S, Lama JR, Guanira JV, McMahan V, Bushman LR, Casapia M, Montoya-Herrera O, Veloso VG, Mayer KH, Chariyalertsak S, Schechter M, Bekker LG, Kallas EG, Grant RM, iPrEx Study T. 2012. Emtricitabine-tenofovir concentrations and pre-exposure prophylaxis efficacy in men who have sex with men. Sci Transl Med 4:151ra125.

29. Hendrix CW, Andrade A, Bumpus NN, Kashuba AD, Marzinke MA, Moore A, Anderson PL, Bushman LR, Fuchs EJ, Wiggins I, Radebaugh C, Prince HA, Bakshi RP, Wang R, Richardson P, Shieh E, McKinstry L, Li X, Donnell D, Elharrar V, Mayer KH, Patterson KB. 2016. Dose Frequency Ranging Pharmacokinetic Study of Tenofovir-Emtricitabine After Directly Observed Dosing in Healthy Volunteers to Establish Adherence Benchmarks (HPTN 066). AIDS Res Hum Retroviruses 32:32–43.

30. Food and Drug Administration: Center for Devices and Radiological Health. 2016. Use of International Standard ISO 10993-1, “Biological evaluation of medical devices - Part 1: Evaluation and testing within a risk management process”. US Department of Health and Human Services, Federal Register. https://www.govinfo.gov/content/pkg/FR-2016-06-16/pdf/2016-14190.pdf.

31. Gad SC, Gad-McDonald S. 2016. Biocompatibility Testing and Safety Assessment. CRC Press, Boca Raton, FL.

32. International Organization for Standardization. 2016. Biological Evaluation of Medical Devices 3rd Edition. International Organization for Standardization.

33. Anderson JM. 2015. Exploiting the inflammatory response on biomaterials research and development. J Mater Sci Mater Med 26:121.

34. Karim SSA, Gengiah TN, Karim QA. 2018. CAPRISA 018: A Phase I/II trial to assess the safety, acceptability, tolerability and pharmacokinetics of a sustained-release tenofovir alafenamide sub-dermal implant for HIV prevention in women. Centre for the AIDS Programme of Research in South Africa (CAPRISA), University of KwaZulu-Natal, Durban, South Africa.

35. Abdool Karim Q, Abdool Karim SS, Frohlich JA, Grobler AC, Baxter C, Mansoor LE, Kharsany AB, Sibeko S, Mlisana KP, Omar Z, Gengiah TN, Maarschalk S, Arulappan N, Mlotshwa M, Morris L, Taylor D, Group CT. 2010. Effectiveness and safety of tenofovir gel, an antiretroviral microbicide, for the prevention of HIV infection in women. Science 329:1168–74.

36. Delany-Moretlwe S, Lombard C, Baron D, Bekker LG, Nkala B, Ahmed K, Sebe M, Brumskine W, Nchabeleng M, Palanee-Philips T, Ntshangase J, Sibiya S, Smith E, Panchia R, Myer L, Schwartz JL, Marzinke M, Morris L, Brown ER, Doncel GF, Gray G, Rees H. 2018. Tenofovir 1% vaginal gel for prevention of HIV-1 infection in women in South Africa (FACTS-001): a phase 3, randomised, double-blind, placebo-controlled trial. Lancet Infect Dis 18:1241–1250.

37. Clark JT, Clark MR, Shelke NB, Johnson TJ, Smith EM, Andreasen AK, Nebeker JS, Fabian J, Friend DR, Kiser PF. 2014. Engineering a segmented dual-reservoir polyurethane intravaginal ring for simultaneous prevention of HIV transmission and unwanted pregnancy. PLoS One 9:e88509.

38. Anderson JM. 1994. In vivo Biocompatibility of Implantable Delivery Systems and Biomaterials. Eur J Pharm BioPharm 40:1–8.

39. Yamaguchi K, Anderson JM. 1992. Biocompatibility studies of naltrexone sustained release formulations. Journal of Controlled Release 19:299–314.

40. Lai CL, Shouval D, Lok AS, Chang TT, Cheinquer H, Goodman Z, DeHertogh D, Wilber R, Zink RC, Cross A, Colonno R, Fernandes L, Group BEAS. 2006. Entecavir versus lamivudine for patients with HBeAg-negative chronic hepatitis B. N Engl J Med 354:1011–20.

41. Henry SJ, Barrett SE, Forster SP, Teller RS, Yang Z, Li L, Mackey MA, Doto GJ, Ruth MP, Tsuchiya T, Klein LJ, Gindy ME. 2019. Exploration of long-acting implant formulations of hepatitis B drug entecavir. Eur J Pharm Sci 136:104958.

42. McGowan I, Hoesley C, Cranston RD, Andrew P, Janocko L, Dai JY, Carballo-Dieguez A, Ayudhya RK, Piper J, Hladik F, Mayer K. 2013. A phase 1 randomized, double blind, placebo controlled rectal safety and acceptability study of tenofovir 1% gel (MTN-007). PLoS One 8:e60147.

43. Rodriguez-Garcia M, Patel MV, Shen Z, Bodwell J, Rossoll RM, Wira CR. 2017. Tenofovir Inhibits Wound Healing of Epithelial Cells and Fibroblasts from the Upper and Lower Human Female Reproductive Tract. Sci Rep 8:45725.

44. Keller MJ, Ray L, Atrio J, Espinoza L, Sinclair S, Goymer J, Marzinke MA, Frank B, Anderson PL, Marrazzo JM, Mugo N, Hendrix C, Herold BC. Early Termination of a Phase 1 Trial of Tenofovir Disoproxil Fumarate Vaginal Ring, p. *In* (ed),

45. Mehellou Y. 2016. The ProTides Boom. ChemMedChem 11:1114–6.

46. Salie ZL, Kirby KA, Michailidis E, Marchand B, Singh K, Rohan LC, Kodama EN, Mitsuya H, Parniak MA, Sarafianos SG. 2016. Structural basis of HIV inhibition by translocation-defective RT inhibitor 4’-ethynyl-2-fluoro-2’-deoxyadenosine (EFdA). Proc Natl Acad Sci U S A 113:9274–9.

47. Michailidis E, Huber AD, Ryan EM, Ong YT, Leslie MD, Matzek KB, Singh K, Marchand B, Hagedorn AN, Kirby KA, Rohan LC, Kodama EN, Mitsuya H, Parniak MA, Sarafianos SG. 2014. 4’-Ethynyl-2-fluoro-2’-deoxyadenosine (EFdA) inhibits HIV-1 reverse transcriptase with multiple mechanisms. J Biol Chem 289:24533–48.

48. Michailidis E, Ryan EM, Hachiya A, Kirby KA, Marchand B, Leslie MD, Huber AD, Ong YT, Jackson JC, Singh K, Kodama EN, Mitsuya H, Parniak MA, Sarafianos SG. 2013. Hypersusceptibility mechanism of Tenofovir-resistant HIV to EFdA. Retrovirology 10:65.

49. Sager JE, Begley R, Rhee M, West SK, Ling J, Schroeder SD, Tse WC, Mathias A. 2019. Safety and PK of Subcutaneous GS-6207, A Novel HIV-1 Capsid Inhibitor, abstr Conference on Retroviruses and Opportunistic Infections, Seattle, Washington, March 4–7.

50. Markowitz M, Gettie A, St Bernard L, Andrews CD, Mohri H, Horowitz A, Grasperge BF, Blanchard JL, Niu T, Sun L, Fillgrove K, Hazuda DJ, Grobler JA. 2019. Title: Once-weekly Oral Dosing of MK-8591 Protects Male Rhesus Macaques from Intrarectal SHIV109CP3 Challenge. J Infect Dis doi:10.1093/infdis/jiz271.

51. Matthews R. First-in-Human Trial of MK-8591-Eluting Implants Demonstrates Concentrations Suitable for HIV Propylaxis for at Least One Year, p. *In* (ed),

52. Barrett SE, Teller RS, Forster SP, Li L, Mackey MA, Skomski D, Yang Z, Fillgrove KL, Doto GJ, Wood SL, Lebron J, Grobler JA, Sanchez RI, Liu Z, Lu B, Niu T, Sun L, Gindy ME. 2018. Extended-Duration MK-8591-Eluting Implant as a Candidate for HIV Treatment and Prevention. Antimicrob Agents Chemother 62(10).

53. Hummert P, Parsons TL, Ensign LM, Hoang T, Marzinke MA. 2018. Validation and implementation of liquid chromatographic-mass spectrometric (LC-MS) methods for the quantification of tenofovir prodrugs. J Pharm Biomed Anal 152:248–256.

54. King T, Bushman L, Kiser J, Anderson PL, Ray M, Delahunty T, Fletcher CV. 2006. Liquid chromatography-tandem mass spectrometric determination of tenofovir-diphosphate in human peripheral blood mononuclear cells. J Chromatogr B Analyt Technol Biomed Life Sci 843:147–56.

55. Food and Drug Adminsitration: Center for Drug Evalutaion and Research. 2018. Bioanalytical Method Validation: Guidance for Industry. Department of Health and Human Services, Federal Register, https://www.govinfo.gov/content/pkg/FR-2018-05-22/pdf/2018-10926.pdf.

